# Generation of dynamic three-dimensional genome structure through phase separation of chromatin

**DOI:** 10.1101/2021.05.06.443035

**Authors:** Shin Fujishiro, Masaki Sasai

**Affiliations:** Department of Applied Physics, Nagoya University, Nagoya 464-8601, Japan

**Author notes:** **Author contributions:** SF and MS designed research and wrote the manuscript. SF developed methods, performed research, and analyzed the data.

**Keywords:** genome organization, A/B compartments, chromatin domains, lamina-associated domains, nucleolous-associated domains

## Abstract

Three-dimensional genome structure and dynamics play critical roles in regulating DNA functions. Flexible chromatin structure and movements suggested that the genome is dynamically phase-separated to form A (active) and B (inactive) compartments in interphase nuclei. Here, we examined this hypothesis by developing a polymer model of the whole genome of human cells and assessing the impact of phase separation on genome structure. Upon entry to the G1 phase, the simulated genome expanded according to heterogeneous repulsion among chromatin chains, which moved chromatin heterogeneously, inducing phase separation of chromatin. This repulsion-driven phase separation quantitatively reproduces the experimentally observed chromatin domains, A/B compartments, lamina-associated domains, and nucleolus-associated domains, consistently explaining nuclei of different human cells and predicting their dynamic fluctuations. We propose that phase separation induced by heterogeneous repulsive interactions among chromatin chains largely determines dynamic genome organization.

**Significance:** DNA functions in living cells are crucially affected by the three-dimensional genome structure and dynamics. We analyzed the whole genome of human cells by developing a polymer model of interphase nuclei. The model revealed the essential importance of the unfolding process of chromosomes from the condensed mitotic state for describing the interphase nuclei; through the unfolding process, heterogeneous repulsive interactions among chromatin chains induced phase separation of chromatin, which quantitatively explains the experimentally observed various genomic data. We can use this model structure as a platform to analyze the relationship among genome structure, dynamics, and functions.

---

Three-dimensional genome structure and its dynamics play crucial roles in regulating eukaryotic DNA functions [1, 2, 3]. The recent development of techniques, such as high-throughput chromosome conformation capture (Hi-C and the related methods) [4, 5, 6], electron microscopy [7], and super-resolution microscopy [8, 9, 10, 11], has enhanced our understanding of genome organization. For further clarifying the mechanisms of genome organization, it is necessary to develop reliable computational models that can bridge these different experimental paradigms. Computational polymer models of individual chromosomes and their complexes were developed using the Hi-C data of global chromatin contacts as the input to deduce the knowledge-based forces on chromatin [12, 13, 14, 15]. More refined input data such as the global genome-wide contact pattern in the single-cell Hi-C data was necessary for modeling the whole-genome structure of mouse and human cells [16, 17, 18]. For further elucidating the principles of genome organization, it is highly desirable to develop a physical model of the whole genome using straightforward assumptions instead of fitting the model to the vast amount of experimental data on the global genome conformation.

In physical modeling of the whole genome, the important subject is to examine the possible phase separation of chromatin and assess its impact on genome structure. Chromatin shows flexible configuration [7, 8, 9] and movements [19, 10, 11], suggesting that chromatin in interphase nuclei is dynamically phase-separated to determine the genome structure [20, 21, 22, 23]. In the present study, we tested this hypothesis by examining the mechanism of phase separation. A previously proposed mechanism, which is still under debate [24], is the droplet-like condensation of factors such as HP1; these condensates may mediate attraction between heterochromatin regions, leading to phase separation of heterochromatin from euchromatin [25, 26]. Following this idea, the previous whole-genome models assumed attractive interactions, but these interactions spontaneously gathered heterochromatin toward the nuclear center, leading to the unusual genomic configuration. This anomalous chromatin distribution was remedied in the models by assuming the counteracting attractive interactions between chromatin and the nuclear lamina [27, 22]. However, the mechanism to establish such a balance among competitive interactions in the nucleus was unclear. In the present study, we resolved this difficulty by focusing on repulsion rather than attraction in chromatin interactions. We consider a polymer model, which describes heterogeneously distributed physical properties of chromatin. With heterogeneous repulsive interactions among chromatin regions, the simulated genome unfolded from the mitotic chromosomes, which generated heterogeneous movement of chromatin chains, leading to phase separation of chromatin in the G1 phase. This repulsion-driven phase separation quantitatively explains the genome organization of human fibroblast (IMR90) and lymphoblastoid (GM12878) cells and predicts dynamic fluctuations residing after the genome reached the G1 phase.

## Results

### Neighboring region contact index

The interactions between chromatin regions depend on the local physical properties of chromatin. We inferred these physical properties from the local chromatin contacts. Fig. 1A shows a distribution of the ratio of observed/expected contact frequencies obtained from the Hi-C data [5], 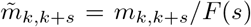, where *m*_*k,k*+*s*_ is the observed contact frequency between *k* and *k* + *s*th positions along the sequence, and *F* (*s*) is the mean contact frequency for the sequence separation *s*. Contacts between chromatin loci with the sequence separation *s* < 300 kb are more frequent in compartment A than compartment B, while contacts with *s* > 300 kb are more frequent in compartment B (Fig. 1A). Here, compartment A/B was identified by the principal component analysis (PCA) of the Hi-C contact matrix of the genome [4]. In other words, the properties of chromatin in a few hundred kb scale are correlated to the compartments defined in hundreds of Mb or the larger scale We quantified this correlation by defining the *neighboring region contact index* (NCI), 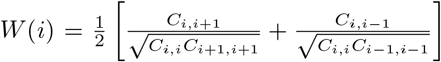, where *C*_*i,j*_ = Σ_*k*∈(*i*th region)_ Σ_*l*∈(*j*th region)_ *m*_*k,l*_ is a sum of the frequency of contacts between 50-kb chromatin regions labeled *i* and *j* along the sequence. As shown in Figs. 1B and 1C, NCI correlates to the compartment signal defined by PCA. This finding suggests that the A/B compartmentalization originates from the heterogeneity of local chromatin properties as captured by NCI. We examined this hypothesis by performing the whole-genome polymer simulation.

**Figure 1:**
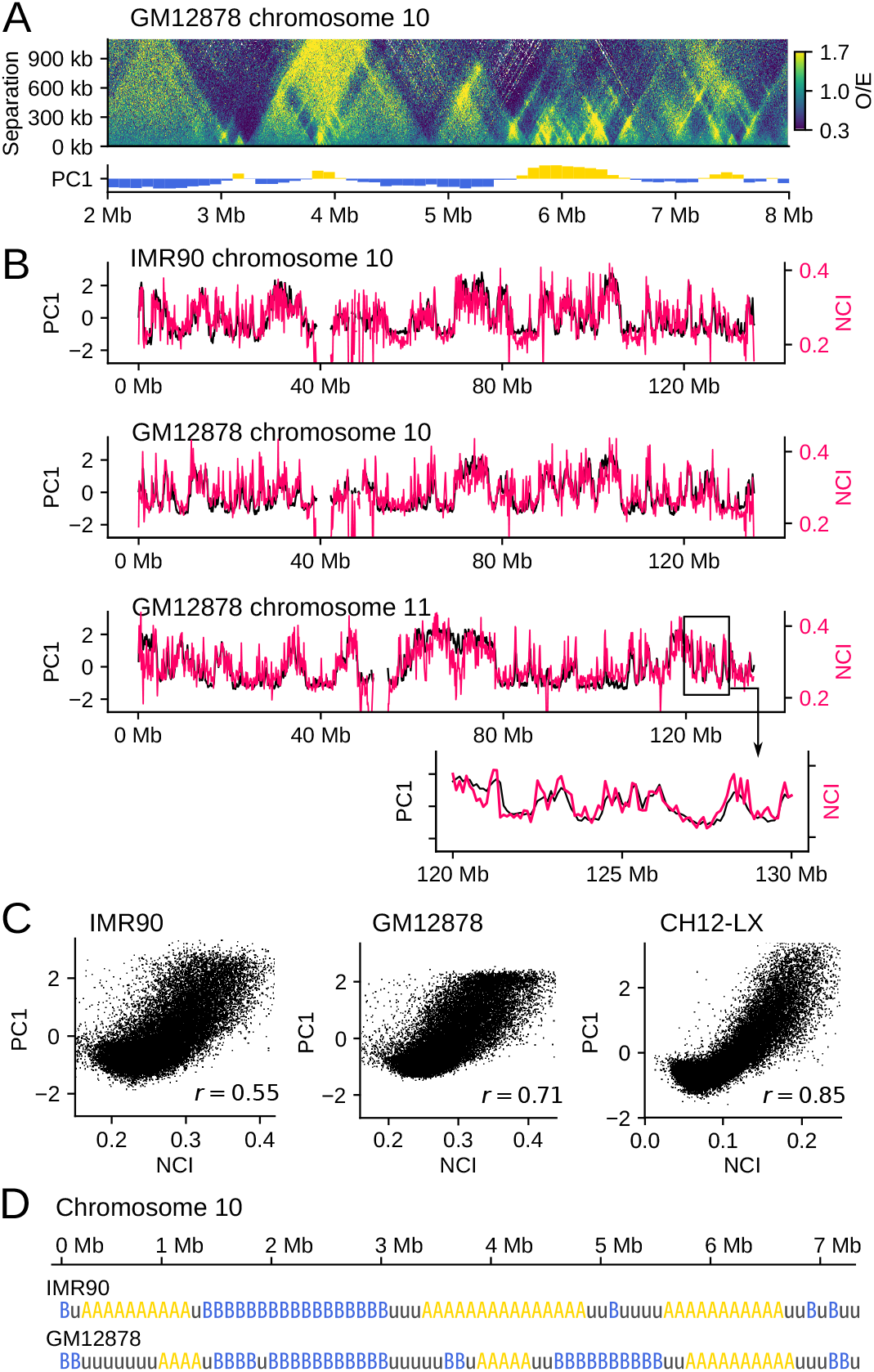
Correlation between the local contact frequency and the global compartment signal.(*A*) The ratio of observed/expected Hi-C contact frequencies 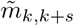 plotted on the plane of the position along the sequence *k* and the sequence separation *s* for chromosome 10 of GM12878 (top) is compared with the first principal component vector (PC1) of 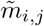 of the whole genome [5], which is the compartment signal [4] distinguishing compartments A and B (bottom). The compartment signal shows compartment A when it is positive (yellow) and compartment B when it is negative (blue). (*B*) The neighboring region contact index (NCI), *W* (*i*) (red), is superposed on the compartment signal PC1 (black) for chromosome 10 of IMR90 and GM12878 and chromosome 11 of GM12878. The inset is a close-up view of an example 10 Mb region. (*C*) Scattered plots for comparing the whole genome data of NCI and the compartment signal PC1 of IMR90, GM12878, and CH12-LX [5]. Each dot is a 100 kb segment. Pearson’s correlation coefficient is *r* = 0.55 (IMR90), 0.71 (GM12878), and 0.85 (CH12-LX). (*D*) According to the NCI value, we annotated each 100-kb region of human chromosomes with three labels: type-A, type-B, and type-u.

First, we define the property of each chromatin region using NCI; the 100 kb region with *Z*_*w*_ ≥ 0.3 is called the type-A region, where *Z*_*w*_ is the *Z* score of NCI. The region with *Z*_*w*_ ≤ −0.3 is called the type-B region, and the region with −0.3 < *Z*_*w*_ < 0.3 is called the type-u region. Type-A (type-B) regions are abundant in euchromatin (heterochromatin). Because the typical size of the loop domain is ~ 200 kb [5], the larger NCI in a type-A region implies the more frequent intra-domain contacts. These intra-domain contacts may arise from DNA-protein complexes organized for transcription or replication [28, 29], and cohesin molecules off the CTCF-bound sites can associate with these complexes [30, 31] to reinforce contacts and enhance NCI. A type-u region represents either the region showing the intermediate feature between euchromatin and heterochromatin as was identified by clustering the Hi-C contact data [32] or the mosaic of type-A and B regions averaged over a 100-kb interval.

We use the type-A/B/u sequence (Fig. 1D), derived from the local Hi-C data, as the input into our whole-genome polymer simulation. We consider heteropolymer chains connecting type-A, B, and u beads by springs, with each bead representing a 100-kb chromatin region. Then, we compare the predicted results from the polymer simulation with the experimentally observed global Hi-C contact data. These global data have a much larger size than the input; therefore, this comparison between the simulated and observed data should give a stringent test on the physical assumptions adopted in the simulation. We note that the other definitions of the sequence of chromatin properties, such as the histone modification patterns [33, 21, 34], are compatible with the present polymer model as the alternative input into the simulation. In the present study, we use the type-A/B/u sequence instead of the other definitions to restrict ourselves to using only the local physical chromatin properties as the input.

### Effective interactions between chromatin regions

Interactions between type-A/B/u regions should reflect their physical and molecular properties. In particular, the H3K9 methylated nucleosomes in heterochromatin-like type-B regions bind HP1 proteins, which glue nucleosomes to induce the effective attraction between nucleosomes [35, 36]. Hence, the previous computational models assumed attractive interactions between coarse-grained heterochromatin-like regions [12, 13, 27, 22, 14, 15]. However, the effective interactions between coarse-grained regions consisting of hundreds of nucleosomes may differ from those between individual nucleosomes. Indeed, analyses of polymer systems showed that coarse-grained interactions largely depend on density [37, 38, 39]; in high polymer density, the coarse-grained interactions between polymer chains can be repulsive even with attractive interactions between polymer segments [39]. Therefore, care should be taken to model the interactions between coarse-grained chromatin regions, particularly when chromatin density is high.

Here, we estimated the effective interactions between 100-kb type-A/B/u regions by modeling chromatin with bead-and-spring chains with a 1-kb resolution. We considered type-A (B) chains in which all beads are type-A (B). Each chain has 500 kb length, and each bead in the chain represents a 1-kb chromatin segment. We considered the system consisting of 8 ~ 30 chains in a box with the periodic boundary condition (Figs. 2A and 2B) with the potential,

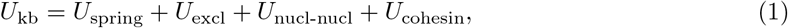

where *U*_spring_ represents springs connecting neighboring beads in each chain, and *U*_excl_ represents the repulsive volume-excluding interactions between beads.

**Figure 2:**
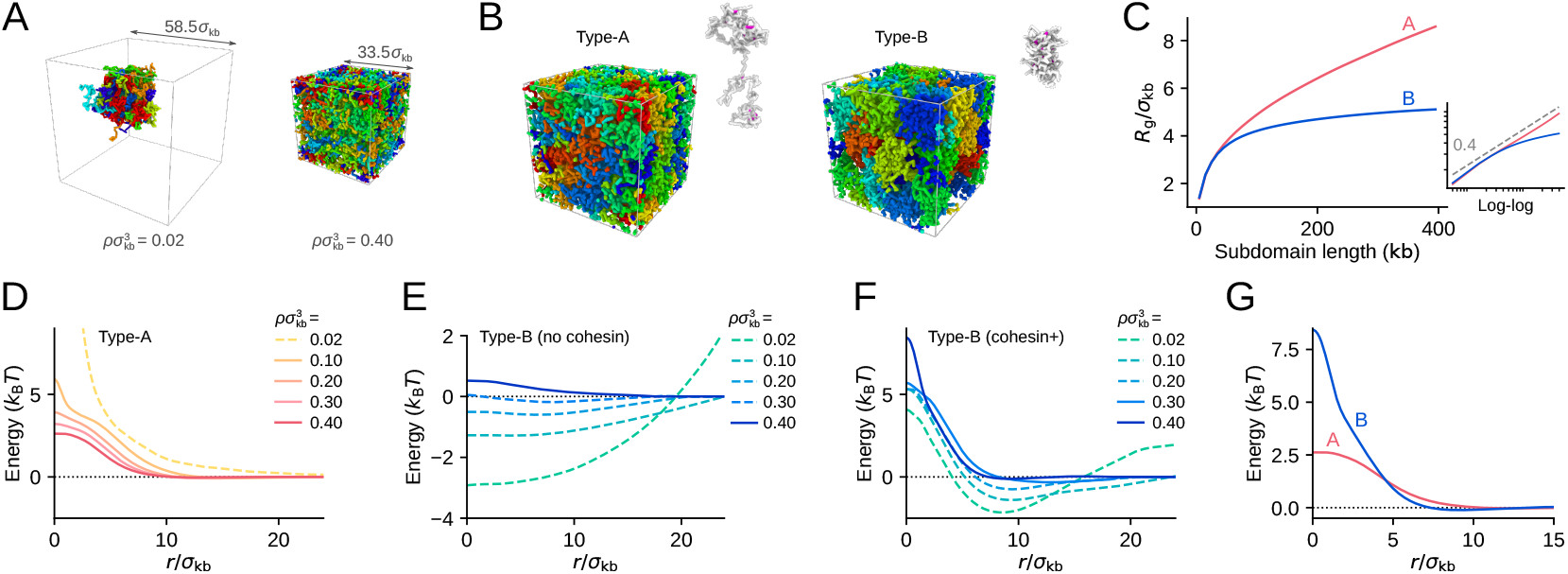
The 1-kb resolution polymer model to estimate the coarse-grained interactions between 100-kb chromatin regions. (*A*) Eight (left) and 30 (right) chains of 500-kb type-B chromatin are confined in periodic boxes of different size. Left: chains in chromatin density 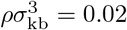 in units of the number density of beads (*ρ* = 0.47 Mb*/μ*m^3^ with the bead diameter *σ*_kb_ = 35 nm), showing chromatin condensation. Right: chains in 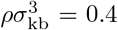(*ρ* = 9.3 Mb/*μ*m^3^), showing repulsion in a crowded system. Cohesin and HP1 are absent in both simulations. Different chains are colored differently. (*B*) Example configurations of 500-kb type-A (left) and type-B (right) chains picked out from the simulated system. Cohesin molecules bundle loops in each chain (magenta). (*C*) The simulated radius of gyration, *R*_g_, of type-A (red) and type-B (blue) chains is plotted as a function of the length *l* (in units of kb) of subdomain used for measuring *R*_g_. Inset is the log-log plot of the same data. A dashed line is *l*^0.4^, showing *R*_g_ of type-B chains is not fitted to a single exponent curve. (*D*) The coarse-grained interaction potential *U*_CG_(*r*) between 100-kb type-A regions obtained with the PRISM theory (solid lines) and the potential of mean force (PMF, a dashed line) in various chromatin density (shown in the legend in units of the number density of beads). (*E* and *F*) The coarse-grained interaction potentials *U*_CG_(*r*) between 100-kb type-B regions obtained with the PRISM theory (solid lines) and PMF (dashed lines) in various chromatin density (shown in the legend in units of the number density of beads). Cohesin and HP1 are absent in *E* and present in *F*. (*G*) *U*_CG_(*r*) between type-A regions (red) and *U*_CG_(*r*) between type-B regions (blue) are superposed. In *D*-–*G*. noise was smoothed out by the Gaussian filter of the size *σ*_kb_*/*2 (*SI Appendix* SI Text and Fig. S1). In *B*, *C*, and *G*, 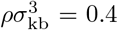. Cohesin is present in the system shown in *B*–*D*, *F*, and *G* and is absent in *A* and *E*

We assumed the short-range bead-bead attraction arising from the interactions between nucleosomes. These interactions depend on the stochastically varying configuration of nucleosomes [40] having a lifetime of ≲ 100 ms [35]. When HP1 binds, such nucleosome-nucleosome association is stabilized with the extended lifetime of ~ 500 ms [35], enhancing the attractive interactions. *U*_nucl-nucl_ in Eq. 1 represents these stochastic attractive interactions in type-B chains. On the other hand, in type-A chains, histone tails are often acetylated, diminishing the attractive nucleosome-nucleosome interactions [41], and HP1 proteins should avoid binding to the acetylated histone tails; therefore, we put *U*_nucl-nucl_ = 0 in type-A chains. We further considered the effects of the binding of cohesin to chromatin chains. Cohesin can bundle the chain into a loop, inducing effectively attractive interactions between the bundled regions. We represented this effect with *U*_cohesin_ by assuming that cohesin stochastically moves along the chain, extruding a loop of the chromatin chain. In type-A chains, various functional complexes should block the cohesin movement [30, 31] as found in the large NCI (Fig. 1), while the cohesin movement is less disturbed in type-B chains [32]. We modeled such difference in the cohesin movement along chromatin chains in the 1-kb resolution model. We simulated the thermally fluctuating ensemble of chains with the 1-kb resolution to derive the coarse-grained (CG) interactions between 100-kb chromatin regions. See *SI Appendix*, SI text for the detailed explanations of simulations.

When we assume the binding of cohesin to chromatin chains and the bead diameter *σ*_kb_ = 35 nm, the calculated spatial distribution of chains showed the radius of gyration, *R*_g_ ≈ 230 nm (160 nm) for 200-kb type-A (B) regions (Figs. 2B and 2C). Importantly, *R*_g_ of type-A chains is scaled as *R*_g_ ~ *l*^0.4^ for the length *l* of subdomain for measurement in the chain, while *R*_g_ of type-B chains is not fitted to a single exponent curve, showing a saturating behavior as a function of *l* (Fig. 2C). These behaviors of *R*_g_ are consistent with the microscope observations of active and repressed chromatin regions [8].

From the calculated radial distribution *g*(*r*) between 100 kb regions in the 1-kb resolution model, we obtained the potential of mean force (PMF), *U*^PMF^(*r*) = −*k*_B_*T* log *g*(*r*). When the chromatin density is sufficiently low, PMF represents the CG interaction between chromatin regions separated by a distance *r*. However, in high chromatin density, many-body effects modulate the PMF. We eliminated these many-body effects by calculating the direct correlation function with the polymer reference interaction site model (PRISM) theory [42], from which we derived the intrinsic pairwise potential *U*^PRISM^(*r*). Therefore, the coarse-grained interactions *U*_CG_(*r*) with a 100-kb resolution are approximated by *U*^PMF^(*r*) in low density and *U*^PRISM^(*r*) in high density. See *SI Appendix*, SI text for the application of the PRISM theory.

In Figs. 2D–2F, we show *U*_CG_(*r*) calculated in various chromatin density *ρ*. For type-A regions with cohesin (Fig. 2D), *U*_CG_(*r*) has a Gaussian-like form showing the repulsive interaction between type-A regions. Without cohesin, the type-A regions are more extended, leading to a milder repulsion with the smaller amplitude of *U*_CG_(*r*). This repulsive interaction is consistent with the observation that chromatin does not condense in vitro when histone tails are acetylated [23]. For type-B regions in the absence of cohesin and HP1 (Fig. 2E), *U*_CG_(*r*) showed a distinctly attractive interaction for density 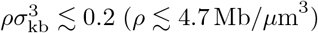, consistent with the in-vitro observation that chromatin is condensed and phase-separated from solvent when histone tails are unacetylated [23]. This attractive interaction is due to the nucleosome-nucleosome interaction, which is further enhanced when HP1 binds to type-B chains. However, in high chromatin density with 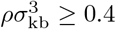, *U*_CG_(*r*) showed a mildly repulsive interaction.

Type-B chains with cohesin and HP1 showed the attractive *U*_CG_(*r*) for 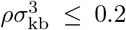, while *U*_CG_(*r*) becomes dominantly repulsive in 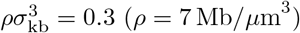 and repulsive in 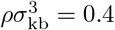 (*ρ* = 9.3 Mb*/μ*m^3^) (Fig. 2F). Cohesin binding to type-B chains compacts each loop domain with the effective intra-loop attraction (Fig. 2B). For inducing the attractive inter-domain interactions, the domain conformation needs to be loosened with some free energy cost to allow the inter-domain HP1 bridging. This effect is particularly evident when domains are compacted in high chromatin density. Therefore, the tighter domain compaction with cohesin binding in high chromatin density diminishes the inter-domain attraction, emphasizing the repulsive interaction. The repulsive slope of the potential becomes steeper as the cohesin density is larger (*SI Appendix*, Fig. S1B). Density of chromatin in human cells, *ρ* ≳ 10 Mb*/μ*m^3^, is high enough to exhibit repulsive interactions. Thus, with the cohesin and HP1 binding, the repulsive interaction between type-B regions arises.

In our whole-genome simulation, we used mathematically convenient forms of potentials *U*_AA_(*r*) and *U*_BB_(*r*) (Methods), which capture the main features of the interactions *U*_CG_(*r*) obtained with the 1-kb resolution model and the PRISM theory (Fig. 2G); the interaction between type-A regions *U*_AA_(*r*) and the interaction between type-B regions *U*_BB_(*r*) are both repulsive with the steeper slope in *U*_BB_(*r*). We set the potential width of *U*_AA_(*r*) and *U*_BB_(*r*) similar to the calculated values in *U*_CG_(*r*); as two chromatin regions approach, *U*_CG_(*r*) start to rise at *r* ≈ 300 nm for type-A regions and 250 nm for type-B regions (Fig. 2G). The calculated height of the potential was *U*_CG_(*r* ≈ 0) ≈ 2.5*k*_B_*T* between two type-A regions. Accordingly, we set the potential height *U*_AA_(*r* = 0) = 2.5*k*_B_*T* . *U*_CG_(*r* ≈ 0) between two type-B regions was considerably larger than that between two type-A regions. We did not use this large value in the whole-genome model because the PRISM theory only derives the inter-chain potential, but we used the same potential functions for the intra- and inter-chain type-B interactions in the whole-genome model. For explaining the intra-chain structure consistently, we assumed the lower height of *U*_BB_(0) than that of *U*_CG_(0). Later in this paper, we discuss the effects of varying *U*_BB_(0) by comparing the calculated results with the experimental Hi-C data (*SI Appendix*, Fig. S11). Fig. 3A shows the functional forms of the potentials used in the whole-genome model; *U*_AA_(*r*) is Gaussian-like, and *U*_BB_(*r*) is a harder repulsive interaction. The repulsion between two type-u regions was assumed to be 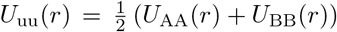, and we set 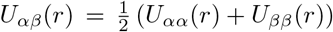 with *α* and *β* being A, B, or u.

**Figure 3:**
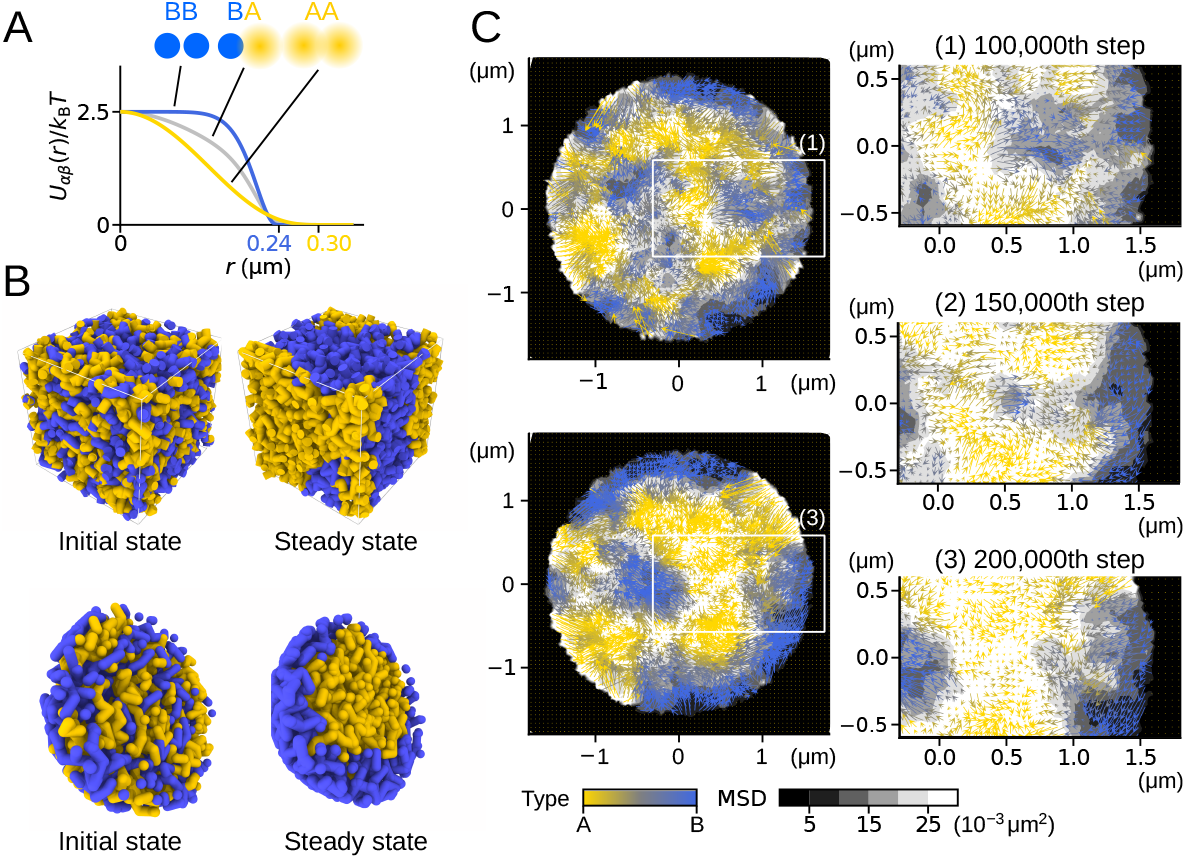
Heterogeneous repulsive interactions and phase separation. (*A*) Potential functions between type-A, B or u regions in distance *r*; *U*_AA_(*r*), *U*_BB_(*r*), and 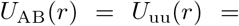 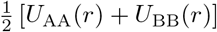. (*B* and *C*) A polymer-blend system was simulated to demonstrate the phase separation induced by heterogeneous repulsive interactions. 100 chains each composed of 20 segments of type-A (yellow) and 100 chains each composed of 20 segments of type-B (blue) were mixed and subjected to the Brownian motion. (*B*) Polymer chains mixed in a box with the periodic boundary condition (top) and in a spherical container shown with the semi-sphere view (bottom). Left: the initial configuration. Right: a snapshot after the system reached a steady state. (*C*) Left: cross-section of the sphere at the 1 × 10^5^th step (top left) and 2 × 10^5^th step (bottom left). Right: close-up views of the rectangular area in the left. Fluctuation (gray-scaled back ground, the 10^3^-step mean-square displacement (MSD)) and flow (arrow, the 10^5^-step displacement) of polymer segments averaged within each 0.12-*μ*m region around the mesh points are shown. Colors of flows represent the ratio of A/B density. The images were rendered using OVITO [43].

### Phase separation driven by the repulsive interactions

The interactions *U*_*αβ*_(*d*) induce phase separation of type-A and B regions when we confine them in a high-density space. We demonstrate this behavior by using a simple model of the polymer blend consisting of polymers of type-A segments and polymers of type-B segments. The Brownian motion of these mixture polymers induces phase separation (Fig. 3B). The mean square displacement (MSD) analysis (grayscale in Fig. 3C) shows that the type-B segments are packed in a more solid-like manner, whereas the type-A segments show a more fluid behavior. The large fluctuations of type-A segments allow the type-A segments to merge into the phase-A domain. In particular, in the system confined in a rigid sphere, the type-A segments occupy the inner region to acquire the volume allowing the motion while the type-B segments are packed at the periphery analogously to the heterochromatin/euchromatin separation in cells (Figs. 3B and 3C).

### The whole-genome simulation

We simulated the whole-genome structrue and dynamics of human cells by using the annotated sequence of type-A/B/u (Fig. 1D) and the repulsive potentials *U*_*αβ*_ with *α, β* = A, B, or u (Fig. 3A).

During interphase in the human nucleus, a chromosome is displaced at most ~ 2 *μ*m [46], a much shorter distance than the nuclear size; the system is neither stirred nor equilibrated during interphase. Therefore, to explain genome organization in interphase, the process of structure formation at the entry to the G1 phase needs to be carefully considered [47]. We simulated anatelophase genome by pulling a centromere locus of each condensed chromosome chain toward one direction in the model space (Fig. 4A). Then, from the thus obtained configuration, we started the simulation of decompression by assuming the disappearance of the condensin constraints at this stage, which allowed chromosomes to expand with repulsion among chromatin chains (Fig. 4A, *SI Appendix*, Movies S1–S3). The nuclear envelope forms during this expansion [48]. We simulated this envelope formation by assuming a spheroid (IMR90) or sphere (GM12878) surface, whose radii varied dynamically by balancing the pressure from outside the nucleus and the one arising from the repulsion among chromatin chains. We repeated this simulation 200 times with diffrent random number realization. These 200 trajectories represent an ensemble of diffferent 200 cells.

**Figure 4:**
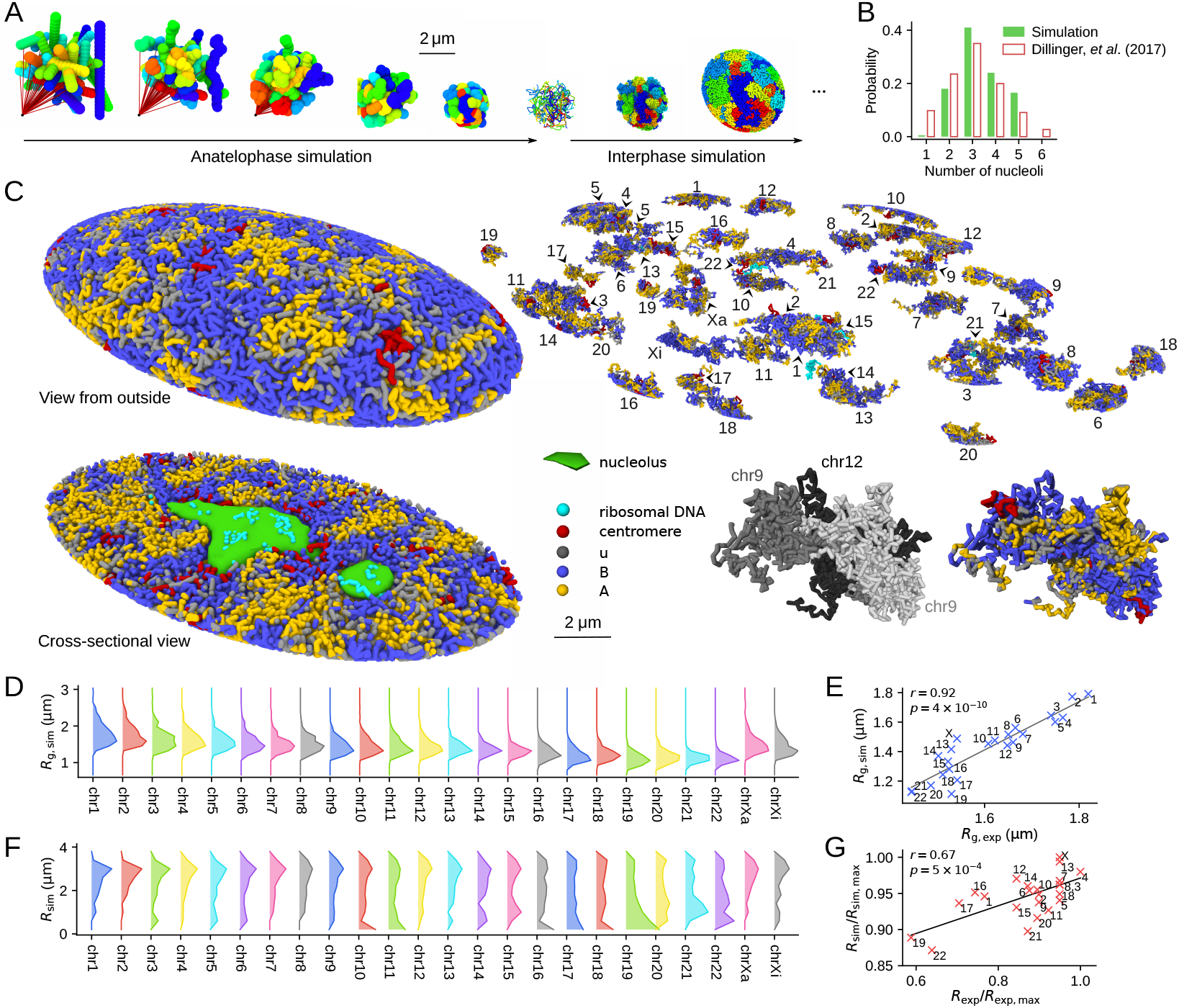
The simulated human genome. (*A*) The anatelophase simulation of the genome and the subsequent expansion of the nucleus at the entry to interphase. 46 chromosome chains are colored differently. From left to right: chromosomes were represented by 46 rods placed in random orientations; the centromere locus of each chromosome was dragged by a spring (red line), whose end was anchored to a point in the space; chromosome chains with a 10 Mb resolution were subjected to the thermal fluctuations during this dragging process; the chains were fine-grained to 100 kb resolution and equilibrated; and expansion of the genome was simulated as the entry process to G1. (*B*) Distribution of the population fraction of cells having the number of nucloli designated on the *x*-axis. Comparison of the simulated 200 cells (green) (*SI Appendix*, SI text and Fig. S2) and the experimental data (white bars with red outlines) [44]. (*C*) A snapshot of the simulated G1 nucleus of IMR90. A view from the outside, the cross-section view, individual chromosomes, and the close-up view of three associated chromosomes are shown. A homologous pair of chromosome 9 and chromosome 12 occasionally associated in this simulated cell, showing an example case that the compartments spread across chromosomes. The genome is phase-separated into type-A (yellow), type-u (gray), and type-B (blue) chromatin regions and the nucleoli (green). Centromeres (red) are found near the type-B regions and rDNA loci (cyan) are near the type-B regions or buried in the nucleoli. (*D*) Distribution of *R*_*g*_ of chromosomes in GM12878 cells. Sampled from 200 simulated cells. (*E*) *R*_*g*_ averaged over the distribution in *D* is compared with the microscope data [9]. The correlation coefficient is *r* = 0.92 with *p* = 4 × 10^−10^. (*F*) Distribution of the radial position of the center of mass of chromosomes in GM12878 cells. Sampled from 200 simulated cells. (*G*) The two-dimensional radial positions of the center of mass of chromosomes are compared with the microscopy data [45]. The simulated chromosomes were projected onto a two-dimensional plane to calculate the two-dimensional radial positions. The correlation coefficient is *r* = 0.67 with *p* = 5 × 10^−4^. In *A* and *C*, the images were rendered using OVITO [43].

The nucleoli also form during this expansion [49]. We represented the nucleoli by assemblies of particles adhering to rDNA, into which the transcription products and the related factors are condensed. The weak short-range, attractive interactions arising from the exchange of diffusive molecules were assumed between the particles, which spontaneously assembled to form nucleoli through the genome expansion process. The number distribution of the resultant nucleoli per cell shows a peak at 3 and fluctuates around 2 ~ 5, showing the agreemet with the microscope observation [44] (Fig. 4B). Each chromosome gained a V-shaped conformation during anatelophase, whose effects remained through the expansion process, leading to the long-range contacts between p- and q-arms of chromosomes in interphase. The nonequilium nature or the memory of the mitotic phase in the interphase genome architecture is consistent with the correlation between the chromosome configuration in the mitotic phase and that in the interphase observed among vastly different organisms [50].

Fig. 4C shows a snapshot of the simulated structure obtained after the nucleus reached a stationary size. In this state, the genome is phase-separated into type-A, B, and u regions and nucleoli, where the type-u regions reside at the boundary of the type-A and B regions. Phase-separated A/B regions are formed in each chromosome and across chromosomes. We should note that this state is a nonequilibrium stationary state obtained by keeping the balance of pressures from outside and inside the nucleus. In this state, chromosomes were not mixed significantly but formed territories because the pressure balance was acquired before the slow entangling and mixing of chromosomes proceed. Size of chromosome territories was measured by the radius of gyration *R*_*g*_ of chromosomes. Distribution of *R*_*g*_ of each chromosome in the stationary 200 cells is plotted in Fig. 4D, showing the extent of cell-to-cell fluctuation. The mean *R*_*g*_ of the distribution agrees with the microscopy data [9] with the correlation coefficient *r* = 0.92 (Fig. 4E). Distribution of the radial position of chromosome territories, i.e., the radial position of the center of mass of chromosomes, also fluctuates from cell-to-cell, but the distribution shows a distinct tendency that chromosomes with the smaller size reside in the inner region of the nucleus, and particularly, chr19 tends to be buried inside the nucleus (Fig. 4F). The averaged radial position is compared with the microscopy data [45], showing the correlation coefficient *r* = 0.67 (Fig. 4G).

### Genome organization generated through phase separation of chromatin

We analyzed the generated structures by comparing the experimentally observed [5] and simulated Hi-C contact matrices. The simulated data reproduces the observed features of contacts among several chromosomes (Fig. 5A) and in the whole genome (*SI Appendix*, Fig. S3), while the noise arising from the small sample size (*n* = 200) of simulated cells remained, reflecting the intense cell-to-cell fluctuation in inter-chromosome contacts. The noise was removed by enhancing the signal/noise ratio with the correlation-coefficient representation of the contact matrix [4], clarifying the agreement between the simulated and observed results (Fig. 5B). Comparisons for the intra-chromosome contacts show a further agreement between the simulated and observed results (Fig. 5C, *SI Appendix*, Figs. S4–S7). Diagonal blocks in the intra-chromosomal matrices represent chromatin domains with several Mb or the larger size. The simulated and observed block patterns agree with each other, suggesting these chromatin domains arise from phase separation (*SI Appendix*, Movies S4, S5). The observed dependence of the contact frequency *P* (*s*) on the sequence distance *s* is well reproduced in the simulation (Fig. 5D), showing that the simulation explains both the global and local chromosome structures in a balanced way. The plaid pattern of the inter- (Figs. 5A, 5B) and intra-chromosomal (Fig 5C) contact matrices represents A/B compartmentalization. This A/B compartmentalization was quantified using the compartment signal. The compartment signals derived from the simulated data were superposed with the observed data with a Pearson’s correlation coefficient of *r* ≈ 0.8 (Fig. 6A). The genome-wide compartment signals for IMR90 and GM12878 show that the simulation reproduces the genome-wide data (Fig. 6B). Therefore, our model explains the A/B compartments and the other features of the intra- and inter-chromosome Hi-C contacts in different cells using the same single set of model parameters. These features were lost when type-A, B, and u loci were randomly assigned along the polymer chains (*SI Appendix*, Fig. S8), showing that the arrangement of local properties along the chromatin chain is essential for proper phase separation and genome organization. Results in Figs. 4–8 were obtained by sampling the data from the calculated 200 trajectories, each having the length of ~ 7h. Results were not much altered when we sampled the data from the trajectories of ~ 14 h (*SI Appendix*, Fig. S9). The results were robust against the change in the definition of the type-A/B/u annotation. In *SI Appendix*, Fig. S10, we show the results obtained by categorizing chromatin regions into two types, A and B, as type-A for *Z*_*w*_ > 0 and type-B *Z*_*w*_ ≤ 0. Effects of varying the functional form of *U*_BB_(*r*) is shown in *SI Appendix*, Fig. S11. The gentler slope of the repulsive potential with the higher value of *U*_BB_(0) also gave the phase-separated genome structure, but the contrast of phase separation was weakened with the gentle repulsive force between type-B regions; the distinct heterogeneity of repulsive force, i.e., soft between type-A regions and hard between type-B regions, is a driving force of phase separation of chromatin.

**Figure 5:**
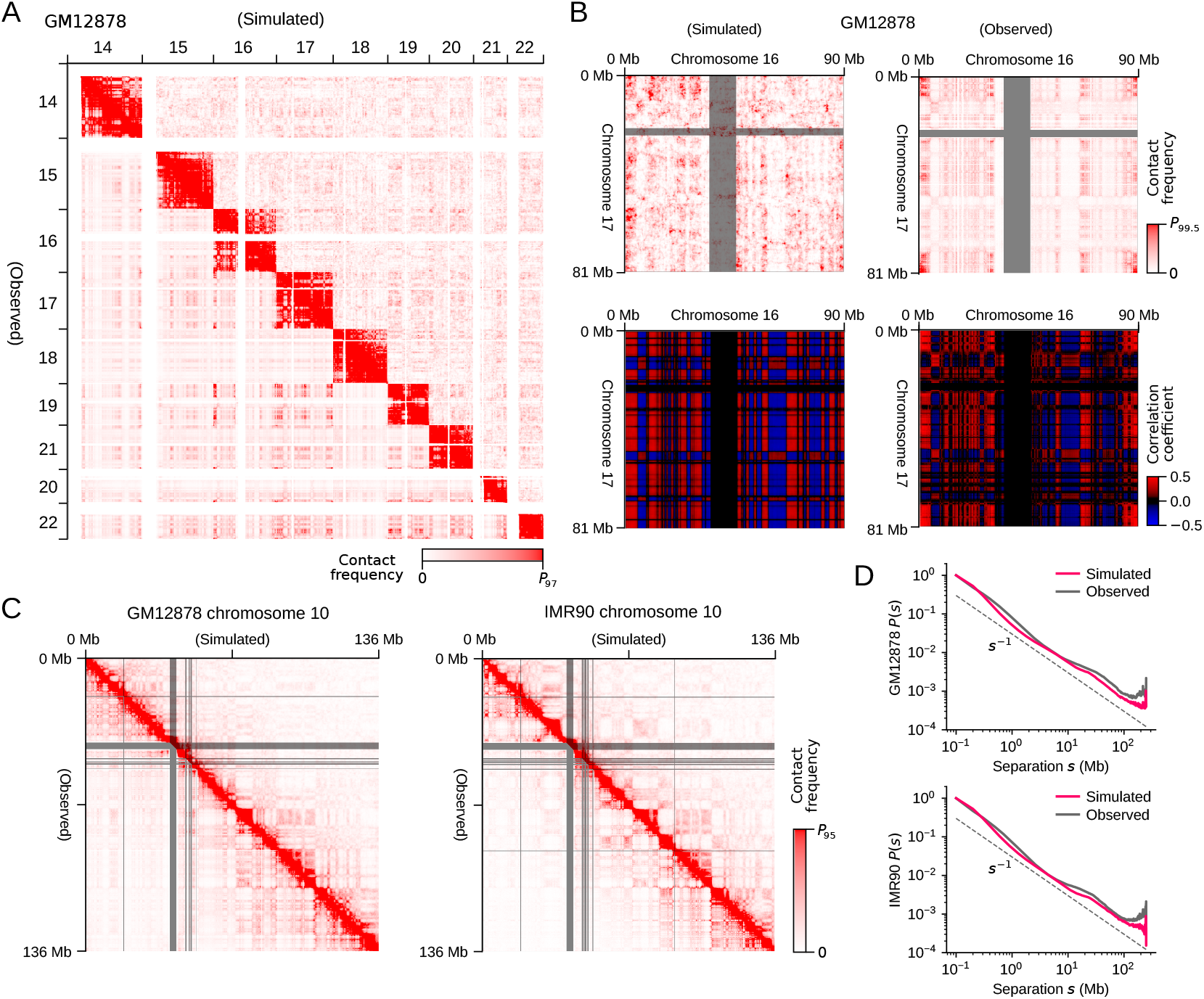
Comparisons of the simulated and experimentally observed chromatin contact frequencies. (*A*) The simulated (upper right) and observed (lower left) contact matrix among several chromosomes of GM12878. (*B*) The simulated (left) and observed (right) inter-chromosome contacts between chromosomes 16 and 17 of GM12878 (top) and those represented with the correlation-coefficient matrix [4] (bottom). (*C*) The simulated (upper right) and observed (lower left) intra-chromosome contact matrix of chromosome 10. (*D*) The simulated (red) and observed (black) contact frequency *P* (*s*) averaged over the genome for the sequence separation *s*. Data are plotted in a 1 Mb resolution in *A* and *B*, and a 100 kb resolution in *C*. In *A*-*C*, the experimental data are lacking in the regions designated by gray bars. The experimental data are from Ref.[5].

**Figure 6:**
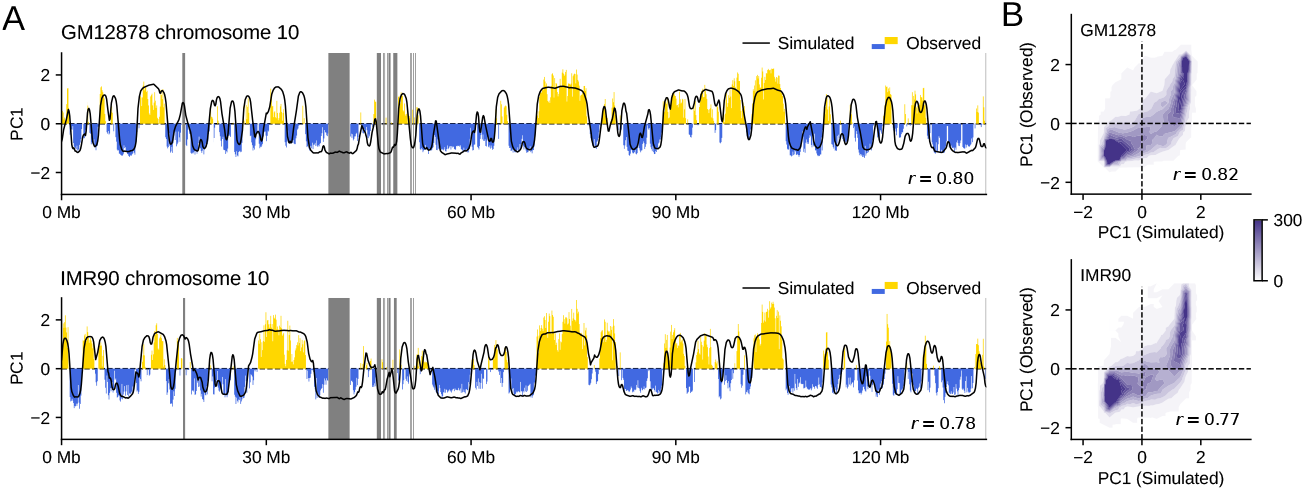
Comparisons of the simulated and experimentally observed compartment signals. (*A*) The simulated (black) and observed (yellow and blue) compartment signals of chromosomes 10 of GM12878 (*r* = 0.80) and IMR90 (*r* = 0.78). The experimental data are lacking in the regions designated by gray bars. (*B*) Contour plot of the density of compartment signals of the whole genome on the plane of the simulated and observed signal values for GM12878 (*r* = 0.82) and IMR90 (*r* = 0.77). Density is the number of 100-kb chromatin segments in a bin of 0.1 × 0.1 square on the plane. The experimental data are from Ref.[5]. Compartment signal is the first principal component (PC1) of the simulated or observed chromatin contact matrix.

We further analyzed the generated genome structure by examining the lamina-chromatin association. Fig. 7A shows that type-B chromatin accumulates, while type-A chromatin depletes near the nuclear envelope (Methods). The specific lamina-chromatin attractive interactions are not considered in the present model; therefore, this accumulation was not due to the tethering of type-B chromatin to the lamina but was induced dynamically similarly to the simulated polymer-blend system (Figs. 3B, 3C). Fig. 7B compares the simulated and experimentally observed [51] lamina-chromatin association data for an example chromosome. The loci showing a large frequency of association, i.e., the lamina-associated domains (LADs), are reproduced by the present simulation. LADs in the other chromosomes are shown in *SI Appendix*, Fig. S12. Fig. 7C compares the genome-wide data showing an agreement between the simulated and observed [51] lamina-chromatin association with *r* = 0.55. Specific factors that tether the LADs to the lamina should delay the dissociation kinetics of the LADs from the lamina, but our simulation revealed that the lamina-LAD association resulted as a consequence of the dynamic genome-wide phase separation process. We found similar results around the nucleoli. The simulation reproduced the observed [44] nucleolus-associated domains (NADs) though the chromatin chains except for the rDNA loci were not tethered to the nucleoli in the model (Fig. 7D). NADs in the other chromosomes are shown in *SI Appendix*, Fig. S13. The simulated and observed genome-wide nucleoli-chromatin associations show an agreement with *r* = 0.70 (Fig. 7E). We note that the A/B compartments were well reproduced even when the system was simulated with the absence of nucleoli (*SI Appendix*, Fig. S14), showing that the nucleoli are not the driving force of phase separation of chromatin, but a perturbation to the genome structure.

**Figure 7:**
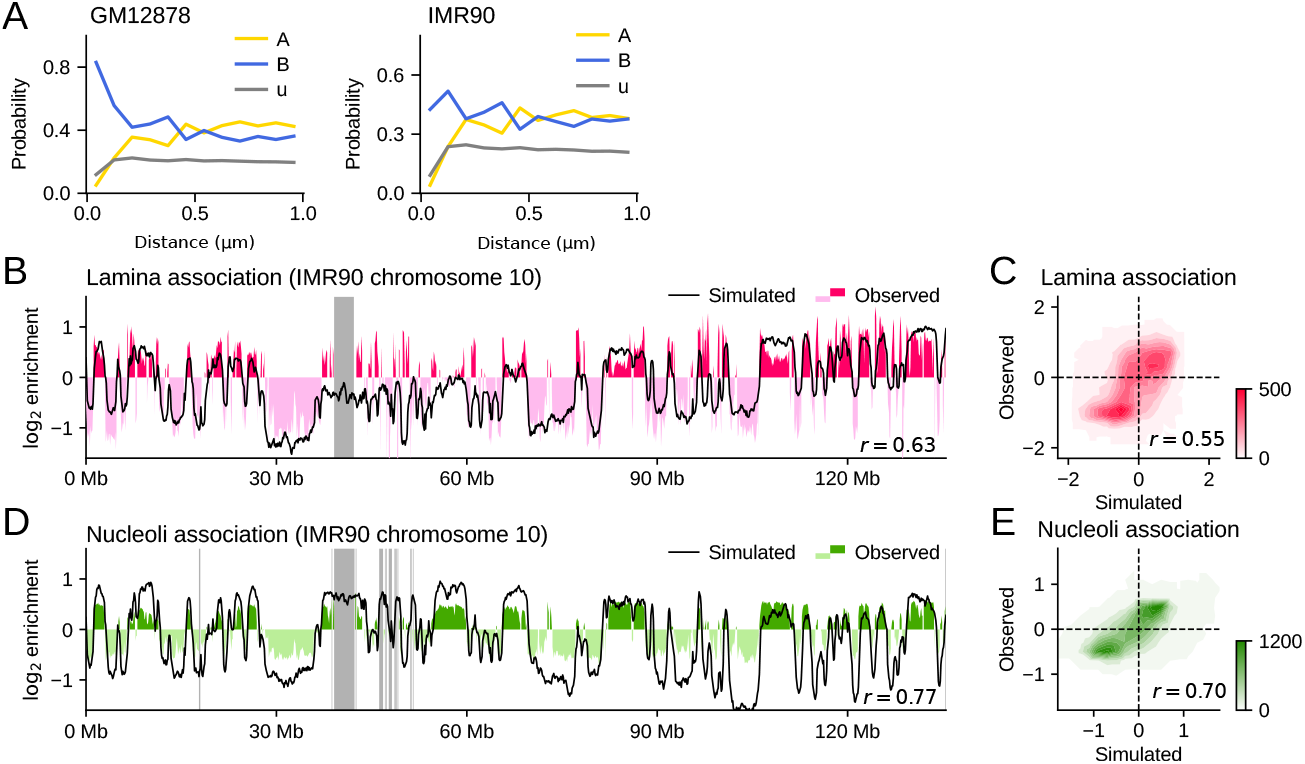
Comparisons of the simulated and experimentally observed data of association of chromatin with nuclear structures. (*A*) The probability to find the type-*α* chromatin with *α* = A (yellow), u (gray), and B (blue) are plotted as functions of distance from the nuclear envelope (Methods). GM12878 (left) and IMR90 (right). The data from *B* to *E* are for IMR90. (*B*) The simulated (black) and observed (red and pink) lamina-chromatin association for chromosome 10. *r* = 0.63. (*C*) Contour plot of the density of the genome-wide lamina-chromatin association on the plane of the simulated and observed values. *r* = 0.55. (*D*) The simulated (black) and observed (green and thin green) nucleoli-chromatin association for chromosome 10. *r* = 0.77. (*E*) Contour plot of the density of the genome-wide nucleoli-chromatin association on the plane of the simulated and observed values. *r* = 0.70. In *C* and *E*, density is the number of 100-kb chromatin segments in a bin of 0.1 × 0.1 square on the plane. In *B* and *D*, the experimental data are lacking in the regions designated by gray bars. The experimental data of *B* and *C* are the ChIP-seq data for Lamin B1 [51]. The experimental data of *D* and *E* are from [44].

### Dynamic fluctuations of the genome

After the simulated genome reached a stationary G1 phase, there remained fluctuations in genome movement such as the dynamic feature of the simulated lamina-chromatin interactions (Fig. 8A). The lamina-chromatin contacts spread as the genome expanded from *t* = 0 min. After the genome became stationary at *t* ≈ 200 min, dynamic fluctuations including association, dissociation, and positional shift of lamina-chromatin contacts continued to take place. A pair of homologous chromosomes showed differing fluctuating patterns from each other. We analyzed these fluctuations by plotting the temporal change of the normalized spatial distribution of chromatin loci after they adhered to the lamina (Fig. 8B, Methods). The distribution became broad over time, reflecting the dynamic dissociation of chromatin from the lamina as observed in the single-cell experiment [52]. The root-mean-square distance (RMSD) of chromatin loci from the lamina is plotted as a function of time passed after each chromatin locus attached to the lamina, showing RMSD ~ *t*^*α*^ with *α* ≈ 0.35 for both type-A and B chromatin but with the larger RMSD for the type-A chromatin (Fig. 8C). With this small *α*, once the loci were attached to the lamina, they stayed within 1 *μ*m of the lamina for a long time as experimentally observed [52]. This reflects the tendency of the phase-separated compartment to retain LADs near the lamina.

**Figure 8:**
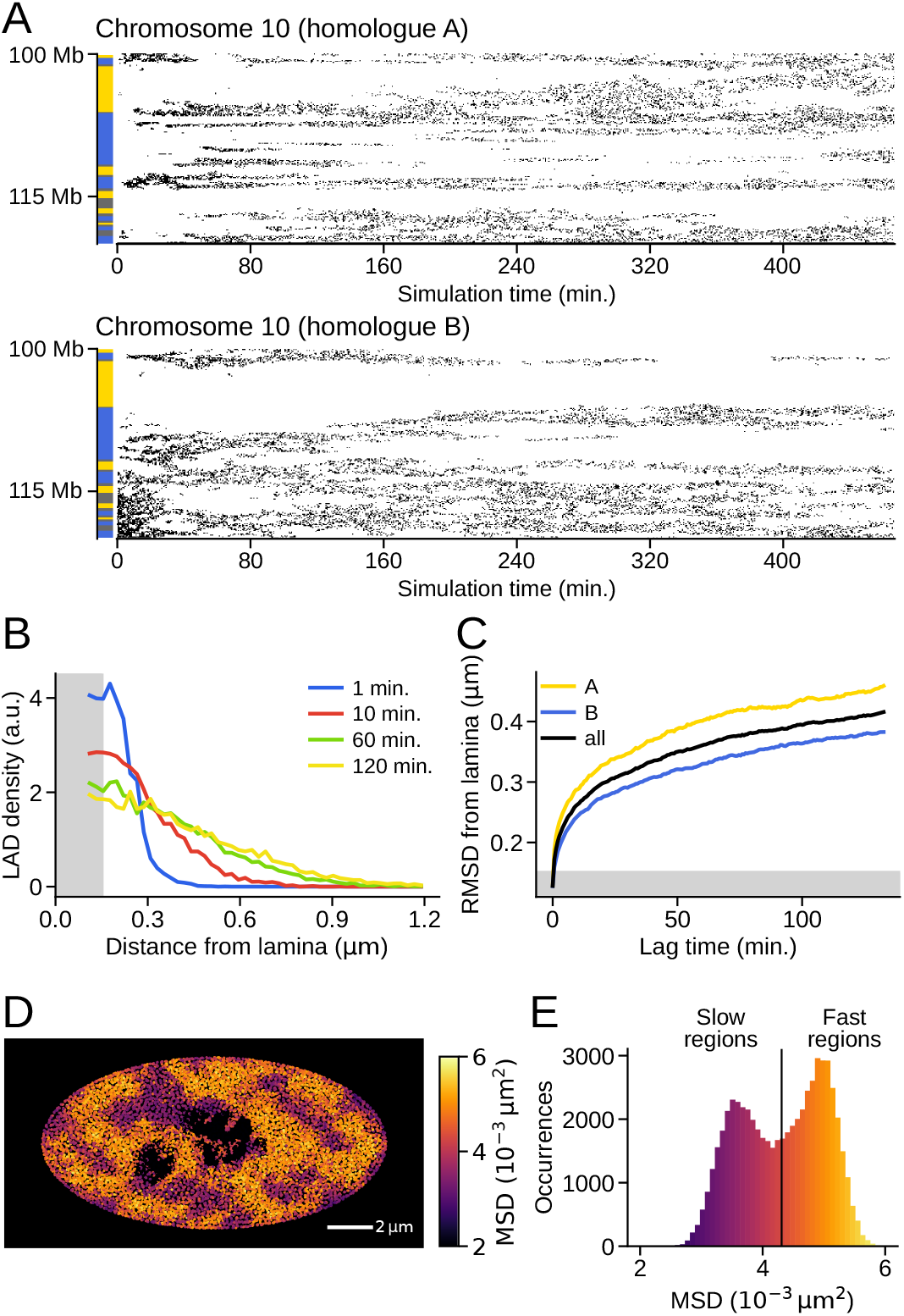
Dynamic fluctuations of the genome structure. The data for the simulated IMR90 nucleus. (*A*) Time-dependent contact development between lamina and chromatin is plotted as points of contact on the plane of time and the sequential position. A pair of homologues of chromosome 10 are compared. Bars on the left show the NCI annotation to distinguish type-A, (yellow), B (blue), and u (gray) regions. (*B*) Temporal change of the normalized distribution (Methods) of distance between lamina and the chromatin loci, where the time duration was measured as time passed after each locus attached to the lamina. (*C*) Temporal change of the root-mean-squared distance (RMSD) between lamina and the chromatin loci after each locus attached to the lamina. The type-A chromatin (yellow), type-B chromatin (blue) and the average (black). In *B* and *C*, the chromatin locus was regarded to be in contact with the lamina when it is in a gray hatched area. (*D*) A snapshot of the cross section with the square displacement of each 100-kb region during Δ*t* = 4 s designated by colors. The dark regions near the center are nucleoli. (*E*) Distribution of the mean square displacement (MSD) of each 100-kb regions during Δ*t* = 4 s. Colors in *E* are the same as in *D*.

Dynamic fluctuations were found in the entire nucleus. Fig. 8D shows a snapshot of the distribution of square displacement of each 100-kb chromatin region during Δ*t* = 4 s. The distribution is heterogeneous with the slow movement at the nuclear periphery and fast movement in the inner regions. Similar heterogeneous distributions were observed using live-cell imaging [53, 10, 11]. Fig. 8E is the distribution of MSD of each chromatin region during Δ*t* = 4 s. The distribution was bimodal as observed with live-cell imaging [54], showing contributions from the fast and slow components. The fast chromatin is mostly type-A, whereas the slow chromatin tends to be type-B (*SI Appendix*, Fig. S15). The pair-correlation functions showed that the positions of fast chromatin are correlated within 1 *μ*m, constituting the fast-moving domains (*SI Appendix*, Fig. S16).

The difference in the movements between type-A and B regions was a driving force of phase separation during the genome expansion process at the entry to the G1 phase. This difference remained in fluctuations of dynamic association/dissociation of chromatin to/from the lamina after the genome reached the G1 phase and in the heterogeneity of the genome-wide dynamic fluctuations of chromatin in interphase nuclei. Coupling of these fluctuations with DNA functions such as transcription [55] and replication [56, 57] is an important issue for which the present model should provide a basis for the analyses.

## Discussion

The simulated results showed that the present whole-genome model helps bridge different experimental analyses, including Hi-C and other high-throughput measurements, as well as live-cell imaging. The model explained various genomic data quantitatively, suggesting that the repulsion-driven phase separation largely determines the genome organization.

The analyses of cohesin-depleted cells with the Hi-C methods [58, 59] and theoretical modeling [60] have shown that A/B compartmentalization discussed in the present study and formation of the sub-megabase cohesin-bound loop domains of chromatin chains or topologically associating domains (TADs) are the two independent mechanisms that characterize the genome organization. In the present whole-genome model, the NCI annotation of local chromatin property represents the effect of loop-domain formation, which should cooperatively determine the intra-loop physical property [61, 62]. However, for analyzing the structure and dynamics of loop domains with its typical size being ~ 200 kb [5], the genome needs to be modeled with a higher resolution than the present 100 kb resolution. Incorporation of the 1-kb resolution model into the present phase-separated whole-genome structure should provide insights into this problem, helping to clarify the interplay between compartmentalization and loop-domain formation/dissolution.

On the other hand, in the larger structure than a 100 kb scale, the determinant role of the phase-separation mechanism implies that tethering of chromatin chains to the lamina, nucleoli, and other droplet-like condensates such as mediator droplets at super-enhancers [55, 63] or nuclear speckles should work as perturbations to the genome structure/dynamics. The present model structure is usable as a starting structure for analyzing these chain-constraining effects and other perturbative effects.

Thus, the repulsion-driven phase-separated structure of the genome provides a platform for analyzing the effects of both small- and large-scale structural effects for comprehensive analyses of the genome organization.

## Methods

### Outline of the interphase genome simulation

We modeled the human genome with 46 heteropolymers; each polymer was a beads-on-a-string chain with a single bead representing a 100-kb chromatin region, which amounts to *N*_chr_ = 60752 beads in the genome. The nucleoli were represented by assemblies of *N*_no_ = 1426 beads. Chromatin and nucleolar beads were confined in a spherical (GM12878) or spheroidal (IMR90) nuclear envelope, whose size was dynamically varied to adjust the balance of pressures inside and outside the nucleus. Dynamical movements of chromatin and nucleolar beads were simulated by numerically solving the overdamped Langevin equation with a discretization time step of *δt* = 10^−5^*τ*_0_ in time units of *τ*_0_ = 1 h. The initial genome conformation was prepared by simulating the anatelophase genome, and the subsequent genome expansion was simulated as the entry process to interphase. The nucleus reached a stationary size after approximately 200,000 steps, and we sampled the genome structure till the 700,000th step. Each simulation run corresponds to a single cell simulation, and we performed *N*_cell_ = 200 independent simulation runs using different random number implementation. See *SI Appendix*, SI text for the detailed explanation.

### Potential functions for repulsive interactions between 100-kb chromatin regions

Repulsive potentials shown in Fig. 3A are

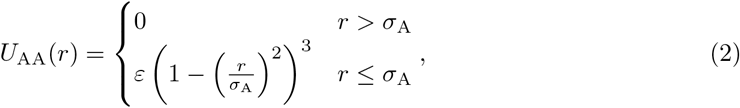

and

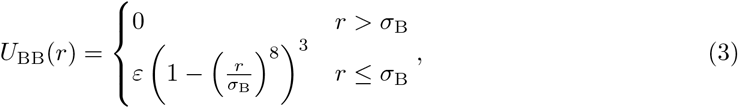

where *r* is the distance between two 100-kb chromatin regions, and *ε* is a measure of the free-energy cost of overlapping of two regions and *σ*_A_ and *σ*_B_ are typical of the spreading of type-A and B resprepulsive forces. The functional form of *U*_AA_(*r*) in Eq. 2 was used because this algebraic form well approximates a Gaussian function while this form shows higher computational efficiency than the Gaussian. The functional form of Eq. 3 was used as an extension of the form in Eq. 2. We set *ε* = 2.5*k*_B_*T* to allow overlapping of chromatin regions with thermal fluctuation, *σ*_A_ = 0.30*A*(*t*) *μ*m, and *σ*_B_ = 0.24*A*(*t*) *μ*m to reproduce chromatin density in nuclei, where *A*(*t*) is a scaling factor that monotonously increases from 0.5 as the genome size increases at the entry to the G1 phase (*SI Appendix*, SI text), and *A*(*t*) ≈ 1 after the genome reached the G1 phase (Fig. 4C).

### Hi-C data of chromatin contacts

#### Simulated Hi-C data

Instantaneous contact matrix *M_ij_* (*t*; *k*) was sampled at time *t* in the *k*th simulation run. A pair of chromatin beads *i* and *j* were considered to be in contact if the beads are closer than a scaled threshold distance:

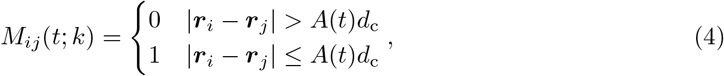

where *A*(*t*) is a scaling factor explained in *SI Appendix*, SI text and *d*_c_ is the threshold distance. We set *d*_c_ = 0.24 *μ*m because the simulated functions of pair correlation in distance peaked at ~ 0.24 *μ*m for all the bead types of A, B and u. *M*_*ij*_ (*t*; *k*) was sampled in every 20 steps from the 20,000th step to the 70,000th step. We write this set of sampling time points as *T*_c_. *M*_*ij*_ (*t*; *k*) was summed into

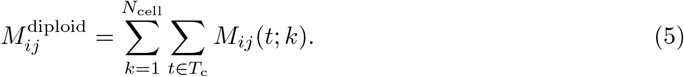

Here, the population contact matrix 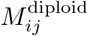 defined above differentiates the alleles on two homologous chromosomes. In order to compare the results with the conventional Hi-C experimental data, we averaged out the homologous contact frequency. With sites *i*′ and *j*′ denoting the homologous copies of sites *i* and *j*, respectively, we calculated a smaller contact matrix,

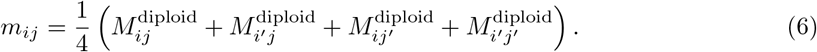

#### Deriving Hi-C contact matrix and compartment signal from the simulated or the experimentally observed data

From the experimentally observed [5] or simulated raw contact matrix, *m*_*ij*_, the ratio 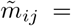 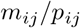 was calculated as described in Ref.[4]. Here, *m*_*ij*_ is the observed or simulated Hi-C contact counts, and *p*_*ij*_ is the expected contact counts between *i* and *j*. For intra-chromosome contacts, *p*_*ij*_ = *F* (*s*) is the mean contact counts for the sequence separation *s* = |*i* − *j*| with its average taken over the genome. For inter-chromosome contacts, we used *p*_*ij*_ = *N*_read_*f*_*i*_*f*_*j*_. Here, *N*_read_ is the total number of inter-chromosome contact counts in the genome and 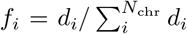 is the fraction of contacts involving site *i* with *d*_*i*_ = Σ_*k*∈other chr_ *m*_*ik*_ where the inter-chromosomal sum over *k* was taken genome-wide excluding intra-chromosome sites. *N*_chr_ is the number of chromosome loci in the genome.

In Fig. 5B, the inter-chromosome contacts were analyzed by showing the normalized correlation-coefficient matrix,

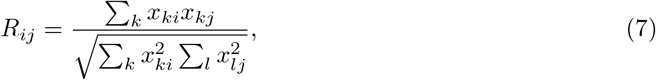

where 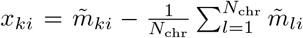. The compartment signal shown in Figs. 1 and 6 is the first principal component vector (PC1) of the whole-genome contact matrix, i.e., the eigenvector for the largest eigenvalue of the whole-genome matrix *R*_*ij*_ . The amplitude of the compartment signal was normalized to make its dispersion unity and its sign was defined as positive for compartment A and negative for compartment B.

### Data of simulated lamina-chromatin association

#### Simulated probability distribution of chromatin residence near the nuclear envelope

We counted the number

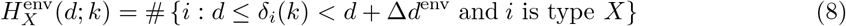

of beads of type X (= A, B, u, or no) lying within a range of distance [*d*, *d* + Δ*d*^env^) from the nuclear envelope of the *k*th cell. Here, beads of type A, B, and u are chromatin beads and beads of type no are nucleolar beads. #{…} is the number of elements in the set {…} and *δ*_*i*_(*k*) is the shortest distance of the *i*th bead from the nuclear envelope in the *k*th cell at the 700,000th step. We collected these numbers from 10 cells and calculated the sum 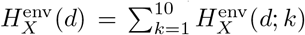. Then, we normalized the frequency 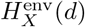 at each distance *d* to get the distribution 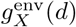 of each type of beads:

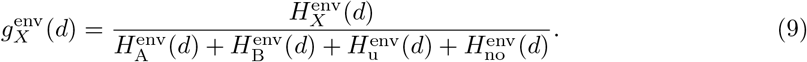

For small enough *d*, 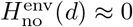 in GM12878, whereas 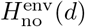 is non-negligible in IMR90 because of the flat shape of the ellipsoidal IMR90 nucleus. Because we modelled each 100-kb chromatin segment as a sphere with soft-core repulsions, 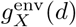 exhibits an oscillatory pattern with a period corresponding to the diameter of the sphere. The overall trend of the distribution is the essential prediction and the oscillation is an artifact of this coarse-graining in the simulation. Hence, we calibrated the bin width to Δ*d*^env^ = 0.083 *μ*m so that the oscillatory component is smoothed out. The thus obtained 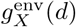 is shown in Fig. 7A.

#### Simulated frequency of lamina-chromatin association

Instantaneous lamina-chromatin contact *λ*_*i*_(*t*; *k*) at the *i*th bead of the *k*th cell was sampled at time *t* as

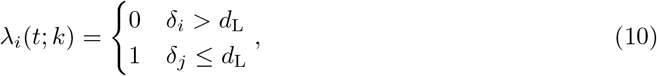

where *δ*_*i*_ is the distance between the *i*th chromatin bead and the nuclear envelope and *d*_L_ = 0.12 *μ*m = *d*_c_/2 is the threshold distance. We sampled *λ*_*i*_(*t*; *k*) in every 1,000 steps from the 500,000th to 700,000th step. Writing this set of sampling instances as *T*_L_, the population lamina-chromatin contact was calculated from *N*_cell_ = 200 simulation runs as

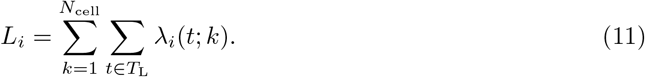

The profile {*L*_*i*_} was normalized to its genome-wide average and log-transformed:

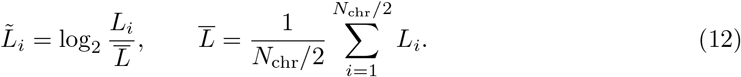

In Figs. 7B and 7C, the resulting profile 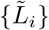 is compared with the experimental data.

#### Dynamic chromatin dissociation from the nuclear envelope

We analyzed a spatiotemporal chromatin distribution near the nuclear envelope in an example single simulated cell. We sampled chromatin configuration in every 1,000 steps from the 450,000th to 700,000th step. We write this set of time instances as *T*_DL_. Let *t*_*i*_ be the time passed after the bead *i* made a first contact to the nuclear envelope;

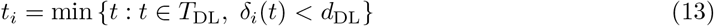

where *δ*_*i*_(*t*) is the distance between the bead *i* and the nuclear envelope, and *d*_DL_ = 0.15 *μ*m is the threshold distance. We assumed *t*_*i*_ = ∞ if the *i*th bead did not approach to the nuclear envelope within the simulated duration; *t*_*i*_ < ∞ means that the *i*th bead was in contact with the lamina at time *t_i_* and stayed at the lamina or diffused away from the lamina thereafter. We counted the number

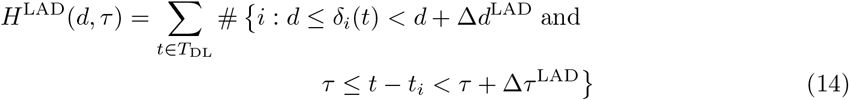

of chromatin beads lying within a range of distance [*d*, *d* + Δ*d*^LAD^) from the nuclear envelope in the time interval [*τ*, *τ* + Δ*τ*^LAD^). Here, *δ*_*i*_(*t*) is the shortest distance of the *i*th bead from the nuclear envelope at time *t*. We also computed the spatial distribution of all the chromatin beads in all the time steps as a reference:

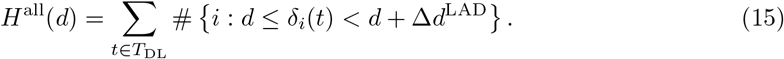

Then, we normalized the frequency by the reference frequency to get the normalized spatiotemporal distribution of chromatin once attached to the lamina at *τ* = 0:

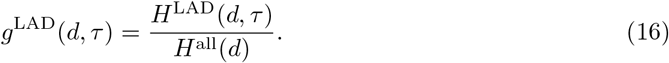

We used the spatial bin size Δ*d*^LAD^ = 0.022 *μ*m and the temporal bin size Δ*τ*^LAD^ = 0.01*τ*_0_ (i.e., 1,000 steps) to show *g*^LAD^(*d, τ*) in Fig. 8B.

### Data of simulated nucleoli-chromatin association

#### Simulated frequency of nucleoli-chromatin association

Nucleoli were represented as droplets formed by the assemblies of nucleolar beads in the present model. The frequency of nucleoli-chromatin association was calculated by contact counts between chromatin beads *i* ∈ *P*_chr_ and nucleolar beads *j* ∈ *P*_no_ in the model. Instantaneous nucleoli-chromatin contact was calculated in the same way as the instantaneous chromatin-chromatin contact of *m*_*ij*_ and summed over time and cells. From the obtained contact 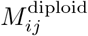 with *i* ∈ *P*_chr_ and *j* ∈ *P*_no_, the population nucleoli-chromatin contact profile {*O*_*i*_} was derived as

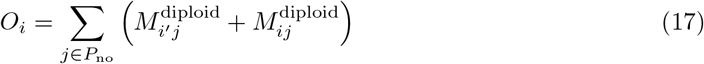

where the bead *i*′ is a homologous copy of the chromatin bead *i*. Then, the profile {*O*_*i*_} was normalized to its genome-wide average and log-transformed:

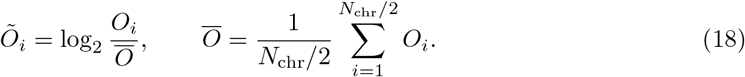

In Figs. 7D and 7E, the resulting profile 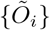 is compared with the experimental data.

## Acknowledgment

We are grateful to Dr. Kazuhiro Maeshima for crtical reading of the manuscript. This work was supprted by the CREST Grant JPMJCR15G2 of Japan Science and Technology Agency; the Riken Pioneering Project; the KAKENHI Grants, 19H01860, 19H05258, 20H05530, and 21H00248, of Japan Society for the Promotion of Science.

