## Supplementary material for "Generation of dynamic three-dimensional genome structure through phase separation of chromatin": SI Appendix, SI Text and SI Figures

### Supplementary Information for Generation of dynamic three-dimensional genome structure through phase separation of chromatin

Shin Fujishiro and Masaki Sasai

February 15, 2022

**8 Captions for Supplementary Figures and Movies** **25**

### 1 Experimental data

#### 1.1 Experimental Hi-C data

For analyzing IMR90, GM12878, and CH12-LX cells, we used the experimental data [1] deposited in the Gene Expression Omnibus (GEO) database [2] with the accession number GSE63525. The contact matrix was normalized with the Knight and Ruiz (KR) method [3] when analyzed with the correlation coefficient between vectors in the contact matrix [4] as shown in Fig. 5B and to calculate the compartment signal (PC1) in Figs. 1 and 6.

#### 1.2 Experimentally observed frequency of lamina-chromatin association

The Lamin-B1 ChIP-seq data for ER:Ras expressing IMR90 cells [5] were obtained from the GEO database [2] with the accession number GSE49341. FASTQ files for the three LMNB1 ChIP-seq experiments marked as *No 4OHT* and the corresponding three input reads were used for analysis. The reads were mapped to the hg19 reference genome sequence [6] using bwa aln command [7]. Alignments on low-mappability regions were filtered out by intersecting with the complement of the wgEncodeDacMappabilityConsensusExcludable dataset [8] using bedtools [9]. Duplicate mappings were filtered out by samtools rmdup command [10]. Then, the pileup count was computed for each alignment using macs2 callpeak command, and the enrichment signal was calculated at each base by dividing ChIP-seq pileup count by input pileup count [11]. The enrichment signal was averaged over each 100-kb bins and over the three replicates. We denote the $\log_2$ of the average enrichment signal by $\{\tilde{L}_i\}$ where $i$ is the index number of the 100-kb bins over the genome. We used the thus derived $\{\tilde{L}_i\}$ for comparison with the simulation results in Fig. 7 and Fig. S12.

#### 1.3 Experimentally observed frequency of nucleoli-chromatin association

The tiling array data for nucleolus-associated chromosomal domains in proliferating IMR90 cells [12] was obtained from the GEO database [2] with the accession number GSE78043. The RATIO_CORRECTED column in the dataset, which is the normalized $\log_2$ enrichment of nucleolus association, was extracted and averaged over each 100-kb bin and over the two replicates. We denote the result by $\{\tilde{O}_i\}$ where $i$ is the index number of the 100-kb bin running over the genome, and used it for comparison with the simulation results in Fig. 7 and Fig. S13.

### 2 Neighboring region contact index (NCI)

#### 2.1 NCI with 50-kb and 100-kb resolution

The experimental Hi-C data analyzed in the present study has the 1-kb resolution [1]. Here, in order to analyze the 50-100 kb scale physical features of chromatin, we summarized these 1-kb loci into bins of 50-kb regions. We define the total contact counts between the $i$th 50-kb region and the $j$th 50-kb region as

$$C_{i,j} = \sum_{k \in i\text{th region}} \sum_{l \in j\text{th region}} m_{kl}, \tag{S1}$$

where $k$ and $l$ label loci with the 1-kb resolution, and $m_{kl}$ is the contact counts between the $k$th and $l$th loci of the sequence. We found that $C_{i,i\pm1}$ is an important factor for describing the contact pattern shown in Fig. 1A in the main text. If the bias of the Hi-C data at the $i$th region is represented by a multiplicative factor $b(i)$, the unbiased contact frequency $P_{i,j}$ is defined by $C_{i,j} = b(i)b(j)P_{i,j}$. Then, the effects of the bias are eliminated from $C_{i,i\pm1}$ by defining

*Neighboring Region Contact Index* (NCI) at the $i$th 50-kb region,

$$W_{50\,\mathrm{kb}}(i) = \frac{1}{2}\left[\frac{C_{i,i+1}}{\sqrt{C_{i,i}C_{i+1,i+1}}} + \frac{C_{i,i-1}}{\sqrt{C_{i,i}C_{i-1,i-1}}}\right] = \frac{1}{2}\left[\frac{P_{i,i+1}}{\sqrt{P_{i,i}P_{i+1,i+1}}} + \frac{P_{i,i-1}}{\sqrt{P_{i,i}P_{i-1,i-1}}}\right]. \quad \text{(S2)}$$

To derive the second line of Eq. S2, $C_{i,i} = b(i)^2 P_{i,i}$ and $C_{i\pm1,i\pm1} = b(i \pm 1)^2 P_{i\pm1,i\pm1}$ were used. In Figs. 1B and 1C, NCI was plotted with a 100-kb resolution, using

$$W_{100\,\mathrm{kb}}(k) = \frac{1}{2}\left[W_{50\,\mathrm{kb}}(2k) + W_{50\,\mathrm{kb}}(2k+1)\right] \quad \text{(S3)}$$

for NCI of the $k$th 100-kb chromatin region.

#### 2.2 The ABu annotation of the genome

The genome was divided into bins of 100 kb. The NCI was smoothed over 500 kb around the $k$th 100-kb region as

$$w(k) = \frac{1}{5}\sum_{l=-2}^{2} W_{100\,\mathrm{kb}}(k+l). \quad \text{(S4)}$$

Then, from the distribution of $w(k)$, the $Z$ score of NCI, $Z_w(k)$, was calculated using the mean $\langle w \rangle$ and the deviation $\sigma_w$ as $Z_w(k) = (w(k) - \langle w \rangle)/\sigma_w$, where $\langle w \rangle$ and $\sigma_w$ were calculated by taking average over 22 chromosomes excluding the X chromosome. For these 22 chromosomes, a pair of homologous chromosomes were assumed to have the same NCI and therefore the same $Z_w(k)$. Then, each 100-kb region was classified into type A for $Z_w \geq 0.3$, type B for $Z_w \leq -0.3$, and type u for $-0.3 < Z_w < 0.3$. DNA regions having repetitive sequence (e.g., pericentromeric regions), for which the Hi-C data are not available, were regarded as type-B.

The active and inactive X chromosomes should have different NCI values. However, we cannot directly calculate their difference from the non-allelic Hi-C data. Here, we approximated the NCI to be small everywhere in the inactive X chromosome; therefore, the entire inactive X chromosome was approximated to be type B. With this small NCI approximation for the inactive X chromosome, the variation of the non-allelic NCI should reflect the variation of NCI in the active X chromosome. Therefore, we defined $\langle w \rangle$ and $\sigma_w$ from the non-allelic Hi-C data of the X chromosome to calculate $Z_w$ in the X chromosome, and we used this $Z_w$ to annotate the active X chromosome in the same way as in the other chromosomes.

### 3 Estimating the coarse-grained interactions between 100-kb chromatin regions

We simulated a system of chromatin chains with a 1-kb resolution to estimate the coarse-grained interactions between 100-kb chromatin regions. In this section, the system with 1-kb resolution is referred to as the fine-grained (FG) system and a 100-kb region is referred to as the coarse-grained (CG) region.

#### 3.1 The 1-kb resolution model of chromatin chains

The FG system consists of $N_\mathrm{c}$ copies of chromatin chains, and each chain consists of $N_\mathrm{b}$ beads; $N_\mathrm{FG} = N_\mathrm{c} N_\mathrm{b}$ is the total number of beads in the system. Each bead represents a 1-kb chromatin region and three-dimensional vector $\mathbf{r}_i$ represents the position of the $i$th bead. Each bead is either type A (active, euchromatin-like) or B (inactive, heterochromatin-like), and we considered type-A

(B) chains in which all beads are type-A (B). In our 1-kb resolution model, type-A and type-B chains are distinguished by (1) the one-dimensional features of the loop kinetics and by (2) the presence/absence of attractive interactions between nucleosomes.

###### 3.1.1 The fine-grained potential function

We used the FG interaction potential between 1-kb beads, $U_{\text{kb}}$,

$$U_{\text{kb}} = U_{\text{spring}} + U_{\text{excl}} + U_{\text{nucl-nucl}} + U_{\text{cohesin}}, \tag{S5}$$

where $U_{\text{spring}} = \sum_i u_{\text{spring}}(r_{ii+1})$ connects neighboring beads along the chain with $r_{ii+1} = |\mathbf{r}_i - \mathbf{r}_{i+1}|$ and

$$u_{\text{spring}}(r) = \begin{cases} \frac{1}{2}K_{\text{kb}}(r-\sigma_{\text{kb}})^2 & (r > \sigma_{\text{kb}}) \\ 0 & (r \le \sigma_{\text{kb}}) \end{cases}. \tag{S6}$$

When two 1-kb chromatin regions spatially overlap with each other, they should show soft volume-excluding repulsion. We modelled this effect using a potential, $U_{\text{excl}} = \sum_{i,j} u_{\text{excl}}(r_{ij})$, with $r_{ij} = |\mathbf{r}_i - \mathbf{r}_j|$ and

$$u_{\text{excl}}(r) = \begin{cases} 0 & (r > \sigma_{\text{kb}}) \\ \varepsilon_{\text{kb}}\left(1-\left(\frac{r}{\sigma_{\text{kb}}}\right)^2\right)^3 & (r \le \sigma_{\text{kb}}) \end{cases}. \tag{S7}$$

Here, we did not consider the hard repulsion as topoisomerase II should disentangle the colliding chains. The functional form of Eq. S7 closely resembles to a Gaussian, and we used this form instead of the Gaussian to gain the comutational efficiency.

$U_{\text{nucl-nucl}}$ in Eq. S5 represents the attractive interactions arising from the nucleosome-nucleosome contact. These interactions should involve interactions between positively charged histone tails and acidic patches of nucleosomes, which depend on the configuration of individual nucleosomes. Therefore, unlike the central force such as the Coulomb force between point charges, these interactions depend on the geometric configuration of nucleosomes [13] and are directional [14]. In our 1-kb resolution model, we represented the interactions between nucleosomes with stochastically varying configurations by assuming interactions between stochastically chosen pairs of beads residing near each other. We call the chosen pairs "glued" pairs. Writing the collection of glued pairs as $G(t) = \{(i,j)\}$ at time $t$, we assumed the attractive interactions, $U_{\text{nucl-nucl}} = \sum_{(i,j)\in G(t)} u_{\text{glue}}(r_{ij})$, with

$$u_{\text{glue}}(r) = \begin{cases} 0 & (r > \sigma_{\text{glue}}) \\ -\varepsilon_{\text{glue}}\left(1-\left(\frac{r}{\sigma_{\text{glue}}}\right)^8\right)^3 & (r \le \sigma_{\text{glue}}) \end{cases}. \tag{S8}$$

We set $\varepsilon_{\text{glue}} = 0$ for type-A chains and $\varepsilon_{\text{glue}} > 0$ for type-B chains. A glued contact between nucleosomes with stochastically varying configuration has a lifetime of $\lesssim 100\,\text{ms}$ [15]. When HP1 binds to type-B chains, the glued contact is stabilized, whose lifetime extends to $\sim 500\,\text{ms}$ [15]. We represented the effect of the HP1 binding by reducing the rate of ungluing as explained in the subsection "Glue kinetics" of this SI text.

A cohesin molecule can bundle two sites of the chain to form a loop. When cohesin bundles beads $i$ and $j$, the chromatin chain forms a loop whose boundaries are at $i$ and $j$. We represent this effect with the attractive potential $U_{\text{cohesin}}$ in Eq. S5. Let $L(t) = \{(i,j)\}$ be the collection of pairs of beads residing at the loop boundaries at time $t$. We used the same $u_{\text{spring}}(r)$ as in Eq. S6,

$$U_{\text{cohesin}} = \sum_{(i,j)\in L(t)} u_{\text{spring}}(r_{ij}). \tag{S9}$$

##### 3.1.2 Glue kinetics

We simulated the attractive gluing interactions between nucleosomes by stochastically choosing a pair of 1-kb chromatin beads. We write the collection of glued pairs of beads at time $t$ as $G(t) = \{(i,j)\}$. On each Langevin simulation timestep, $G(t)$ was updated to $G(t+\delta t)$ with the following three steps:

1. (Separation) The attractive nucleosome-nucleosome interactions are short-ranged; therefore, they should turn off if the distance between a pair of nucleosomes exceeds a certain thershold. The distance $r_{ij}$ between each glued pair $(i,j) \in G$ was monitored, and the pair was eliminated from $G$ if $r_{ij} > \sigma_{\text{glue}}$.

2. (Ungluing) Nucleosomal interactions are directional, which can be disrupted with the stochastic fluctuation of nucleosomes. The HP1-mediated interactions are also amenable to stochastic turnover [15]. Hence, each pair $(i,j) \in G$ was eliminated from $G$ with the probability $p_{\text{unglue}} = 1 - \exp(-k_{\text{unglue}}\delta t)$, where $k_{\text{unglue}}$ is the rate constant and $\delta t$ is the size of a Langevin timestep. We assumed that $1/k_{\text{unglue}}$ is the lifetime of the nucleosome configuration keeping the direction of gluing interaction when HP1 is absent. When HP1 can stabilize the glued contacts, we assumed that $1/k_{\text{unglue}}$ is the lifetime of the HP1 bound state on the nucleosome pair.

3. (Gluing) We define a set of glueable pairs: $\tilde{G} = \{(i,j) \notin G : r_{ij} \leq \sigma_{\text{glue}}\}$. Each glueable pair $(i,j) \in \tilde{G}$ was newly added to $G$ with the probability $p_{\text{glue}} = 1 - \exp(-k_{\text{glue}}\delta t)$, where $k_{\text{glue}}$ is the rate constant.

##### 3.1.3 Loop kinetics

A pair $(i,j) \in L(t)$ represents a loop formed between the $i$ and $j$th beads. We simulated kinetics of $(i,j)$ as a sequence of Poisson processes by three steps; (1) unloading, (2) loading, and (3) movement of cohesin, at every time step during the discretized Langevin simulation of the chain motion:

1. (Unloading) In the first step, we simulated the cohesin unloading from chains. When cohesin detaches from either $i$ or $j$, the loop $(i,j)$ is resoleved and eliminated from $L(t)$. We assumed that this unloading takes place at each $(i,j)$ with the probability,

$$p_{\text{off}}(i,j) = 1 - \exp\left(-\gamma_{\text{off}}(i,j)k_{\text{off}}\delta t\right) \tag{S10}$$

   where $k_{\text{off}}$ is a constant, $\gamma_{\text{off}}(i,j) = \min(\gamma_{\text{off}}(i), \gamma_{\text{off}}(j))$, and $0 \leq \gamma_{\text{off}}(i) \leq 1$ represents the tendency of cohesin detachement from the site $i$. The locus $i$ that binds cohesin with some free-energetic prefernce should have the lower detachability $\gamma_{\text{off}}(i)$. We assumed that $\gamma_{\text{off}}(i)$ is small at the CTCF bound sites.

2. (Loading) In the second step, we simulated the cohesin loading onto chains. We considered that a new loop is formed by cohesin loading. We determined the number of loop-forming events in a timestep $\delta t$ by a Poisson-distributed random number $m \sim \text{Poisson}(\lambda)$ with the average $\lambda = N_{\text{FG}}k_{\text{on}}\delta t$. Then, we loaded $m$ new loops at random sites $(i,i) \in L$ where $i \sim$ i.i.d. $U[1, N_{\text{FG}}]$. Each loading move is accepted with the probability $\gamma_{\text{on}}(i)$.

3. (Movement) In the third step, we considered the effect of kinetic sliding of cohesin along the chain. With the cohesin movement, $i$ and $j$ of $(i,j) \in L(t)$ slide along the chromatin chain. There are four moves to consider; (1) the left boundary of the loop at the site $i$ slides in the loop-extruding direction $i' = i - 1$, (2) $i$ slides in the loop-contracting direction $i' = i + 1$, (3) the right boundary at the site $j$ slides in the loop-extruding direction $j' = j + 1$, or (4) $j$ slides in the loop-contracting direction $j' = j - 1$. We write the rate of loop-extruding slide as $k_+$ and the rate of loop-contracting slide as $k_-$. We modified these rates with the

detachability $0 \le \gamma_{\text{off}}(i) \le 1$ of the departing site $i$ and the attachability $0 \le \gamma_{\text{on}}(i') \le 1$ of the landing site $i'$. Then, we accepted a move with the probability:

$$p_{\pm}(i,i') = 1 - \exp\left(-\gamma_{\text{off}}(i)\gamma_{\text{on}}(i')k_{\pm}\delta t\right). \tag{S11}$$

A complex on the chromatin chain such as an enhancer-promoter complex should block the cohesin movement. Such a blocking site $i$ should have the smaller attachability $\gamma_{\text{on}}(i)$. Type-A chains should accomodate multiple comlexes for transcription or duplication, while type-B chains should be rather homogeneous with the absence of such functional complexes. Therefore, we assumed multiple low-attachability sites (cohesin blockers) on a type-A chain and none on a type-B chain. See the "One-dimensional features of the loop kinetics" subsection below for the details. We considered that cohesin promotes loop extrusion through a biased diffusion along the chromatin chain with the diffusion constant $D_{\text{loop}}$ and the biased velocity $v_{\text{loop}}$; we defined the kinetic rates of sliding as $k_+ = D_{\text{loop}} + v_{\text{loop}}$ and $k_- = D_{\text{loop}}$.

Upon collision of two sliding cohesin molecules, we allow loop boundaries to pass through each other to form a z-loop [16] at the rate $k_z$. Let $\omega(i)$ be the occupancy of the site $i$ by loop boundaries. If one or more loop boundaries already lie on the landing site $i'$ with $\omega(i') \ge 1$, we generated the number of z-looping events as $m_z \sim \text{Poisson}(k_z\delta t)$. The move was accepted if $m_z \ge \omega(i')$, and rejected otherwise.

##### 3.1.4 One-dimensional features of the loop kinetics

The CTCF bound sites block the loop extrusion as they bind and trap cohesin molecules. We represented this affinity of the $i$th CTCF-bound site for cohesin as $\gamma_{\text{off}}(i+1) = \gamma_{\text{off}}(i-1) = 0.1$, while $\gamma_{\text{off}} = 1$ at the other sites. We represented the blocking effect of the $i$th CTCF-bound site as $\gamma_{\text{on}}(i) = 0$. The Hi-C data measured upon disruption of DNA methyltransferase activity showed that the heterochromatin-like features deplete the CTCF binding to DNA, leading to the larger topologically associated domains (TADs) in heterochromatin-like regions [17]. Therefore, we assumed that the TAD size is $\sim 500$ kb in type-B chains and $\sim 100$ kb in type-A chains. Complexes formed for transcription or duplication should hinder the cohesin movement. We represented this road-blocking effect at the site $i$ as $\gamma_{\text{on}}(i) = 0.01$. Such functional complexes should be abundant in type-A chains. Therefore, we assumed multiple road blocks in type-A chains while none in type-B chains. These assumptions are summarized in Table 1.

Table 1: One-dimensional features of the loop kinetics

| | Parameters | Sites |
|---|---|---|
| **Type-A chains** | ($N_b = 500$: Chains of 500-kb length) | |
| attachability | $\gamma_{\text{on}}(i) = 0$ | $i = 1, 101, 201, 301, 401, 500$ |
| | $\gamma_{\text{on}}(i) = 0.01$ | $i = 11, 21, 31, \ldots, 491$ |
| | $\gamma_{\text{on}}(i) = 1$ | other sites |
| detachability | $\gamma_{\text{off}}(i) = 0.1$ | $i = 2, 100, 202, 102, 300, 302, 400, 402, 499$ |
| | $\gamma_{\text{off}}(i) = 1$ | other sites |
| **Type-B chains** | ($N_b = 500$: Chains of 500-kb length) | |
| attachability | $\gamma_{\text{on}}(i) = 0$ | $i = 1, 500$ |
| | $\gamma_{\text{on}}(i) = 1$ | other sites |
| detachability | $\gamma_{\text{off}}(i) = 0.1$ | $i = 2, 499$ |
| | $\gamma_{\text{off}}(i) = 1$ | other sites |

Other parameters of the cohesin kinetics are summarized in Table 2. We set the unloading rate $k_{\text{off}}$ of cohesin from the chromatin chain to match $1/k_{\text{off}}$ to the observed mean-residence time of cohesin on the chromatin chain in the G1 nucleus $\sim$ 20min [18]. The observed number of chromatin-bound cohesin complexes was $\sim 160,000$ in a G1 nucleus of HeLa cells [19], suggesting the density of the chromatin-bound cohesin is $0.01\,\text{kb}^{-1}$ to $0.1\,\text{kb}^{-1}$. Thus, the inferred ratio of the on-rate to the off-rate is $0.01\,\mathrm{kb}^{-1} \lesssim k_{\mathrm{on}}/k_{\mathrm{off}} \lesssim 0.1\,\mathrm{kb}^{-1}$, and we used $k_{\mathrm{on}} = k_{\mathrm{off}} \times 0.05\,\mathrm{kb}^{-1}$ in Fig. 2. The observed diffusion coefficient of cohesin along dsDNA *in vitro* was $\sim 2\,\mu\mathrm{m}^2/\mathrm{s}$ [20]. We assumed that the diffusion constant $D_{\mathrm{loop}}$ of cohesin along the chromatin chain *in vivo* is reduced by $\sim 1/1000$ from the value for dsDNA with various chromatin-bound obstacles including nucleosomes as $D_{\mathrm{loop}} \sim 0.002\,\mu\mathrm{m}^2/\mathrm{s} \sim 0.12\,\mu\mathrm{m}^2/\mathrm{min}$. We should note that this is comparable to the diffusion constant of the chromatin movement $D_{\mathrm{kb}}$ as discussed below. Using the ratio of (spatial distance)/(sequential distance) $\sim 25\,\mathrm{nm/kb}$, we have $D_{\mathrm{loop}} \sim 200\,\mathrm{kb}^2/\mathrm{min}$.

The net velocity $v_{\mathrm{loop}}$ of the cohesin movement *in vivo* has not yet been identified quantitatively. The relatively large value of $\sim 577\,\mathrm{bp/s} \sim 35\,\mathrm{kb/min}$ was assumed in a theoretical modeling [21] and the value $\sim 0.4\,\mathrm{kb/s} \sim 25\,\mathrm{kb/min}$ was observed in the *in vitro* measurement of the cohesin movement along dsDNA [22]. However, the *in vitro* motion assay [22] and the structural analyses [16] suggested the non-topological binding of cohesin to the chromatin chain during the loop-extruding process, which may lead to weak persistence of the movement and hence the smaller net velocity of cohesin along the chromatin chain [23]. The hypothesis of purely diffusive movement with $v_{\mathrm{loop}} = 0$ was also proposed [24, 25]. Here, we used the value similar to the velocity of RNA Polymerase $\sim 1\,\mathrm{kb/min}$ [26]. The rate $k_{\mathrm{z}}$ of z-looping, or the crossing rate of loop boundaries across a 1-kb bead, was set to be the same as the rate with diffusive sliding.

Table 2: Parameters of cohesin kinetics

| | Parameters | Remarks |
|---|---|---|
| unloading rate | $k_{\mathrm{off}} = 0.05\,\mathrm{min}^{-1}$ | Ref. [18] |
| loading rate | $k_{\mathrm{on}} = 0.05 k_{\mathrm{off}}$ | Ref. [19] |
| symmetric diffusion constant | $D_{\mathrm{loop}} = 200\,\mathrm{kb}^2/\mathrm{min}$ | Ref. [20] |
| biased velocity | $v_{\mathrm{loop}} = 2\,\mathrm{kb/min}$ | $\sim$ velocity of RNA Polymerase |
| z-looping rate | $k_{\mathrm{z}} = 200\,\mathrm{min}^{-1}$ | comparable to the diffusive sliding |

###### 3.1.5 Parameters of bead-and-spring chains

Size of a 1-kb chromatin bead should be $\sigma_{\mathrm{kb}} \simeq 20$–$40\,\mathrm{nm}$. We assumed the repulsion between beads is strong enough to prevent the frequent chain crossing as $\varepsilon_{\mathrm{kb}} \simeq 10 k_{\mathrm{B}}T$. We determined the spring constant $K_{\mathrm{kb}}$ to connect neighboring beads to make the bond fluctuations less than the average bond length as $\Delta r \sim \sqrt{k_{\mathrm{B}}T/K_{\mathrm{kb}}} < \sigma_{\mathrm{kb}}$. We estimated the diffusion coefficient $D_{\mathrm{kb}}$ of each 1 kb bead from the live-cell microscope data [27]. The motion of an EGFP-tagged *lacO* array inserted on a mammalian chromosome followed a short-term confined diffusion *in vivo*, where the fast diffusion constant under confinement was $D_{\mathrm{f}} = 3.13 \times 10^{-3}\,\mu\mathrm{m}^2/\mathrm{s}$ and the slow diffusion constant of the motion going out of the confinement was $D_{\mathrm{s}} = 2.4 \times 10^{-4}\,\mu\mathrm{m}^2/\mathrm{s}$. We assumed that the fast one $D_{\mathrm{f}}$ is the intrinsic diffusion constant, which is not affected by the many-body hindrance in the chromatin environment; therefore, we set $D_{\mathrm{kb}} \simeq D_{\mathrm{f}} \sim 0.2\,\mu\mathrm{m}^2/\mathrm{min}$.

The experimentally observed lifetime of a tetra-nucleosome conformation *in vitro* is $\lesssim 100\,\mathrm{ms}$, and it extends to $\sim 500\,\mathrm{ms}$ when HP1 binds to chromatin [15]. We assumed that the lifetime of a glued pair of nucleosomes across chains is in a similar order. Therefore, we set $k_{\mathrm{unglue}} = 1000\,\mathrm{min}^{-1}$ (time constant 60 ms) for the HP1-free heterochromatin simulations and $k_{\mathrm{unglue}} = 100\,\mathrm{min}^{-1}$ (time constant 600 ms) for the heterochromatin simulations with HP1. We set the rate of forming a nucleosome-nucleosome gluing interaction $k_{\mathrm{glue}}$ is large enough to condense chromatin in the simulation. These values are summarized in Table 3.

###### 3.1.6 Simulations

We simulated the FG system by numerically integrating the Langevin molecular dynamics,

$$\frac{d\mathbf{r}_i}{dt} = -\mu_{\mathrm{kb}} \frac{\partial U_{\mathrm{kb}}}{\partial \mathbf{r}_i} + \sqrt{2D_{\mathrm{kb}}}\boldsymbol{\xi}_i(t), \tag{S12}$$

Table 3: Parameters of bead-and-spring chains

| | Parameters | Remarks |
|---|---|---|
| bead size | $\sigma_{\mathrm{kb}} = 0.025\,\mu\mathrm{m}$ | in the range of $\approx$ 20–40 nm |
| bead-bead repulsion | $\varepsilon_{\mathrm{kb}} = 10\,k_{\mathrm{B}}T$ | infrequent chain crossing |
| bond spring constant | $K_{\mathrm{kb}} = 6400\,k_{\mathrm{B}}T/\mu\mathrm{m}^2$ | fluctuations less than the average bond length |
| diffusion constant | $D_{\mathrm{kb}} = 0.2\,\mu\mathrm{m}^2/\mathrm{min}$ | Ref. [27] |
| range of attraction | $\sigma_{\mathrm{glue}} = 1.6\sigma_{\mathrm{kb}}$ | attraction range comparable to the bead size |
| rate of gluing | $k_{\mathrm{glue}} = 3000\,\mathrm{min}^{-1}$ | sufficeint for the in-vitro chromatin condensation Ref. [28] |
| bead-bead attraction | $\varepsilon_{\mathrm{glue}} = 0$ (type-A chains) | no in-vitro chromatin condensation Ref. [28] |
| | $\varepsilon_{\mathrm{glue}} = k_{\mathrm{B}}T$ (type-B chains) | sufficeint for the in-vitro chromatin condensation Ref. [28] |
| rate of ungluing | $k_{\mathrm{unglue}} = 100\,\mathrm{min}^{-1}$ (with HP1 binding)<br>$k_{\mathrm{unglue}} = 1000\,\mathrm{min}^{-1}$, (without HP1 binding) | Ref. [15] |

where $\mu_{\mathrm{kb}} = D_{\mathrm{kb}}/k_{\mathrm{B}}T$ is the mobility of a bead and $\boldsymbol{\xi}_i(t)$ is a Gaussian white noise vector satisfying $\langle \xi_{i\nu}(t_1)\xi_{j\nu'}(t_2)\rangle = \delta_{ij}\delta_{\nu\nu'}\delta(t_1 - t_2)$ and the labels $\nu$ and $\nu'$ distinguish the $xyz$ component of the vector. Eq. S12 was discretized with the timestep of $\delta t = 10^{-4}$ min, which satisfies $\delta t \mu_{\mathrm{kb}} K_{\mathrm{kb}} < 1$, insuring the numerical stability of the integration. The trajectories were followed up to $3 \times 10^6$ steps = 300 min, which was sufficiently long for the slow cohesin kinetics ($1/k_{\mathrm{off}} = 20$ min) to reach the stationary state.

The size of a simulation box and the number of chains $N_{\mathrm{c}}$ for given densities are summarized in Supplementary Table 4. The numbers were determined for maximizing the quality of the sampled pair correlation function $g(r)$ within the available computational resources. The chains were initialized as random walk paths of a step size $\sigma_{\mathrm{kb}}$ starting at random positions, and simulated under the periodic boundary condition. If we assume $\sigma_{\mathrm{kb}} = 35$ nm, the lowest and the highest density in the table translate to $\rho = 0.47\,\mathrm{Mb}/\mu\mathrm{m}^3$ and $\rho = 9.33\,\mathrm{Mb}/\mu\mathrm{m}^3$, respectively. The latter estimation corresponds to 6 Gb of genomic chromosomes in a nucleus of volume $643\,\mu\mathrm{m}^3$.

Table 4: Definitions of fine-grained simulation system

| Reduced number density $\rho\sigma_{\mathrm{kb}}^3$ | Box size ($\mu$m) | Number of chains $N_{\mathrm{c}}$ |
|---|---|---|
| 0.02 | 1.462 | 8 |
| 0.10 | 1.328 | 30 |
| 0.20 | 1.054 | 30 |
| 0.30 | 0.9210 | 30 |
| 0.40 | 0.8368 | 30 |

#### 3.2 The PRISM theory for coarse graining

We derived the coarse-grained potential function, $u_{\mathrm{CG}}(r)$, between 100 kb chromatin regions from the simulated results of the FG system explained in the last subsection. We carried out this coarse-graining procedure by using the polymer reference interaction site model (PRISM) theory [29]. In the simulated system, we have $N_{\mathrm{CG}} = \left(\frac{1\,\mathrm{kb}}{100\,\mathrm{kb}}\right) N_{\mathrm{FG}} = 0.01 N_{\mathrm{FG}}$ segments of 100 kb size and each chain has $N_{b,\mathrm{CG}} = 0.01 N_b$ segments. Neglecting the end effect of polymer chains as in the usual treatment in the PRISM theory, we can regard that the system has $N_{\mathrm{CG}}$ equivalent segments. Let $\rho$ be the bulk density of segments. Then, the intra-chain structure factor $\omega(k)$ and the inter-chain segment-segment pair correlation function $g(r)$ become

$$\omega(k) = 1 + N_{\mathrm{CG}} \left\langle \frac{\sin(kr_{ij})}{kr_{ij}} \right\rangle_{\mathrm{intra}}, \tag{S13}$$

$$g(r) = \frac{1}{\rho} \left\langle \delta(r - r_{ij}) \right\rangle_{\mathrm{inter}}, \tag{S14}$$

where $\langle \cdots \rangle_{\mathrm{intra}}$ represents the average over all realizations of intra-chain segment-segment distances $r_{ij}$, and $\langle \cdots \rangle_{\mathrm{inter}}$ is the average over all realizations of inter-chain segment-segment distances $r_{ij}$. We should note that the periodic images are not included in calculating $\langle \cdots \rangle_{\mathrm{intra}}$ because the segments appearing in the periodic images are on different chains.

In the Fourier $k$-space, the single-species PRISM equation relates the inter-chain total correlation function $h(r) = g(r) - 1$ and the inter-chain direct correlation function c(r) as

$$\hat{h}(k) = \omega(k)\hat{c}(k)\left(\omega(k) + \rho\hat{h}(k)\right), \tag{S15}$$

where the hatted quantities are the three-dimensional Fourier transforms of the unhatted ones. When we have the data of $\omega(k)$ and $h(r)$ in the FG model, we can derive $c(r)$ using Eq. S15. However, the PRISM equation Eq. S15 raises a technical issue; $h(r)$ derived from simulations has a cutoff at the distance of half of the simulation-box size. The cutoff induces artifactual ripples in the Fourier-transformed total correlation function $\hat{h}(k)$, which leads to an incorrect solution of $c(r)$. Here, we resolved this issue by introducing the cutoff in $c(r)$ as proposed by Bernheardt *et al.* [30]; because $c(r)$ decays much faster to 0 than $h(r)$, we can introduce an explicit cutoff in the PRISM equation as

$$\mathcal{FB}h(r) = \omega(k)\left(\mathcal{FB}c(r)\right)\left(\omega(k) + \rho\mathcal{FB}h(r)\right), \tag{S16}$$

where $\mathcal{B}$ is the zero-extension operator as explained below for the numerical treatment and $\mathcal{F}$ is the Fourier transformation operator. From Eq. S16, we have

$$h(r) = \mathcal{B}^{-1}\mathcal{F}^{-1}\left(\frac{\omega(k)^2\mathcal{FB}c(r)}{1 - \rho\omega(k)\mathcal{FB}c(r)}\right), \tag{S17}$$

which we call the PRISM-cut equation. Then, with a closure relation, we obtain the coarse-grained potential function, $u_{\mathrm{CG}}(r)$, from $h(r)$ and $c(r)$. When we adopt the Perkus-Yevick closure relation, we have

$$u_{\mathrm{CG}}(r) = -k_{\mathrm{B}}T\log\left(\frac{c(r)}{h(r) - c(r) + 1} + 1\right). \tag{S18}$$

Using $h(r)$ and $\omega(k)$ obtained from the Langevin simulations of the FG system, we numerically solved Eq. S17 using an iterative root-finding method proposed by Bernheardt [30]. We first discretized the distance coordinate $r$ with uniform intervals $\Delta r$;

$$r_n = (n - 0.5)\Delta r, \quad \text{with } n = 1, \ldots, N. \tag{S19}$$

The wavenumber coordinate $k$ is discretized accordingly,

$$k_n = (n - 0.5)\Delta k, \quad \text{with } n = 1, \ldots, N \text{ and } \Delta k = 2\pi/(N\Delta r). \tag{S20}$$

Let $M$ be the discretized index for the cutoff distance $r_M$ of the simulation-derived pair correlation functions. Then, the correlation functions are implemented as $M$-dimensional vectors as $h, c, \omega \in \mathbf{R}^M$ with $h_n = h(r_n)$, $c_n = c(r_n)$, and $\omega_n = \omega(k_n)$. The operators are implemented as matrices,

$$\mathcal{B} \in \mathbf{R}^{N\times M}, \quad \mathcal{B}_{nm} = \delta_{nm}, \tag{S21}$$

$$\mathcal{F} \in \mathbf{R}^{N\times N}, \quad \mathcal{F}_{nm} = \frac{\sin(k_n r_m)}{k_n r_m} 4\pi r_m^2 \Delta r. \tag{S22}$$

The PRISM-cut equation now gives the total correlation $h^\dagger$ as a vector-valued function of $c$;

$$h^\dagger = \mathcal{B}^t \mathcal{F}^{-1} \left( \frac{\omega \odot \omega \odot \mathcal{F}\mathcal{B}c}{1 - \rho\omega \odot \mathcal{F}\mathcal{B}c} \right), \tag{S23}$$

where $\mathcal{B}^t$ is the transpose of the $\mathcal{B}$, $\odot$ is the element-wise vector multiplication operator, and the division is also calculated element-wise. The Jacobian of the function $h^\dagger$ is an $M \times M$ matrix given by

$$J(c) = \frac{\delta h^\dagger}{\delta c} = \mathcal{B}^t \mathcal{F}^{-1} \left( \frac{\omega}{1 - \rho\omega \odot \mathcal{F}\mathcal{B}c} \right)^{\odot 2} \mathcal{F}\mathcal{B}, \tag{S24}$$

where the $\odot$-exponentiation is calculated element-wise. We then find a root of the equation $h^\dagger(c) - h = 0$ by the Newton-Raphson iterative procedure. On each iteration, the direct correlation $c^{(n)}$ at the $n$th iteration was updated as $c^{(n+1)} = c^{(n)} - \Delta c^{(n)}$, where $\Delta c^{(n)}$ is a solution (given by the NumPy lstsq function) of the linear equation

$$J(c^{(n)})\Delta c^{(n)} = h^\dagger(c^{(n)}) - h. \tag{S25}$$

We used the initial value $c^{(0)} = 0$ and the convergence criterion, $\max|\Delta c^{(n)}| < 10^{-3}$. From the converged $c(r)$, we calculated the effective coarse-grained potential $u_{\mathrm{CG}}(r)$ using the closure Eq. S18, which we plotted in Fig. 2 with a smoothing procedure described below.

The coarse-grained potential $u_{\mathrm{CG}}(r)$ suffers from the high-frequency noise arising from the sampling noise of the pair correlation function $g(r)$; see Fig. S1. To smooth out the noise, we applied a Gaussian filter to the potential function. Let $\hat{u}_{\mathrm{CG}}(k) = \mathcal{F}u_{\mathrm{CG}}$ be the three-dimensional Fourier transformation of $u_{\mathrm{CG}}(r)$. Then, we multiplied a Gaussian function in the Fourier $k$-space

$$\hat{u}_{\mathrm{CG,smooth}}(k) = \exp\left( -\frac{k^2 w^2}{2} \right) \hat{u}_{\mathrm{CG}}(k) \tag{S26}$$

to suppress high-frequency component. Then, we transformed back the result $u_{\mathrm{CG,smooth}}(r) = \mathcal{F}^{-1}u_{\mathrm{CG,smooth}}$ to obtain a smoothed potential. This is equivalent to three-dimensional convolution of a Gaussian kernel $\left(2\pi w^2\right)^{-3/2} \exp\left(-r^2/(2w^2)\right)$; we used the radius of a bead as the size of the Gaussian kernel $w = \sigma_{\mathrm{kb}}/2$.

Finally, we should note about the density parameter $\rho$. The number density $\rho$ of segments quantifies the amount of indirect inter-chain correlations in the system. In an infinitely large system, we may use the bulk density of segments for evaluating $\rho$. However, in a finite system, the density needs to be corrected to exclude the segments on the directly correlated pair of chains. Therefore, when a chain consists of $N_{b,\mathrm{CG}}$ segments and there are $N_{\mathrm{CG}}$ segments in a system of volume $V$, we used the corrected density

$$\rho = \frac{N_{\mathrm{CG}} - 2N_{b,\mathrm{CG}}}{V} \tag{S27}$$

when solving the PRISM-cut equation. We used the bulk density for estimating inter-chain RDF $g(r)$.

### 4 Simulation of anatelophase genome

We simulated the anatelophase human genome to prepare a suitable configuration of chromosomes at the entry to interphase. In anatelophase, chromosomes are first segregated by microtubule contraction (anaphase), then they relax within the nascent nuclear envelope (telophase). Throughout these two stages in the simulation, the 46 human chromosomes were coarse-grained into 46 model homopolymers. Each model polymer is a beads-on-a-string chain with a single bead representing a 10-Mb chromatin region; the resulting system consists of $N_{\mathrm{anatelo}} = 632$ beads. We denote by $\boldsymbol{r}_i$ the 3D coordinate vector of the $i$th bead.

The anatelophase simulation was critical to reproducing the experimentally observed long-range contact frequency between the p-arm and the q-arm of each interphase chromosome. Through the anaphase simulation, the chromosome gained a V-shaped conformation due to the force applied to the kinetochore. Effects of these V-shaped conformations of 46 chromosomes remained in interphase leading to the long-range, inter-arm chromatin contacts.

#### 4.1 Interactions among beads

We assumed the repulsive interactions between overlapping beads and the harmonic bonds along each chain. The potential energy of the chains was given by

$$U_{\mathrm{anatel,chains}} = \sum_{1\le i<j\le N_{\mathrm{anatel}}} u_{\mathrm{anatel,rep}}(r_{ij}) + \sum_{(i,j)\in B_{\mathrm{anatel}}} u_{\mathrm{anatel,bond}}(r_{ij}) \tag{S28}$$

where $r_{ij} = |\boldsymbol{r}_i - \boldsymbol{r}_j|$ and $B_{\mathrm{anatel}}$ is a set of all the pairs of bonded beads. $u_{\mathrm{anatel,rep}}(r_{ij})$ is a pairwise repulsive potential,

$$u_{\mathrm{anatel,rep}}(r) = \begin{cases} 0 & r > \sigma_{\mathrm{anatel}} \\ \varepsilon_{\mathrm{anatel}} \left(1 - \left(\frac{r}{\sigma_{\mathrm{anatel}}}\right)^2\right)^3 & r \le \sigma_{\mathrm{anatel}}, \end{cases} \tag{S29}$$

and $u_{\mathrm{anatel,bond}}(r_{ij})$ is a harmonic potential,

$$u_{\mathrm{anatel,bond}}(r) = \frac{1}{2} K_{\mathrm{anatel}} \left(r - \sigma_{\mathrm{anatel}}\right)^2 . \tag{S30}$$

We used the bead diameter $\sigma_{\mathrm{anatel}} = 0.3\,\mu\mathrm{m}$, the repulsion strength $\varepsilon_{\mathrm{anatel}} = 2k_{\mathrm{B}}T$, and the bond strength $K_{\mathrm{anatel}} = 1000k_{\mathrm{B}}T/\mu\mathrm{m}^2$ at temperature $T$ with the Boltzmann constant $k_{\mathrm{B}}$.

#### 4.2 Initial random positioning of rods

We started each anatelophase simulation run by choosing the initial configuration of each chain to be a randomly directed straight rod; writing the bead number of the start and end of each chain as $s_k$ and $e_k$ with $k = 1, \ldots, 46$, the initial coordinate of the $i$th bead on the $k$th chain is

$$\boldsymbol{r}_i = \boldsymbol{c}_k + \left(i - \frac{s_k + e_k}{2}\right) \sigma_{\mathrm{anatel}} \boldsymbol{w}_k \qquad (s_k \le i \le e_k) \tag{S31}$$

where $\boldsymbol{c}_k$ is a normally distributed random point around the point $(0\,\mu\mathrm{m}, 5\,\mu\mathrm{m}, 0\,\mu\mathrm{m})$ with the standard deviation $1\,\mu\mathrm{m}$, and $\boldsymbol{w}_k$ is a randomly directed unit vector; the $k$th rod centering at the point $\boldsymbol{c}_k$ was directed towards $\boldsymbol{w}_k$. From this initial positioning, the anaphase simulation was started by dragging the kinetochore-attached beads of chains toward the point $(0\,\mu\mathrm{m}, 0\,\mu\mathrm{m}, 0\,\mu\mathrm{m})$.

#### 4.3 Anaphase (dragging) simulation

The region annotated as *acen* in the UCSC hg19 cytoBand dataset [31] was assigned as the centromeric region of each chromosome. Then, we identified the three consecutive beads at the center of the centromeric region as the kinetochore-attached beads. Let $P_{\mathrm{K}}$ be the set of all the beads with kinetochore attachment. We assumed a harmonic potential $U_{\mathrm{mt}}$ representing the effects of microtubule contraction on the beads in $P_{\mathrm{K}}$ as

$$U_{\mathrm{mt}} = \sum_{i\in P_{\mathrm{K}}} \frac{1}{2} K_{\mathrm{mt}} \boldsymbol{r}_i^2 . \tag{S32}$$

Here, the spring constant was chosen to be $K_{\mathrm{mt}} = 1000k_{\mathrm{B}}T/\mu\mathrm{m}^2$ to prevent chains from being apart. Thus, the total potential energy used in the anaphase simulation is

$$U_{\mathrm{ana}} = U_{\mathrm{anatel,chains}} + U_{\mathrm{mt}} . \tag{S33}$$

With this potential energy, we simulated the Brownian dynamics of the beads following the over-damped Langevin equation:

$$\frac{d\boldsymbol{r}_i}{dt} = -\mu \frac{\partial U_{\mathrm{ana}}}{\partial \boldsymbol{r}_i} + \sqrt{2\mu k_{\mathrm{B}} T}\boldsymbol{\xi}_i(t), \tag{S34}$$

where $\boldsymbol{\xi}_i(t)$ is a Gaussian white noise vector satisfying $\langle \xi_{i\nu}(t_1)\xi_{j\nu'}(t_2)\rangle = \delta_{ij}\delta_{\nu\nu'}\delta(t_1 - t_2)$ and the labels $\nu$ and $\nu'$ distinguish the $xyz$ component of the vector. We set the mobility parameter as $\mu = 1\,\mu\mathrm{m}^2/\tau_0 k_{\mathrm{B}}T$ ($\tau_0$ is the unit of simulation time). Starting from the initial positioning described above, we simulated 200,000 steps of the Brownian dynamics with a discretized time step $\delta t = 10^{-4}\tau_0$. The telophase simulation followed thereafter.

#### 4.4 Telophase (packing) simulation

In the telophase simulation, we applied a weak field to beads to confine them in a spherical envelope. The potential energy of the field is

$$U_{\mathrm{env}} = \sum_{i=1}^{N_{\mathrm{anatel}}} u_{\mathrm{env}}(r_i), \tag{S35}$$

with $r_i = |\boldsymbol{r}_i|$ and

$$u_{\mathrm{env}}(r) = \begin{cases} 0 & r \le R_{\mathrm{env}} \\ \frac{1}{2}K_{\mathrm{env}}\left(r - R_{\mathrm{env}}\right)^2 & r > R_{\mathrm{env}}, \end{cases} \tag{S36}$$

where the radius of the envelope is $R_{\mathrm{env}} = 1.2\,\mu\mathrm{m}$ and the strength of the constraint is $K_{\mathrm{env}} = 10 k_{\mathrm{B}}T/\mu\mathrm{m}^2$. Then, using the potential energy,

$$U_{\mathrm{telo}} = U_{\mathrm{anatel,chains}} + U_{\mathrm{env}} + U_{\mathrm{mt}}, \tag{S37}$$

we simulated the Brownian dynamics of the beads for 50,000 steps with the same parameters used in the anaphase simulation.

#### 4.5 Fine-graining with cubic spline curves

A bead in the anatelophase simulation represents a 10-Mb chromatin region. To proceed to the interphase simulation with a 100-kb resolution, we fine-grained the anatelophase chains by 100 fold using not-a-knot cubic spline interpolation curves. We write the 3D curve of the chain $k$ as $\tilde{\boldsymbol{r}}^k = (\tilde{x}^k, \tilde{y}^k, \tilde{z}^k)$, where $\tilde{x}^k = (x_{s_k}, \ldots, x_i, \ldots, x_{e_k})$, $\tilde{y}^k = (y_{s_k}, \ldots, y_i, \ldots, y_{e_k})$, and $\tilde{z}^k = (z_{s_k}, \ldots, z_i, \ldots, z_{e_k})$ with $s_k \le i \le e_k$, and $\boldsymbol{r}_i = (x_i, y_i, z_i)$ is the 3D coordinate vector of the $i$th bead on the chain $k$. We generated a cubic spline function $f_x^k(u)$ so as to make $f_x^k(u)$ trace the $x$ component of the curve $k$, $\tilde{x}^k$, with $u$ being the curve parameter as

$$f_x^k(u_i) = x_i, \qquad u_i = \frac{i - s_k + 0.5}{e_k - s_k + 1} \qquad (s_k \le i \le e_k). \tag{S38}$$

Similarly, we generated cubic spline functions, $f_y^k(u)$ and $f_z^k(u)$, to trace the $y$ and $z$ coordinates. Then, the interpolated coordinates $(\hat{x}_i, \hat{y}_i, \hat{z}_i)$ of fine-grained beads were obtained from $f_x^k(u)$, $f_y^k(u)$, and $f_z^k(u)$. For the $x$ coordinate, for example, we defined

$$\hat{x}_i = f_x^k(\hat{u}_i), \qquad \hat{u}_i = \frac{i - \hat{s}_k + 0.5}{\hat{e}_k - \hat{s}_k + 1} \qquad (\hat{s}_k \le i \le \hat{e}_k). \tag{S39}$$

Here, $\hat{s}_k$ and $\hat{e}_k$ are the bead numbers at the start and the end of the $k$th fine-grained (100-kb resolution) chain, respectively. The $y$ and $z$ coordinates were interpolated similarly.

### 5 Interphase simulation

Interphase genome was modeled with 46 heteropolymers. Each heteropolymer is a beads-on-a-string chain with a single bead representing a 100-kb chromatin region. The p-arms of chromosome 13, 14, 15, 21 and 22 were defined to be rDNA regions, and we regarded one copy in each homologous rDNA pair as active and the other copy as inactive. We attached extra two particles to each bead in the active rDNA regions to represent the volume occupied by the nucleolar mass. We call these particles nucleolar beads. The system has $N_{\mathrm{chr}} = 60752$ chromatin beads representing 100-kb chromatin regions and $N_{\mathrm{no}} = 1426$ nucleolar beads amounting to $N = N_{\mathrm{chr}} + N_{\mathrm{no}} = 62178$ beads in total. The numbering of these beads is summarized in the Supplementary Table 5.

In the interphase simulation, we confined the genome in a spherical (GM12878) or ellipsoidal (IMR90) container, which models the nuclear envelope. In this section, we denote by $\boldsymbol{r}_i$ the coordinate vector of the $i$th bead ($i = 1, \ldots, N$). For the mathematical convenience, we define factors $\alpha_i$ and $\beta_i$ to distinguish the type of beads as

$$\alpha_i = \begin{cases} 1 & \text{bead } i \text{ is type-A chromatin} \\ 0.5 & \text{type-u chromatin} \\ 0 & \text{type-B chromatin} \end{cases}, \qquad \beta_i = \begin{cases} 0 & \text{bead } i \text{ is type-A chromatin} \\ 0.5 & \text{type-u chromatin} \\ 1 & \text{type-B chromatin} \end{cases}. \tag{S40}$$

Table 5: Numbering of the beads in the interphase simulations*

| Entity | Start | End | Entity | Start | End | Entity | Start | End |
|---|---|---|---|---|---|---|---|---|
| chr1:a | 1 | 2493 | chr17:a | 25012 | 25823 | chr10:b | 47186 | 48541 |
| chr2:a | 2494 | 4925 | chr18:a | 25824 | 26604 | chr11:b | 48542 | 49892 |
| chr3:a | 4926 | 6906 | chr19:a | 26605 | 27196 | chr12:b | 49893 | 51231 |
| chr4:a | 6907 | 8818 | chr20:a | 27197 | 27827 | chr13:b | 51232 | 52383 |
| chr5:a | 8819 | 10628 | chr21:a | 27828 | 28309 | chr14:b | 52384 | 53457 |
| chr6:a | 10629 | 12340 | chr22:a | 28310 | 28823 | chr15:b | 53458 | 54483 |
| chr7:a | 12341 | 13932 | chrX:a | 28824 | 30376 | chr16:b | 54484 | 55387 |
| chr8:a | 13933 | 15396 | chr1:b | 30377 | 32869 | chr17:b | 55388 | 56199 |
| chr9:a | 15397 | 16809 | chr2:b | 32870 | 35301 | chr18:b | 56200 | 56980 |
| chr10:a | 16810 | 18165 | chr3:b | 35302 | 37282 | chr19:b | 56981 | 57572 |
| chr11:a | 18166 | 19516 | chr4:b | 37283 | 39194 | chr20:b | 57573 | 58203 |
| chr12:a | 19517 | 20855 | chr5:b | 39195 | 41004 | chr21:b | 58204 | 58685 |
| chr13:a | 20856 | 22007 | chr6:b | 41005 | 42716 | chr22:b | 58686 | 59199 |
| chr14:a | 22008 | 23081 | chr7:b | 42717 | 44308 | chrX:b | 59200 | 60752 |
| chr15:a | 23082 | 24107 | chr8:b | 44309 | 45772 | nucleoli | 60753 | 62178 |
| chr16:a | 24108 | 25011 | chr9:b | 45773 | 47185 | | | |

* 1-based, inclusive ranges.

#### 5.1 Scheduled scaling of the bead size

Chromosomes expand at the onset of interphase with the repulsive interactions among chromatin chains. Through this expansion process, the factors excluded from the mitotic chromosomes should start to bind to chromatin dependently on the heterogeneous local chromatin features, enhancing the chromatin interactions. We modeled this evolution of interactions by continuously scaling the diameter of 100-kb beads with a predefined schedule throughout the interphase simulation. The scaling factor $A(t)$ was exponentially scheduled, starting from 0.5 and quickly converging to 1:

$$A(t) = 1 - 0.5e^{-t/\tau_A}. \tag{S41}$$

We used the time constant $\tau_A = 0.5\tau_0$, which we estimate to be 0.5 hour. We note that this scaling was not necessary for the phase separation to occur; we obtained the similar results on the genome structure without using this scaling procedure by setting $A(t) = 1$ for all $t$.

#### 5.2 Interactions

The potential energy $U_{\mathrm{inter}}$ of beads in the interphase simulation consists of terms representing repulsions $U_{\mathrm{beads}}$, bonds $U_{\mathrm{chains}}$, nucleolus formation $U_{\mathrm{no}}$, and enclosure by the nuclear envelope $U_{\mathrm{env}}$:

$$U_{\mathrm{inter}} = U_{\mathrm{beads}} + U_{\mathrm{chains}} + U_{\mathrm{no}} + U_{\mathrm{env}}. \tag{S42}$$

##### 5.2.1 Repulsive interactions between chromatin beads

Every overlapping pair of chromatin beads exhibit a repulsive interaction. We write this repulsive interaction between beads $i$ and $j$ as $u_{\mathrm{rep}}(r_{ij}; i, j)$ with $r_{ij} = |\boldsymbol{r}_i - \boldsymbol{r}_j|$. The term $U_{\mathrm{beads}}$ in Eq. S42 is a sum of $u_{\mathrm{rep}}$ as

$$U_{\mathrm{beads}} = \sum_{1 \le i < j \le N_{\mathrm{chr}}} u_{\mathrm{rep}}(r_{ij}; i, j). \tag{S43}$$

Here, $u_{\mathrm{rep}}$ depends on the type of interacting beads $i$ and $j$. When $i$ and $j$ are both type A, we assumed a soft repulsive interaction, $u_{\mathrm{rep}}(r_{ij}; i, j) = u_{\mathrm{AA}}(r_{ij})$, as

$$u_{\mathrm{AA}}(r) = \begin{cases} 0 & r > \sigma_{\mathrm{A}} \\ \varepsilon \left(1 - \left(\frac{r}{\sigma_{\mathrm{A}}}\right)^2\right)^3 & r \le \sigma_{\mathrm{A}} \end{cases}, \tag{S44}$$

which shapes like a Gaussian function, where $\varepsilon$ is a measure of the free-energy cost of overlapping and $\sigma_{\mathrm{A}}$ is a typical size of the chain spreading or the soft diameter. The functional form of Eq. S44 was adopted because this algebraic form approximates a Gaussian with a higher computational efficiency than the Gaussian. When $\varepsilon$ is of the order of the thermal energy $k_{\mathrm{B}}T$, type-A chromatin regions frequently overlap with each other under the thermal fluctuation. On the other hand, the repulsion between two type-B chromatin chains should be harder when two type-B chains begin to overlap because of the condensed nature of the type-B chains. However, once type-B chains manage to overlap, we expect the repulsion to get milder due to the molecular interactions between two type-B chains. We model this situation by writing $u_{\mathrm{rep}}(r_{ij}; i, j) = u_{\mathrm{BB}}(r_{ij})$ as

$$u_{\mathrm{BB}}(r) = \begin{cases} 0 & r > \sigma_{\mathrm{B}} \\ \varepsilon \left(1 - \left(\frac{r}{\sigma_{\mathrm{B}}}\right)^8\right)^3 & r \le \sigma_{\mathrm{B}} \end{cases}, \tag{S45}$$

which shapes rather box-like than the Gaussian-like. For the interaction between two type-u beads, we used the average of the type-A and type-B potentials:

$$u_{\mathrm{uu}}(r) = \frac{1}{2}\left(u_{\mathrm{AA}}(r) + u_{\mathrm{BB}}(r)\right). \tag{S46}$$

Including cases of interactions between different types of beads, the general functional form of $u_{\mathrm{rep}}$ is defined by

$$u_{\mathrm{rep}}(r; i, j) = \alpha_{ij} u_{\mathrm{AA}}(r) + \beta_{ij} u_{\mathrm{BB}}(r), \tag{S47}$$

where the weights $\alpha_{ij}$ and $\beta_{ij}$ are defined by the average of factors in Eq. S40 as

$$\alpha_{ij} = \frac{\alpha_i + \alpha_j}{2}, \qquad \beta_{ij} = \frac{\beta_i + \beta_j}{2}. \tag{S48}$$

The potential strength was set to be $\varepsilon = 2.5k_\mathrm{B}T$. Using the scheduled scaling factor $A(t)$ defined in Eq. S41, the width of the chain spreading was defined as $\sigma_\mathrm{A} = 0.30A(t)\,\mu$m and $\sigma_\mathrm{B} = 0.24A(t)\,\mu$m.

We note that our definition of the repulsive interactions vanish the Flory-Huggins parameter as $\chi = 0$. In many problems of the system with mixed components, the large enough positive value of $\chi$ has been used as a mark of phase separation [32]; when the component X and component Y are mixed, the Flory-Huggins parameter is

$$\chi = u_{XY}(r) - \frac{1}{2}\left(u_{XX}(r) + u_{YY}(r)\right). \tag{S49}$$

With the definition of Eq. S47, we have $\chi = 0$ for every $r$, implying that the phase separation is not induced by the bare effect of the difference in the repulsion strength, but is induced in the present model with the collective motions of different types of beads.

##### 5.2.2 Bond interactions

Bond interactoins are defined for the set $B_\mathrm{chain}$ of every adjacent pair of beads along all chromosome chains:

$$U_\mathrm{inter,chains} = \sum_{(i,j)\in B_\mathrm{chain}} u_\mathrm{bond}(r_{ij}; i, j), \tag{S50}$$

where $u_\mathrm{bond}(r_{ij}; i, j)$ is a harmonic interaction;

$$u_\mathrm{bond}(r; i, j) = \frac{1}{2}K_{ij}r^2. \tag{S51}$$

We assumed that the spring constant $K_{ij}$ depends on types of beads $i$ and $j$. In the similar way to the definition of repulsive interactions in Eq. S47, we used A-A and B-B interactions as bases and defined the average of the two for the mixed-type bonds. In general cases, we have

$$K_{ij} = \alpha_{ij}K_\mathrm{AA} + \beta_{ij}K_\mathrm{BB}, \tag{S52}$$

where $\alpha_{ij}$ and $\beta_{ij}$ are defined in Eq. S48. We defined the spring constants as $K_\mathrm{AA} = 100A^{-2}(t)k_\mathrm{B}T/\mu\mathrm{m}^2$ and $K_\mathrm{BB} = 50A^{-2}(t)k_\mathrm{B}T/\mu\mathrm{m}^2$. Here, we incorporated the scaling factors for consistency with the repulsive potentials. The difference between $K_\mathrm{AA}$ and $K_\mathrm{BB}$ had little effects on the phase separation or A/B compartmentalization, but was crucial to reproducing the experimentally observed decay profile of the intra-chromosomal contact frequency $P(s)$ (Fig. 5D).

##### 5.2.3 Nucleolus formation

We represent the volume of nucleolar mass by the assembly of beads in the model. Let $P_\mathrm{no}$ be the set of nucleolar beads and let $B_\mathrm{no}$ be the set of associated pairs of a nucleolar bead and an rDNA bead in the chromatin chain (Table 5). We defined the potential energy for nucleolus formation as

$$U_\mathrm{no} = \sum_{i,j\in P_\mathrm{no},\, i\neq j} u_\mathrm{no,drop}(r_{ij}) + \sum_{(i,j)\in B_\mathrm{no}} u_\mathrm{no,assoc}(r_{ij}). \tag{S53}$$

The first term is a sum of attractive interactions $u_\mathrm{no,drop}$ between nucleolar beads,

$$u_\mathrm{no,drop}(r) = \begin{cases} 0 & r > 2\sigma_\mathrm{drop} \\ -\frac{\varepsilon_\mathrm{drop}}{1+(r/\sigma_\mathrm{drop})^6} & r \leq 2\sigma_\mathrm{drop} \end{cases}, \tag{S54}$$

where $\varepsilon_\mathrm{drop} = 0.3k_\mathrm{B}T$ and $\sigma_\mathrm{drop} = 0.2\,\mu$m. In the present model, nucleolar beads are associated to the five active nucleolus-organizer regions (NORs) of the genome. The second term in Eq. S53 is a sum of the harmonic bond interactions, representing the tendency of nucloli to reside near the rDNA loci in the NOR;

$$u_{\mathrm{no,assoc}}(r) = \frac{1}{2}K_{\mathrm{no}}r^2. \tag{S55}$$

We used $K_{\mathrm{no}} = 10k_{\mathrm{B}}T/\mu\mathrm{m}^2$, the weak spring interactions allowing the fluctuating motion of nucleolar beads. These nucleolar interactions in the model ensure the formation and occasional merger of nucleoli in the present model. As shown in Fig. S14, when we removed the nucleolar beads from our model, we found no significant change in the results except for the disappearance of the nucleolus-associated domains; therefore, the nucleolar interactions are not necessary for the phase separation or A/B compartmentalization.

When the simulated interphase nucleus reached the stationary state, we counted the number of generated nucleoli and showed its distribution in Fig. 4B of the main text; at the final timestep of each trajectory, positions of nucleolar beads were monitored and clustered using the DBSCAN algorithm provided in the scikit-learn python package. The parameters used were DBSCAN eps = $0.36\,\mu$m, DBSCAN min_samples = 5, and clusters smaller than 100 were ignored. Here, the minimal cluster size of 100 was chosen from the following reasons; two nucleolar beads were associated to each 100-kb bead of the NOR in the present model. The short arm of chr21, which is treated as the shortest NOR in our model, is 10.9 Mb to which 218 nucleolar beads were associated at the starting time step of the interphase simulation. During the interphase simulation, most of these nucleolar beads merged with other beads to form nucleoli (Fig. S2A) but the thermal fluctuation separated a small number of rogue nucleolar beads from the nucleoli (Figs. S2A and S2B). Because these rogue beads would not be detected by the microscope measurment, we did not count these beads as independent nucleoli when their cluster size is smaller than a threshold. The distribution of the calculated cluster size (Fig. S2C) showed that the threshold should be $\approx 100$ beads.

##### 5.2.4 Repulsive nuclear envelope

To enclose the beads inside the nuclear envelope, we defined the repulsive interactions between beads and the nuclear envelope. The potential energy has the interior and exterior parts:

$$U_{\mathrm{env}} = \sum_{i=1}^{N} \left[u_{\mathrm{env,in}}(\boldsymbol{r}_i; i)\theta(\boldsymbol{r}_i) + u_{\mathrm{env,ex}}(\boldsymbol{r}_i)\left(1 - \theta(\boldsymbol{r}_i)\right)\right], \tag{S56}$$

where $\theta(\boldsymbol{r}_i) = 1$ when $\boldsymbol{r}_i$ is inside the nucleus and $\theta(\boldsymbol{r}_i) = 0$ when it is outside. The interior potential $u_{\mathrm{env,in}}$ is the essential part. The exterior potential $u_{\mathrm{env,ex}}$ was added here to prevent beads from jumping out of the nucleus with a large infrequent fluctuation. The interaction between beads and the interior of the nuclear envelope is the average of type-A and type-B repulsive interactions:

$$u_{\mathrm{env,in}}(\boldsymbol{r}; i) = \frac{\alpha_i + \alpha_{\mathrm{env}}}{2} u_{\mathrm{env,A}}(\boldsymbol{r}) + \frac{\beta_i + \beta_{\mathrm{env}}}{2} u_{\mathrm{env,B}}(\boldsymbol{r}) \tag{S57}$$

where $u_{\mathrm{env,A}}$ and $u_{\mathrm{env,B}}$ are similar to the repulsive bead-bead interactions but their width is half the width of the bead-bead interactions because a sphere and the wall start to overlap at the half distance of two spheres start to overlap;

$$u_{\mathrm{env,A}}(\boldsymbol{r}) = \begin{cases} 0 & \delta > \frac{\sigma_{\mathrm{A}}}{2} \\ \varepsilon\left(1 - \left(\frac{\delta}{\sigma_{\mathrm{A}}/2}\right)^2\right)^3 & \delta \leq \frac{\sigma_{\mathrm{A}}}{2}, \end{cases} \tag{S58}$$

$$u_{\mathrm{env,B}}(\boldsymbol{r}) = \begin{cases} 0 & \delta > \frac{\sigma_{\mathrm{B}}}{2} \\ \varepsilon\left(1 - \left(\frac{\delta}{\sigma_{\mathrm{B}}/2}\right)^8\right)^3 & \delta \leq \frac{\sigma_{\mathrm{B}}}{2}. \end{cases} \tag{S59}$$

Here, $\delta$ is the distance between the point $\boldsymbol{r}$ and the nearest point on the nuclear envelope. The nuclear envelope is represented by an ellipsoid with semiaxis lengths $(R_\mathrm{a}, R_\mathrm{b}, R_\mathrm{c})$ in general, while the spherical envelope is a special case with $R_\mathrm{a} = R_\mathrm{b} = R_\mathrm{c}$. As explained later in the Subsection 5.4, $\delta$ is given as the length of the vector

$$\boldsymbol{\delta} = \frac{(\boldsymbol{r} \cdot E\boldsymbol{r} - 1)\, E\boldsymbol{r}}{2\,|E\boldsymbol{r}|^2}, \qquad E = \begin{pmatrix} 1/R_\mathrm{a}^2 & & \\ & 1/R_\mathrm{b}^2 & \\ & & 1/R_\mathrm{c}^2 \end{pmatrix}. \tag{S60}$$

We expect that the nuclear lamina makes the interior of the nuclear envelope rigid. In the present analyses, we consider the case with the rigid interior by adopting $\alpha_\mathrm{env} = 0$ and $\beta_\mathrm{env} = 20$, but the results do not change largely when either $\alpha_\mathrm{env}$ or $\beta_\mathrm{env}$ is large enough. For the exterior part, we used a harmonic constraint

$$u_\mathrm{env,ex}(\boldsymbol{r}) = \frac{1}{2} K_\mathrm{env,ex} \delta^2, \tag{S61}$$

with a sufficiently large spring constant $K_\mathrm{env,ex} = 1000 k_\mathrm{B}T/\mu\mathrm{m}^2$.

##### 5.2.5 Motion of the nuclear envelope

We allowed the nuclear envelope to dynamically expand or shrink under the internal and external pressures. As described in the last subsection, the nuclear envelope pushes the peripheral beads towards inside the nucleus, so the beads exert the opposite reaction force to the nuclear envelope. Therefore, we calculated the force $F_\mathrm{env,in}$ originating from the internal pressure as the sum of repulsive forces between beads and the nuclear envelope projected onto the surface normal:

$$F_\mathrm{env,in} = \sum_{i=1}^{N} \left| \frac{\partial U_\mathrm{env}}{\partial \boldsymbol{r}_i} \right|. \tag{S62}$$

We also assumed the cytoskeletal force acting on the nuclear membrane as an external pressure and thus applied an anisotropic harmonic potential to the semiaxes:

$$U_\mathrm{env,ex} = \frac{1}{2} \left( K_\mathrm{env,a} R_\mathrm{a}^2 + K_\mathrm{env,b} R_\mathrm{b}^2 + K_\mathrm{env,c} R_\mathrm{c}^2 \right). \tag{S63}$$

The parameters used are $K_\mathrm{env,a} = 0.75 \times 10^4 k_\mathrm{B}T/\mu\mathrm{m}^2$, $K_\mathrm{env,b} = 3.0 \times 10^4 k_\mathrm{B}T/\mu\mathrm{m}^2$ and $K_\mathrm{env,c} = 12.0 \times 10^4 k_\mathrm{B}T/\mu\mathrm{m}^2$ for the IMR90 nucleus and $K_\mathrm{env,a} = K_\mathrm{env,b} = K_\mathrm{env,c} = 3.0 \times 10^4 k_\mathrm{B}T/\mu\mathrm{m}^2$ for the GM12878 nucleus. Under these forces, the semiaxis length $R_X$ of each axis $X = \mathrm{a}, \mathrm{b}$ and c was assumed to obey the overdamped dynamics:

$$\frac{dR_X}{dt} = \mu_\mathrm{env} \left( F_\mathrm{env,in} - \frac{\partial U_\mathrm{env,ex}}{\partial R_X} \right). \tag{S64}$$

We used $\mu_\mathrm{env} = 2 \times 10^{-4}\, \mu\mathrm{m}^2/\tau_0 k_\mathrm{B}T$ as the mobility in Eq. S64, which is much smaller than the mobility of beads in the nucleus. With such an extremely small mobility, the nucleus expands much slower than the chromatin movement. For all the cell types, we used $R_\mathrm{a} = R_\mathrm{b} = R_\mathrm{c} = 2\,\mu\mathrm{m}$ as the initial condition at the beginning of interphase.

#### 5.3 The Langevin dynamics simulation

Along with the motion of the nuclear envelope, we simulated the overdamped Langevin dynamics of chromatin and the nucleolar beads:

$$\frac{d\boldsymbol{r}_i}{dt} = -\mu \frac{\partial U_\mathrm{inter}}{\partial \boldsymbol{r}_i} + \sqrt{2\mu k_\mathrm{B}T}\, \boldsymbol{\xi}_i(t). \tag{S65}$$

Here, $\mu$ is the mobility of a bead and $\boldsymbol{\xi}_i(t)$ is a Gaussian white noise vector satisfying $\langle \xi_{i\nu}(t_1)\xi_{j\nu'}(t_2)\rangle = \delta_{ij}\delta_{\nu\nu'}\delta(t_1 - t_2)$, where $\nu$ and $\nu'$ label the $xyz$ component of the vector. Chemical reactions in the nucleus may generate excess heat that dissipates into the nucleoplasm around the reaction sites. This dissipation should be much faster than the chromatin motion with the time scale of $\tau_0$; therefore, we approximately neglect the detailed-balance breaking fluctuations arising from the excess heat assuming that the nonequilibrium activating effects are implicitly considered in the renormalized homogeneous $T$, which is independent of the chromatin locus $i$ in Eq. S65.

Starting from the initial configuration generated by fine-graining the anatelophase structure, we first simulated a short relaxation for 10,000 steps and then simulated for 700,000 steps for the production sampling. The discretization step was $\delta t = 10^{-5}\tau_0$. We used the same mobility for all beads, $\mu = 1\,\mu\mathrm{m}^2/\tau_0 k_\mathrm{B}T$. If we consider that the production sampling time $700000 \times \delta t = 7\tau_0$ roughly corresponds to the duration of G1 phase $\sim 7\,\mathrm{h}$, the unit simulation time should be $\tau_0 = 3.6 \times 10^3\,\mathrm{s}$. With these units, the single-particle diffusion coefficient should be $D = \mu k_\mathrm{B}T = 2\times10^{-4}\,\mu\mathrm{m}^2/\mathrm{s}$, giving a reasonable value for the motion of a 100-kb chromatin region in interphase nucleus. 200 independent simulations were run for each type of cell (IMR90 and GM12878) using different random seeds (thus with different initial conditions).

#### 5.4 Interactions with the surface

In this subsection, we describe a general mathematical expression for the short-range interaction between a point mass and a smooth surface.

##### 5.4.1 Surface translation vector for simulations

Let $\phi(\boldsymbol{r})$ be a smooth function that implicitly defines a surface $S = \{\boldsymbol{r} : \phi(\boldsymbol{r}) = 0\}$. Now, let $\boldsymbol{r}$ be a point near the surface $S$. Let us find the shortest vector $\boldsymbol{\delta}$ from the surface $S$ to the point $\boldsymbol{r}$. Because the point $\boldsymbol{r} - \boldsymbol{\delta}$ is on the surface, we have

$$\phi(\boldsymbol{r} - \boldsymbol{\delta}) = 0. \tag{S66}$$

Solving Eq. S66, the vector $\boldsymbol{\delta}$ is expressed as a function of $\boldsymbol{r}$. We call this vector field $\boldsymbol{\delta}(\boldsymbol{r})$ the surface translation vector.

Linear approximation gives

$$0 = \phi(\boldsymbol{r} - \boldsymbol{\delta}) \simeq \phi(\boldsymbol{r}) - \nabla\phi(\boldsymbol{r}) \cdot \boldsymbol{\delta}, \tag{S67}$$

which represents a plane defined by the vector $\boldsymbol{\delta}$. When $\delta = |\boldsymbol{\delta}|$ is small enough, we can simply choose a solution vector $\boldsymbol{\delta}$ that is directed along the gradient of the function $\phi$:

$$\boldsymbol{\delta}(\boldsymbol{r}) = \delta(\boldsymbol{r})\boldsymbol{n}(\boldsymbol{r}), \qquad \boldsymbol{n}(\boldsymbol{r}) = \frac{\nabla\phi(\boldsymbol{r})}{|\nabla\phi(\boldsymbol{r})|}. \tag{S68}$$

This expression asymptotically gives the solution of the shortest vector $\boldsymbol{\delta}$ when $\phi(\boldsymbol{r}) \to 0$. Then, we can solve the plane equation for the scalar factor $\delta(\boldsymbol{r})$, and we have

$$\delta(\boldsymbol{r}) = \frac{\phi(\boldsymbol{r})}{|\nabla\phi(\boldsymbol{r})|} \quad \text{and} \quad \boldsymbol{\delta}(\boldsymbol{r}) = \frac{\phi(\boldsymbol{r})\nabla\phi(\boldsymbol{r})}{|\nabla\phi(\boldsymbol{r})|^2}. \tag{S69}$$

##### 5.4.2 Near-surface force

Now, suppose that there is a short-range interaction between the surface and a point near the surface. The potential energy $U(\boldsymbol{r})$ is defined using the surface translation vector as

$$U(\boldsymbol{r}) = u(\boldsymbol{\delta}(\boldsymbol{r})) \tag{S70}$$

where $u$ is a base potential function. Let $\boldsymbol{f}(\boldsymbol{\delta}) = -\frac{\partial}{\partial \boldsymbol{\delta}} u(\boldsymbol{\delta})$ be the base force function, and let us calculate the force $\boldsymbol{F}(\boldsymbol{r}) = -\nabla U(\boldsymbol{r})$ with $\nabla = \frac{\partial}{\partial \boldsymbol{r}}$. The chain rule dictates that

$$\boldsymbol{F}(\boldsymbol{r}) = -\nabla u(\boldsymbol{\delta}(\boldsymbol{r})) = \nabla \boldsymbol{\delta}(\boldsymbol{r}) \cdot \boldsymbol{f}(\boldsymbol{\delta}(\boldsymbol{r})). \tag{S71}$$

Here, $\nabla \boldsymbol{\delta} = \sum_i \partial \boldsymbol{\delta} / \partial x_i \otimes \hat{\boldsymbol{x}}_i$ (where $\hat{\boldsymbol{x}}_i$ is the basis vector with respect to the Cartesian coordinate $x_i$) is the derivative of the vector field $\boldsymbol{\delta}$, which results in a rank-2 tensor field. Manipulation gives:

$$\begin{aligned}
\nabla \boldsymbol{\delta}(\boldsymbol{r}) &= \nabla \left( \frac{\phi(\boldsymbol{r}) \nabla \phi(\boldsymbol{r})}{|\nabla \phi(\boldsymbol{r})|^2} \right) && \text{(S72)} \\
&= \nabla \phi(\boldsymbol{r}) \otimes \frac{\nabla \phi(\boldsymbol{r})}{|\nabla \phi(\boldsymbol{r})|^2} + \nabla\nabla \phi(\boldsymbol{r}) \frac{\phi(\boldsymbol{r})}{|\nabla \phi(\boldsymbol{r})|^2} + \nabla \left( \frac{1}{|\nabla \phi(\boldsymbol{r})|^2} \right) \otimes (\phi(\boldsymbol{r}) \nabla \phi(\boldsymbol{r})) && \text{(S73)} \\
&= \frac{\nabla \phi(\boldsymbol{r}) \otimes \nabla \phi(\boldsymbol{r})}{|\nabla \phi(\boldsymbol{r})|^2} + \frac{\phi(\boldsymbol{r}) \nabla\nabla \phi(\boldsymbol{r})}{|\nabla \phi(\boldsymbol{r})|^2} - \frac{2\phi(\boldsymbol{r}) \left( \nabla\nabla \phi(\boldsymbol{r}) \cdot \nabla \phi(\boldsymbol{r}) \right) \otimes \nabla \phi(\boldsymbol{r})}{|\nabla \phi(\boldsymbol{r})|^4} && \text{(S74)} \\
&= K(\boldsymbol{r}) + (I - 2K(\boldsymbol{r})) \frac{\nabla \phi(\boldsymbol{r}) \otimes \nabla \phi(\boldsymbol{r})}{|\nabla \phi(\boldsymbol{r})|^2} && \text{(S75)}
\end{aligned}$$

where $I$ is the identity matrix and $K$ is a rank-2 tensor field given by

$$K(\boldsymbol{r}) = \frac{\phi(\boldsymbol{r}) \nabla\nabla \phi(\boldsymbol{r})}{|\nabla \phi(\boldsymbol{r})|^2}. \tag{S76}$$

Here, $\nabla\nabla \phi(\boldsymbol{r}) = \sum_i \sum_j \partial^2 \phi / \partial x_i \partial x_j \, \hat{\boldsymbol{x}}_i \otimes \hat{\boldsymbol{x}}_j$.

##### 5.4.3 Force calculation for an ellipsoidal surface

Let the surface be an ellipsoid given by a symmetric positive definite matrix $E$:

$$\phi(\boldsymbol{r}) = \boldsymbol{r} \cdot E\boldsymbol{r} - 1. \tag{S77}$$

Then, we have

$$\nabla \phi(\boldsymbol{r}) = 2E\boldsymbol{r} \quad \text{and} \quad \nabla\nabla \phi(\boldsymbol{r}) = 2E. \tag{S78}$$

So, the surface displacement vector is given by

$$\boldsymbol{\delta}(\boldsymbol{r}) = \frac{(\boldsymbol{r} \cdot E\boldsymbol{r} - 1) E\boldsymbol{r}}{2 |E\boldsymbol{r}|^2}, \tag{S79}$$

which gives Eq. S60, and its derivative is

$$\nabla \boldsymbol{\delta}(\boldsymbol{r}) = K(\boldsymbol{r}) + (I - 2K(\boldsymbol{r})) \frac{E\boldsymbol{r} \otimes E\boldsymbol{r}}{|E\boldsymbol{r}|^2}, \qquad K(\boldsymbol{r}) = \frac{(\boldsymbol{r} \cdot E\boldsymbol{r} - 1) E}{2 |E\boldsymbol{r}|^2}. \tag{S80}$$

With these expressions in hand, we can calculate the near-surface force $\boldsymbol{F}$ by passing $\boldsymbol{\delta}(\boldsymbol{r})$ to the base force function $\boldsymbol{f}$ and transforming the result with $\nabla \boldsymbol{\delta}(\boldsymbol{r})$ as

$$\boldsymbol{F}(\boldsymbol{r}) = \nabla \boldsymbol{\delta}(\boldsymbol{r}) \cdot \boldsymbol{f}(\boldsymbol{\delta}(\boldsymbol{r})). \tag{S81}$$

##### 5.4.4 Surface translation vector for analyses

The linear approximation in Eq. S67 is accurate enough for calculating the short-range interactions with $\delta \lesssim 0.15\,\mu$m in the simulation. However, it is not suitable for analyzing the simulated results on the long-range distribution of beads spanning up to $\delta \sim 1.2\,\mu$m. Hence, we used a computationally expensive but more accurate estimation of the surface translation vector in our analyses. Let us calculate the surface translation vector $\boldsymbol{\delta}$ for an ellipsoid:

$$\begin{aligned} 0 &= \phi(\boldsymbol{r} - \boldsymbol{\delta}) \\ &= (\boldsymbol{r} - \boldsymbol{\delta}) \cdot E\,(\boldsymbol{r} - \boldsymbol{\delta}) - 1 \\ &= \boldsymbol{\delta} \cdot E\boldsymbol{\delta} - \boldsymbol{\delta} \cdot E\boldsymbol{r} - \boldsymbol{r} \cdot E\boldsymbol{\delta} + \boldsymbol{r} \cdot E\boldsymbol{r} - 1. \end{aligned} \tag{S82}$$

Now, assuming that the surface translation vector is directed along the gradient $\nabla\phi(\boldsymbol{r}) = 2E\boldsymbol{r}$, we have $\boldsymbol{\delta} = sE\boldsymbol{r}$ with a scalar factor $s$. Then, Eq. S82 becomes

$$0 = \boldsymbol{r} \cdot E\boldsymbol{r} - 2\,(\boldsymbol{r} \cdot EE\boldsymbol{r})\,s + (E\boldsymbol{r} \cdot EE\boldsymbol{r})\,s^2 - 1. \tag{S83}$$

The smaller solution for this quadratic equation gives the surface translation vector:

$$\boldsymbol{\delta}(\boldsymbol{r}) = sE\boldsymbol{r}, \quad s = \frac{\boldsymbol{r} \cdot EE\boldsymbol{r} - \sqrt{(\boldsymbol{r} \cdot EE\boldsymbol{r})^2 - (E\boldsymbol{r} \cdot EE\boldsymbol{r})\,(\boldsymbol{r} \cdot E\boldsymbol{r} - 1)}}{E\boldsymbol{r} \cdot EE\boldsymbol{r}}. \tag{S84}$$

### 6 Model of a polymer-blend system

As an example system having heterogeneous repulsive interactions, we simulated a simplified polymer-blend system as shown in Fig. 3. The system is a mixture of 100 homopolymers, each of which is a beads-on-a-string chain consisting of 20 type-A beads, and 100 homopolymers, each of which is a beads-on-a-string chain consisting of 20 type-B beads. Therefore, the system dynamics are described by the movement of $N_{\text{blend}} = 100 \times 20 + 100 \times 20 = 4000$ beads. We denote by $\boldsymbol{r}_i$ the three-dimensional coordinate vector of the $i$th bead with $1 \leq i \leq N_{\text{blend}}$. Similarly to Eq. S40, we define factors $\alpha_i$ and $\beta_i$ to distinguish the type of beads as

$$\alpha_i = \begin{cases} 1 & \text{bead } i \text{ is type-A} \\ 0 & \text{bead } i \text{ is type-B} \end{cases}, \qquad \beta_i = \begin{cases} 0 & \text{bead } i \text{ is type-A} \\ 1 & \text{bead } i \text{ is type-B} \end{cases}. \tag{S85}$$

#### 6.1 Simulation in a periodic box

We packed 200 type-A and type-B chains in a cubic box with the periodic boundary condition. We set the side length of the box to be $L_{\text{blend}} = 2.5\,\mu$m so that the density approximately matches that of chromatin beads in the interphase genome simulation.

##### 6.1.1 Force field

We assumed the repulsive interactions between overlapping beads and the harmonic bonds along each chain. The potential energy of the chains was given by

$$U_{\text{blend}} = \sum_{1 \leq i < j \leq N_{\text{blend}}} u_{\text{rep}}(r_{ij}; i, j) + \sum_{(i,j) \in B_{\text{blend}}} u_{\text{blend,bond}}(r_{ij}) \tag{S86}$$

where $r_{ij} = |\boldsymbol{r}_i - \boldsymbol{r}_j|$ and $B_{\text{blend}}$ is a set of all the pairs of bonded beads. The repulsive potential $u_{\text{rep}}$ is the same as that used in the interphase simulations;

$$u_{\text{rep}}(r; i, j) = \frac{\alpha_i + \alpha_j}{2} u_{\text{AA}}(r) + \frac{\beta_i + \beta_j}{2} u_{\text{BB}}(r) \tag{S87}$$

with the same definitions of $u_{\text{AA}}(r)$ and $u_{\text{BB}}(r)$ and the same parameters. The bond potential is a harmonic interaction

$$u_{\text{blend,bond}}(r) = \frac{1}{2} K_{\text{blend}} r^2 \tag{S88}$$

with the uniform spring constant $K_{\text{blend}} = 70\,k_{\text{B}}T/\mu\text{m}^2$.

##### 6.1.2 Initial configuration

The initial configuration of each chain was chosen to be a randomly directed straight rod. Denoting the bead number at start and end of $k$th chain with $s_k$ and $e_k$, respectively, the initial coordinate of the $i$th bead on the $k$th chain with $k = 1, \ldots, 200$ was given by

$$\boldsymbol{r}_i = \boldsymbol{c}_k + \left( i - \frac{s_k + e_k}{2} \right) \sigma_{\mathrm{blend,rod}} \boldsymbol{w}_k \qquad (s_k \leq i \leq e_k) \tag{S89}$$

where $\boldsymbol{c}_k$ is a uniformly distributed random point in the box, $\boldsymbol{w}_k$ is a randomly directed unit vector and $\sigma_{\mathrm{blend,rod}}$ is the initial bond length. We set $\sigma_{\mathrm{blend,rod}} = 0.1\,\mu\mathrm{m}$ in our simulations.

##### 6.1.3 The Langevin dynamics simulation

We simulated the overdamped Langevin dynamics of the beads:

$$\frac{d\boldsymbol{r}_i}{dt} = -\mu \frac{\partial U_{\mathrm{blend}}}{\partial \boldsymbol{r}_i} + \sqrt{2\mu k_{\mathrm{B}} T} \boldsymbol{\xi}_i(t). \tag{S90}$$

Here, $\mu$ is the mobility of a bead and $\boldsymbol{\xi}_i(t)$ is a Gaussian white noise vector satisfying $\langle \xi_{i\nu}(t_1) \xi_{j\nu'}(t_2) \rangle = \delta_{ij} \delta_{\nu\nu'} \delta(t_1 - t_2)$, where $\nu$ and $\nu'$ label the $xyz$ component of the vector. Starting from the initial configuration described above, we simulated the dynamics for 1,000,000 steps with the discretization step $\delta t = 10^{-5} \tau_0$. We used the same mobility for all beads $\mu = 1\,\mu\mathrm{m}^2/\tau_0 k_{\mathrm{B}} T$.

#### 6.2 Simulation in a spherical container

We packed 200 chains in a sphere with the wall potentials. We set the radius of the sphere to be $R_{\mathrm{blend}} = 1.6\,\mu\mathrm{m}$, so that the density approximately matches that of chromatin beads in the interphase genome simulation.

##### 6.2.1 Force field

For simulations in a spherical container, we imposed additional potentials to constrain the beads into the sphere:

$$U_{\mathrm{blend,sphere}} = U_{\mathrm{blend}} + \sum_{i=1}^{N_{\mathrm{blend}}} \left[ u_{\mathrm{env,in}}(\boldsymbol{r}_i) \theta(\boldsymbol{r}_i) + u_{\mathrm{env,ex}}(\boldsymbol{r}_i) \left(1 - \theta(\boldsymbol{r}_i)\right) \right], \tag{S91}$$

where $\theta(\boldsymbol{r}_i) = 1$ when $\boldsymbol{r}_i$ is inside the sphere and $\theta(\boldsymbol{r}_i) = 0$ when it is outside. The potentials $u_{\mathrm{env,in}}(\boldsymbol{r})$ and $u_{\mathrm{env,ex}}(\boldsymbol{r})$ are the same as the ones used in the interphase simulations, except that we defined the distance from the wall as $\delta(r) = |r - R_{\mathrm{blend}}|$ for this simulation.

##### 6.2.2 Initial configuration

The chains were initialized as randomly directed straight rods in the sphere in the similar way as in the periodic box by placing the cetroid of each rod $\boldsymbol{c}_k$ at the uniformly distributed random position in the sphere. We allowed the rods to extrude out of the sphere in the initial configuration; the extruding parts were quickly pulled back into the sphere by the action of $U_{\mathrm{blend,sphere}}$.

##### 6.2.3 The Langevin dynamics simulation

The Langevin dynamics simulations were carried out in the same protocol as in the periodic box.

##### 6.2.4 Analysis of phase-separation dynamics

We denote by $\boldsymbol{r}_i(t)$ the trajectory of the $i$th bead, and we sampled the values of $\boldsymbol{r}_i(t)$ from $t = \tau_0$ to $2\tau_0$ at the interval of $10^{-2}\tau_0$ (1000 steps). We write this set of time instances as $T_{\text{blend,MSD}}$. We calculated the short-time mean-square distance (MSD) of the $i$th bead,

$$m_i = \left\langle |\boldsymbol{r}_i(t+\tau) - \boldsymbol{r}_i(t)|^2 \right\rangle_{t \in T_{\text{blend,MSD}}}, \quad \tau = 10^{-2}\tau_0, \tag{S92}$$

as a measure of the fluctuation of the $i$th bead. For visualization, we randomly generated 10000 mesh points, $\boldsymbol{s}_1, \ldots, \boldsymbol{s}_{10000}$ with $\boldsymbol{s}_k = (x_k, y_k)$, on a square region $-1.8\,\mu\text{m} \leq x_k, y_k \leq 1.8\,\mu\text{m}$ covering the cross section of the spherical container in the simulation. Then, the MSD values $m_i$ of beads were averaged in a region around each mesh point $\boldsymbol{s}_k$ as

$$m(\boldsymbol{s}_k) = \langle m_i \rangle_{|\boldsymbol{s}_k - \boldsymbol{r}_i(\tau_0)| < \Delta}, \quad \Delta = 0.12\,\mu\text{m}, \tag{S93}$$

which defines the average MSD at $\boldsymbol{s}_k$. In Fig. 3C, this scalar field $m(\boldsymbol{s}_k)$ is displayed as a gray-scale contour map.

In Fig. 3C, we also show the flow of bead movement, which leads to phase separation, by calculating the displacement $\boldsymbol{u}_i$ of beads during the time interval $\tau_0$ as

$$\boldsymbol{u}_i = \boldsymbol{r}_i(2\tau_0) - \boldsymbol{r}_i(\tau_0). \tag{S94}$$

For visualization, we defined an evenly spaced mesh $\boldsymbol{q}_k = (x_k, y_k)$ on the square region $-1.8\,\mu\text{m} \leq x_k, y_k \leq 1.8\,\mu\text{m}$ with the space interval $0.05\,\mu\text{m}$ along each coordinate axis. Here, the even spacing was adopted for legibility of the results. The average displacement $\boldsymbol{u}(\boldsymbol{q}_k)$ around each mesh point $\boldsymbol{q}_k$ was calculated as

$$\boldsymbol{u}(\boldsymbol{q}_k) = \langle \boldsymbol{u}_i \rangle_{|\boldsymbol{q}_k - \boldsymbol{r}_i(\tau_0)| < \Delta}, \quad \Delta = 0.12\,\mu\text{m}. \tag{S95}$$

Thus obtained two fields, $m(\boldsymbol{s})$ in gray scale and $\boldsymbol{u}(\boldsymbol{q})$ with arrows, are superposed in Fig. 3C. The figure shows that adjacent type-A and type-B regions move in different directions, and the type-A regions fluctuate more largely than the type-B regions.

### 7 Fluctuating dynamics

The present model shows a clear correlation between compartmentalization and the fluctuating dynamics: The type-A chromatin regions fluctuate larger than the type-B regions, thus allowing type-A chromatin domains to proactively merge into the larger A compartment. We describe the analysis that supports this view in this section and Figs. S15 and S16.

#### 7.1 Slow and fast chromatin regions

We arbitrarily chose an example simulation run for the IMR90 cell and sampled the dynamics from $t = 6\tau_0$ to $7\tau_0$ at the interval of $10^{-3}\tau_0$ (100 steps). We denote by $\boldsymbol{r}_i(t)$ the trajectory of $i$th bead and by $T_{\text{MSD}}$ the set of analyzed time points. Then, the mean-squared displacement (MSD) with a lag time $\tau$ was computed:

$$\text{MSD}_i(\tau) = \left\langle |\boldsymbol{r}_i(t+\tau) - \boldsymbol{r}_i(t)|^2 \right\rangle_{t \in T_{\text{MSD}}}. \tag{S96}$$

We calculated the average MSD at the shortest non-zero lag time over the chromatin regions

$$\mathit{\Delta}^2 = \frac{1}{N_{\text{chr}}} \sum_{i=1}^{N_{\text{chr}}} \text{MSD}_i(10^{-3}\tau_0) \tag{S97}$$

$$= 4.3 \times 10^{-3}\,\mu\text{m}^2 \tag{S98}$$

and labeled those chromatin regions with $\text{MSD}_i(10^{-3}\tau_0) < \mathit{\Delta}^2$ as *slow* and the other chromatin regions as *fast*. These labels are used in the pair-correlation function analysis described below.

##### 7.2 Time-dependent diffusion coefficients

We calculated the average MSDs of type-A beads and type-B beads:

$$\mathrm{MSD}_X(\tau) = \langle \mathrm{MSD}_i(\tau) \rangle_{i \text{ is type-}X}\,, \quad X \in \{\mathrm{A}, \mathrm{B}\}\,. \tag{S99}$$

$\mathrm{MSD}_\mathrm{A}(\tau)$ and $\mathrm{MSD}_\mathrm{B}(\tau)$ behave differently in the short lag time $\tau < 1\,\mathrm{min}$ and similarly in the longer lag time (Fig. S15A) from/to each other. To characterize the diffusion characteristics in different timescales, we calculated the time-dependent diffusion coefficient for each bead type:

$$D_X(\tau) = \frac{1}{6}\frac{d\mathrm{MSD}_X(\tau)}{d\tau}, \quad X \in \{\mathrm{A}, \mathrm{B}\}\,. \tag{S100}$$

The time-dependent diffusion coefficient $D(\tau)$ in general describes the diffusive motion of a particle at different timescales [33, 34], and by definition, MSD with the lag time $\tau$ is given by collecting infinitesimal MSD, $6D(\tau')\,d\tau'$, over the lag time: $\mathrm{MSD}(\tau) = \int_0^\tau 6D(\tau')\,d\tau'$.

Fig. S15B shows $D_\mathrm{A}(\tau)$ and $D_\mathrm{B}(\tau)$ differing only in the short timescale of $\tau < 1\,\mathrm{min}$ and converging to the same function as $\tau \to \infty$. The power-law scaling of the convergent region was $D_X(\tau) \propto \tau^{-0.55}$, meaning that the MSDs approach to the same subdiffusive behavior $\mathrm{MSD}_X(\tau) \propto \tau^\alpha$ with the anomalous exponent $\alpha = -0.55 + 1 = 0.45$ in the long timescale. In the shorter timescale, the type-A chromatin regions diffuse more freely than the type-B chromatin regions. This short-time difference is the hallmark of A/B compartmentalization.

##### 7.3 Normalized pair-correlation functions

We analyzed the spatial distribution of each type of chromatin region by calculating the normalized functions of pair-correlation in distance. Let $X$ and $Y$ be two types of chromatin regions; $X$ and $Y$ can be either A, B, or u as explained above, or fast ($\mathcal{F}$) or slow ($\mathcal{S}$) as discussed at the end of this subsection. Let $\boldsymbol{r}_i$ be the coordinate vector of the $i$th chromatin region at time $t = 6\tau_0$ in a simulation run. We count the number of inter-chromosomal pairs of type-$X$ regions and type-$Y$ regions separated by distance $d$:

$$H_{XY}^{\mathrm{pair}}(d) = \#\left\{(i,j) : d \le r_{ij} < d + \Delta d^{\mathrm{pair}},\ i \in X,\ j \in Y,\ c_i \ne c_j\right\} \tag{S101}$$

where $r_{ij} = |\boldsymbol{r}_i - \boldsymbol{r}_j|$ is the distance between the $i$th and the $j$th chromatin regions, $\Delta d^{\mathrm{pair}}$ is a bin size, and $c_i$ designates the chromosome that contains the $i$th chromatin region. We used $\Delta d^{\mathrm{pair}} = 0.04\,\mu\mathrm{m}$ as the bin size. Intra-chromosomal pairs are excluded from the analysis because a trivial cluster of the similarly-labeled chromatin regions are included in each TAD in the same chromosome, which obscures the spatial correlation of chromatin regions induced in the phase separation dynamics..

$H_{XY}^{\mathrm{pair}}(d)$ can be interpreted as the sum of the number of type-$Y$ regions around each type-$X$ region. Therefore, dividing $H_{XY}^{\mathrm{pair}}(d)$ by the number of type-$X$ regions and by the volume $V^{\mathrm{pair}}(d) = \frac{4\pi}{3}\left(\left(d + \Delta d^{\mathrm{pair}}\right)^3 - d^3\right)$ of the bin at distance $d$, we get the average density of type-$Y$ regions at distance $d$ from a type-$X$ region;

$$\rho_{XY}^{\mathrm{pair}}(d) = \frac{1}{\#X}\frac{H_{XY}^{\mathrm{pair}}(d)}{V^{\mathrm{pair}}(d)}. \tag{S102}$$

The semiaxes lengths of the nuclear envelope in the simulation were $R_\mathrm{a} = 8.28\,\mu\mathrm{m}$, $R_\mathrm{b} = 4.18\,\mu\mathrm{m}$ and $R_\mathrm{c} = 2.11\,\mu\mathrm{m}$. Therefore, the volume $V$ of the nucleus was

$$V = \frac{4\pi}{3}R_\mathrm{a}R_\mathrm{b}R_\mathrm{c} = 306\,\mu\mathrm{m}^3, \tag{S103}$$

and the expected density of type-$Y$ regions in the entire nucleus is

$$\rho_Y = \frac{\#Y}{V}. \tag{S104}$$

Then, the pair-correlation function $g_{XY}(d)$ between type-$X$ and $Y$ regions is defined as

$$g_{XY}(d) = \frac{\rho^{\mathrm{pair}}_{XY}(d)}{\rho_Y}. \tag{S105}$$

The pair-correlation function $g_{XY}(d)$ decays to zero as the $d$ grows beyond $\sim 1\,\mu$m because the void space outside the nucleus in the simulation is sampled with the large $d$. This makes it hard to identify a decay of the pair correlation arising from the clustering of chromatin regions. To cancel out this effect, we normalize $g_{XY}(d)$ by the reference pair-correlation function among all inter-chromosome chromatin regions, $g(d)$, as

$$\tilde{g}_{XY}(d) = \frac{g_{XY}(d)}{g(d)}, \tag{S106}$$

where $g(d) = g_{\mathcal{C}\mathcal{C}}(d)$ with $\mathcal{C} = \{i : 1 \le i \le N_{\mathrm{chr}}\}$. Here, $\tilde{g}_{XY}(d)$ is the *normalized pair-correlation function* between type-$X$ and type-$Y$ regions in the nucleus; the normalized pair-correlation functions converge to $\tilde{g}_{XY}(d) \to 1$ as $d \to \infty$, showing that the decaying profile of $\tilde{g}_{XY}(d)$ captures the clustering of type-$Y$ regions around the type-$X$ regions.

The slow and fast chromatin regions described in the subsection 7.1 are defined as the sets

$$\mathcal{S} = \left\{ i \in \mathcal{C} : \mathrm{MSD}_i(10^{-3}\tau_0) < \mathit{\Delta}^2 \right\}, \tag{S107}$$

$$\mathcal{F} = \left\{ i \in \mathcal{C} : \mathrm{MSD}_i(10^{-3}\tau_0) \ge \mathit{\Delta}^2 \right\}. \tag{S108}$$

We calculated the normalized pair-correlation functions among slow and fast chromatin regions, $\tilde{g}_{\mathcal{S}\mathcal{S}}(d)$, $\tilde{g}_{\mathcal{S}\mathcal{F}}(d)$ and $\tilde{g}_{\mathcal{F}\mathcal{F}}(d)$, and plotted them in Fig. S16.

### 8 Captions for Supplementary Figures and Movies

**Fig. S1** : **Characterization of type-A and type-B chromatin in FG simulations.** (**A**) Coarse-grained interaction potentials $U_{\mathrm{CG}}(r)$ before applying smoothing. The coarse-grained potential energy between 100-kb type-A chromatin regions (left), type-B chromatin regions without cohesin (middle) and type-B chromatin regions with cohesin (right) are shown for various chromatin density $\rho$. Solid lines are the potentials derived with the PRISM theory and dashed lines are the potentials of mean force. The plots confirm that the smoothing applied in Fig. 2D-2F in the main text does not alter the essential shape of the potentials. (**B**) Type-B coarse-grained potentials with different cohesin loading constant $k_{\mathrm{on}}/k_{\mathrm{off}} = 0, 0.05, 0.1\,\mathrm{kb}^{-1}$. The potential curve becomes steeper as more cohesin is loaded. (**C**) The simulated radius of gyration of type-A (red), cohesin-loaded type-B (blue) and cohesin-free type-B (pale blue) chromatin as a function of the length of a subdomain. The curve for cohesin-free type-B chromatin does not show a saturating behavior.

**Fig. S2** : **Simulated nucleoli defined as clusters of nucleolar beads.** (**A**) Clustering of the distributied nucleolar beads in an example IMR90 cell. Simulated nucleolar beads were clustered according to their three-dimensional positions and projected onto the two-dimensional xy plane for visualization. Five clusters (blue, cyan, green, yellow and red) and the DBSCAN outliars (purple) were detected. (**B**) Cluster size of the example shown in **A**. The first and fourth clusters are small, representing the rogue beads separated by thermal fluctuations from nucleoli; these rogue beads were not counted as nucleoli in Fig. 4B of the main text. (**C**) Distribution of the cluster size in 200 simulated IMR90 cells. The dotted line at the cluster size 100 separates the small-cluster outliars.

**Fig. S3** : **Comparison of the experimentally observed (Rao et al 2014 *Cell*, lower left in each panel) and simulated (upper right in each panel) genome wide contact frequencies.** Top: GM12878 and Bottom: IMR90. Shown with a 1 Mb resolution. The observed and the simulated contact frequencies are normalized to the 93-percentile of the respective data.

**Fig. S4** :**Comparison of the experimentally observed (Rao et al 2014 *Cell*, lower left in each panel) and simulated (upper right in each panel) contact frequencies of chromosomes 1 to 12 of GM12878.** Shown with a 100 kb resolution. The observed and the simulated contact frequencies are normalized to the 96-percentile of the respective data. The experimental data are lacking in the gray shaded areas.

**Fig. S5** : **Comparison of the experimentally observed (Rao et al 2014 *Cell*, lower left in each panel) and simulated (upper right in each panel) contact frequencies of chromosomes 13 to 22 and chromosome X of GM12878.** Data of active and inactive X chromosomes were averaged. Shown with a 100 kb resolution. The observed and the simulated contact frequencies are normalized to the 96-percentile of the respective data. The experimental data are lacking in the gray shaded areas.

**Fig. S6** : **Comparison of the experimentally observed (Rao et al 2014 *Cell*, lower left in each panel) and simulated (upper right in each panel) contact frequencies of chromosomes 1 to 12 of IMR90.** Shown with a 100 kb resolution. The observed and the simulated contact frequencies are normalized to the 96-percentile of the respective data. The experimental data are lacking in the gray shaded areas.

**Fig. S7** : **Comparison of the experimentally observed (Rao et al 2014 *Cell*, lower left in each panel) and simulated (upper right in each panel) contact frequencies of chromosomes 13 to 22 and chromosome X of IMR90.** Data of active and inactive X chromosomes were averaged. Shown with a 100 kb resolution. The observed and the simulated contact frequencies are normalized to the 96-percentile of the respective data. The experimental data are lacking in the gray shaded areas.

**Fig. S8** : **Comparison of the simulated chromosomes having randomly annotated chromatin types with the experimental data.** (**A**) Comparison of the experimentally observed (Rao et al 2014 *Cell*, lower left in each panel) and simulated (upper right in each panel) contact frequencies of chromosomes 1, 10, and 19 of IMR90, shown with a 100 kb resolution. The observed and the simulated contact frequencies are normalized to the 95-percentile of the respective data. The experimental data are lacking in the gray shaded areas. (**B**) Contour plot comparing the genome-wide distribution of observed compartment signal (PC1) of IMR90 with the one obtained from the simulation. Density is the number of 100-kb chromatin segments in a bin of $0.1 \times 0.1$ square on the plane. Pearson's correlation coefficient is $r = -0.03$. In **A**, a periodic pattern near the diagonal of the contact matrix of simulated chromosome 1 reflects the helical conformation of the chain in telophase. The simulated fine-graded chain of mitotic chromosome takes a helical structure with a $\sim 10$ Mb pitch due to the repulsion among chromatin regions, which disappears from the properly annotated chromosome chains as the chain expands at the entry to interphase, but remains here in the randomly annotated chromosome chain.

**Fig. S9** : **Comparison of the simulated results obtained by doubling the trajectory length with the experimental data.** Length of the simulation trajectories was doubled from 70,000 steps in the ordinary sampling to 140,000 steps. (**A**) Comparison of the experimentally observed (Rao et al 2014 *Cell*, lower left in each panel) and simulated (upper right in each panel) contact frequencies of chromosomes 1, 10, and 19 of IMR90, shown with a 100 kb resolution. The observed and the simulated contact frequencies are normalized to the 95-percentile of the respective data. The experimental data are lacking in the gray shaded areas. (**B**) Contour plot comparing the genome-wide distribution of observed compartment signal (PC1) of IMR90 with the one obtained from the simulation. Density is the number of 100-kb chromatin segments in a bin of $0.1 \times 0.1$ square on the plane. Pearson's correlation coefficient is $r = 0.76$. (**C**) The simulated (red) and observed (black) contact frequency $P(s)$ averaged over the genome for the sequence separation $s$; this simulated $P(s)$ profile is almost same as the one in Fig. 5D in the main text, showing the robustness of the results against the trajectory extension.

**Fig. S10** : **Comparison of the simulated results obtained with the binary annotation of sequence with the experimental data.** Each 100-kb region in the genome sequence was annotated as type-A ($Z_w > 0$) or type-B ( $Z_w \leq 0$). (**A**) Comparison of the experimentally observed (Rao et al 2014 *Cell*, lower left in each panel) and simulated (upper right in each panel) contact frequencies of chromosomes 1, 10, and 19 of IMR90, shown with a 100 kb resolution. The observed and the simulated contact frequencies are normalized to the 95-percentile of the respective data. The experimental data are lacking in the gray shaded areas. (**B**) Contour plot comparing the genome-wide distribution of observed compartment signal (PC1) of IMR90 with the one obtained from the simulation. Density is the number of 100-kb chromatin segments in a bin of $0.1 \times 0.1$ square on the plane. Pearson's correlation coefficient is $r = 0.74$.

**Fig. S11** : **Simulated results with an alternative type-B potential.** (**A**) The functional shapes of the type-A potential (yellow) and an altenative type-B potential (blue) are supoerposed. The alternative type-B potential is defined as $\tilde{U}_{\rm BB}(r) = \tilde{\varepsilon}(1-(r/\sigma_{\rm B})^6)^3$ for $r \leq \sigma_{\rm B}$ and $\tilde{U}_{\rm BB}(r) = 0$ for $r > \sigma_{\rm B}$, with $\tilde{\varepsilon} = 5k_{\rm B}T$. Compared to $U_{\rm BB}$, $\tilde{U}_{\rm BB}$ is more faithful to the coarse-grained potential derived from the fine-grained simulations of type-B chromatin. (**B**) A cross-sectional view of the IMR90 chromosomes in an interphase nucleus simulated with the 100-kb model using the alternative type-B potential. Other forces, parameters and chromatin annotations are the same as those used in the simulations shown in Fig. 4. Mild phase separation of type-A (yellow), type-u (gray) and type-B (blue) chromatin regions are observed. Nucleoli (green) form around rDNA (cyan). (**C**) Comparison of the experimentally observed (Rao *et al* 2014 Cell, lower left) and simulated (upper right) contact frequencies of chromosome 10 of IMR90, shown with a 100 kb resolution. The observed and the simulated contact frequencies are normalized to the 95-percentile of the respective data. The experimental data is lacking in the gray shaded areas. The plaid pattern in the simulated contact map is less evident than the experimental counterpart because of the mild phase separation. (**D**) Contour plot comparing the genome-wide distribution of observed compartment signal (PC1) of IMR90 with the one obtained from the simulation. Density is the number of 100-kb chromatin segments in a bin of $0.1 \times 0.1$ square on the plane. Pearson's correlation coefficient is $r = 0.72$. The mild phase separation still reproduces the genome-wide compartmentalization of the chromosomes observed in the experiment.

**Fig. S12** : **Simulated LADs of chromosomes of IMR90 are compared with the experimental data.** The simulated (black lines) and experimental (Sadaie et al. 2013 *Genes Dev.*, red and pink areas) data are shown. The experimental data are lacking in the gray shaded regions.

**Fig. S13** : **Simulated NADs of chromosomes of IMR90 are compared with the experimental data.** The simulated (black lines) and experimental (Dillinger et al. 2017 *PLoS One*, green and light green areas) data are shown. The experimental data are lacking in the gray shaded regions.

**Fig. S14** : Comparison of the simulated results in the absence of nucleolus with the experimental data. (**A**) Comparison of the experimentally observed (lower left in each panel) and simulated (upper right in each panel) contact frequencies of chromosomes 1, 15, and 19 of IMR90, shown with a 100 kb resolution. The observed and the simulated contact frequencies are normalized to the 95-percentile of the respective data. The experimental data are lacking in the gray shaded areas. (**B**) Contour plot comparing the genome-wide distribution of observed compartment signal (PC1) of IMR90 with the one obtained from the simulation. Density is the number of 100-kb chromatin segments in a bin of $0.1 \times 0.1$ square on the plane. Pearson's correlation coefficient is $r = 0.77$. In **A**, the gray shaded area in chromosome 15 contains the rDNA region. In other regions than the rDNA regions, the simulated data are not changed significantly from the one in Figs. S5 and 6.

**Fig. S15** : **Simulated chromatin movements in interphase IMR90 nucleous.** (**A**) Mean square displacement (MSD) of 100 kb regions of chromatin. (**B**) Time-dependent diffusion constant, $D(t)$. Averaged over type-A (yellow), type-B (blue), and all (gray) chromatin regions.

**Fig. S16** : **Normalized pair-correlation functions between slow ($\mathcal{S}$) and fast ($\mathcal{F}$) chromatin regions.** Correlation between regions separated in distance $d$, $\tilde{g}_{\mathcal{SS}}(d)$ (blue), $\tilde{g}_{\mathcal{FF}}(d)$ (yellow), and $\tilde{g}_{\mathcal{SF}}(d)$ (gray).

**Movie S1** : The simulated expansion process of the IMR90 genome at the entry to the interphase and the subsequent fluctuations in interphase. A 0.4 $\mu$m-width slice of the nucleus is shown. Video frames were sampled at every 1000 steps of the Langevin dynamics simulation and rendered at 20 frames per second, showing approximately 12-minute simulated dynamics in one-second movie motion. The genome is phase-separated into type-A (yellow), type-u (gray), type-B (blue) chromatin regions, and the nucleoli (green). Centromeres (red) locate near the type-B regions, and the rDNA loci (cyan) are buried in the nucleoli.

**Movie S2** : The same simulated dynamics of the IMR90 genome as in Supplementary Movie S1 shown with smoothing along the trajectory. High-frequency vibrations were filtered out by convolving a Gaussian kernel with the trajectory of each chromatin region; denoting the time point of the $k$th video frame by $t_k$, the smoothed trajectory of the $i$th chromatin region $\tilde{\boldsymbol{r}}_i(t)$ was given by the convolution $\tilde{\boldsymbol{r}}_i(t_k) = 1/C_\mathrm{w} \sum_{w=-12}^{12} \exp(-w^2/(2w_0^2))\boldsymbol{r}_i(t_{k+w})$ with the reflective boundary condition. Here, $\boldsymbol{r}_i(t)$ is the raw trajectory of the $i$th region, $C_\mathrm{w} = \sum_{w=-12}^{12} \exp(-w^2/(2w_0^2))$, and $w_0 = 4$.

**Movie S3** : Two-dimensional cross-section view of the simulated dynamics of the IMR90 genome with smoothing along the trajectory. The trajectory is the same as in Supplementary Movie S2. Each 100-kb chromatin region is rendered as a point

**Movie S4** : The simulated dynamics of one of the homologues of chromosome 10 of the IMR90 genome. The trajectory is the same as in Supplementary Movie S1. The first 30 Mb of the chromosome is colored as type-A (yellow), type-u (gray), and type-B (blue) chromatin regions to emphasize how local chromatin domains of different types are spatially separated from each other.

**Movie S5** : The simulated dynamics of one of the homologues of chromosome 10 of the IMR90 genome with smoothing along the trajectory. The smoothed trajectory is the same as in Supplementary Movie S2. The first 30 Mb of the chromosome is colored as type-A (yellow), type-u (gray), and type-B (blue) chromatin regions to emphasize how local chromatin domains of different types are spatially separated from each other.

[6] NCBI databace, https://www.ncbi.nlm.nih.gov/assembly/

[7] The Complete Open-Source and Business Software Platform, https://sourceforge.net/projects/bio-bwa/files/

[8] UCSC Genome Browser, http://hgdownload.cse.ucsc.edu/goldenpath/hg19/

[9] Quinlan AR and Hall IM (2010) *Bioinformatics* 26:84. https://bedtools.readthedocs.io

[10] Samtools, https://www.htslib.org/

[11] Model-based Analysis of ChIP-Seq, https://pypi.org/project/MACS2/

[31] UCSC Genome Browser, http://genome.ucsc.edu/index.html

[32] de Gennes PG (1979) *Scaling Concepts in Polymer Physics*, Cornell Univ Press.

[33] Wu J and Berland KM (2008) *Biophys. J.* 95:2049.

[34] Sen PN (2004) *Concepts Mag. Res. A*, 23A:1.

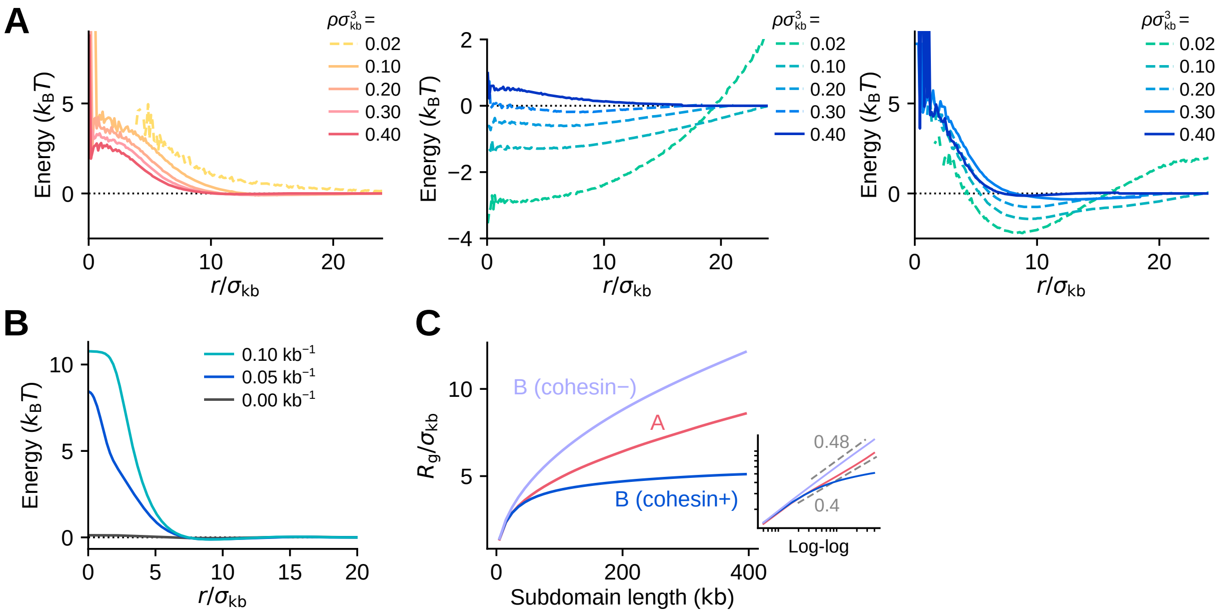

Figure 1: **Characterization of type-A and type-B chromatin in FG simulations.** (**A**) Coarse-grained interaction potentials $U_{\mathrm{CG}}(r)$ before applying smoothing. The coarse-grained potential energy between 100-kb type-A chromatin regions (left), type-B chromatin regions without cohesin (middle) and type-B chromatin regions with cohesin (right) are shown for various chromatin density $\rho$. Solid lines are the potentials derived with the PRISM theory and dashed lines are the potentials of mean force. The plots confirm that the smoothing applied in Fig. 2D-2F in the main text does not alter the essential shape of the potentials. (**B**) Type-B coarse-grained potentials with different cohesin loading constant $k_{\mathrm{on}}/k_{\mathrm{off}} = 0, 0.05, 0.1\,\mathrm{kb}^{-1}$. The potential curve becomes steeper as more cohesin is loaded. (**C**) The simulated radius of gyration of type-A (red), cohesin-loaded type-B (blue) and cohesin-free type-B (pale blue) chromatin as a function of the length of a subdomain. The curve for cohesin-free type-B chromatin does not show a saturating behavior.

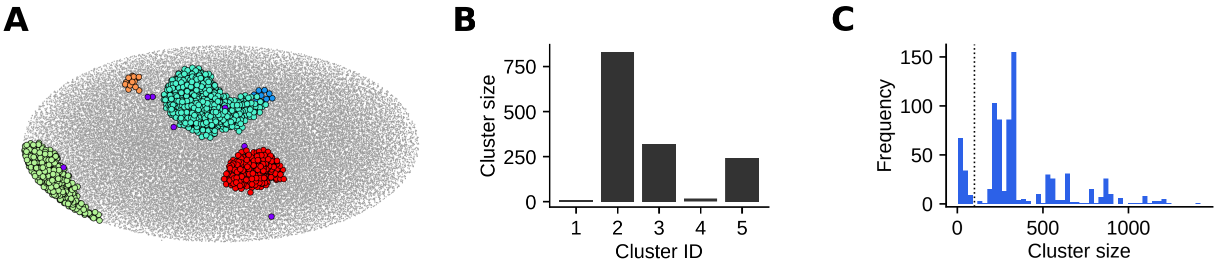

Figure 2: **Simulated nucleoli defined as clusters of nucleolar beads.** (**A**) Clustering of the distributed nucleolar beads in an example IMR90 cell. Simulated nucleolar beads were clustered according to their three-dimensional positions and projected onto the two-dimensional xy plane for visualization. Five clusters (blue, cyan, green, yellow and red) and the DBSCAN outliars (purple) were detected. (**B**) Cluster size of the example shown in **A**. The first and fourth clusters are small, representing the rogue beads separated by thermal fluctuations from nucleoli; these rogue beads were not counted as nucleoli in Fig. 4B of the main text. (**C**) Distribution of the cluster size in 200 simulated IMR90 cells. The dotted line at the cluster size 100 separates the small-cluster outliars.

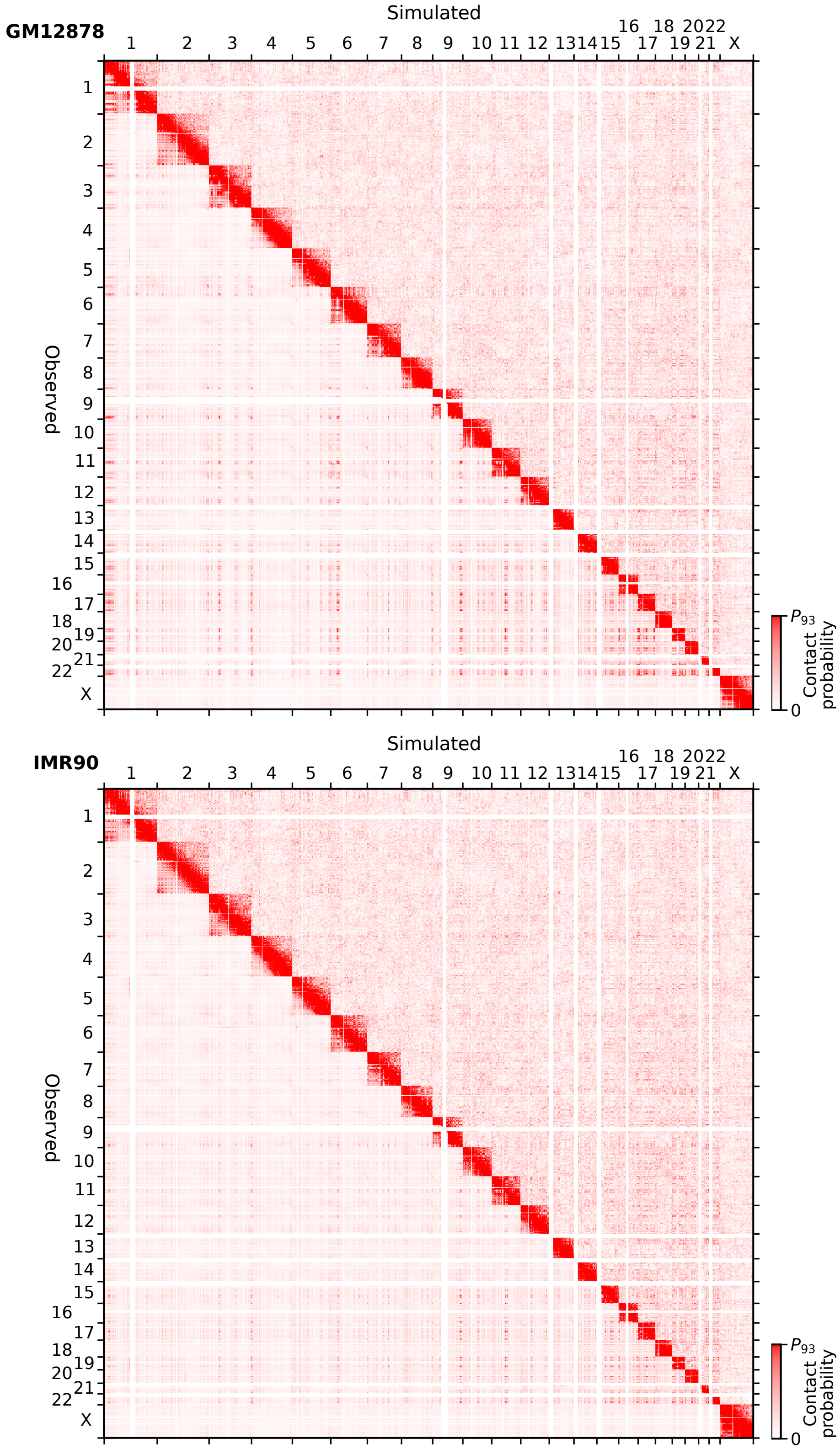

Figure 3: **Comparison of the experimentally observed (Rao et al 2014 *Cell*, lower left in each panel) and simulated (upper right in each panel) genome wide contact frequencies.** Top: GM12878 and Bottom: IMR90. Shown with a 1 Mb resolution. The observed and the simulated contact frequencies are normalized to the 93-percentile of the respective data.

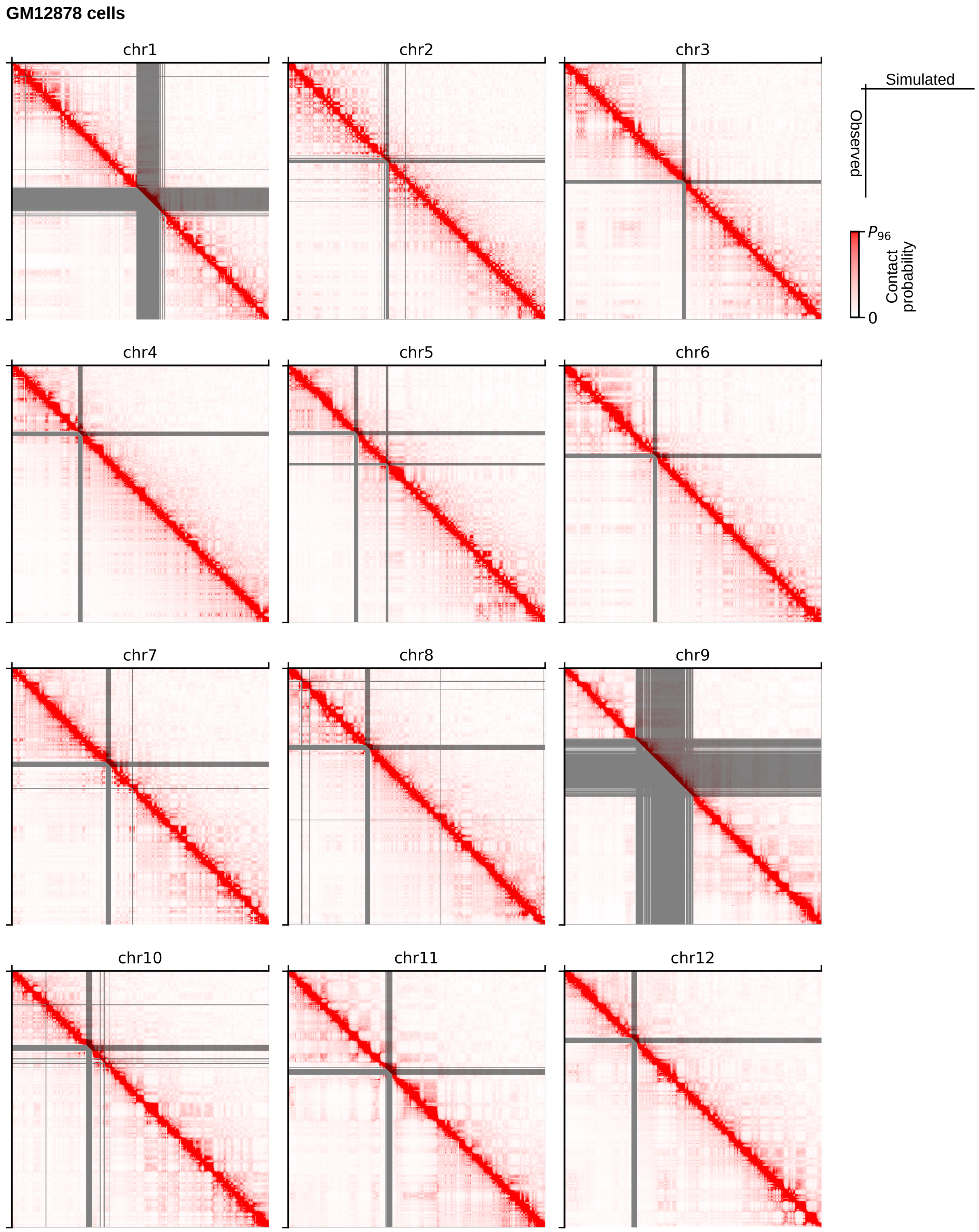

Figure 4: **Comparison of the experimentally observed (Rao et al 2014 *Cell*, lower left in each panel) and simulated (upper right in each panel) contact frequencies of chromosomes 1 to 12 of GM12878.** Shown with a 100 kb resolution. The observed and the simulated contact frequencies are normalized to the 96-percentile of the respective data. The experimental data are lacking in the gray shaded areas.

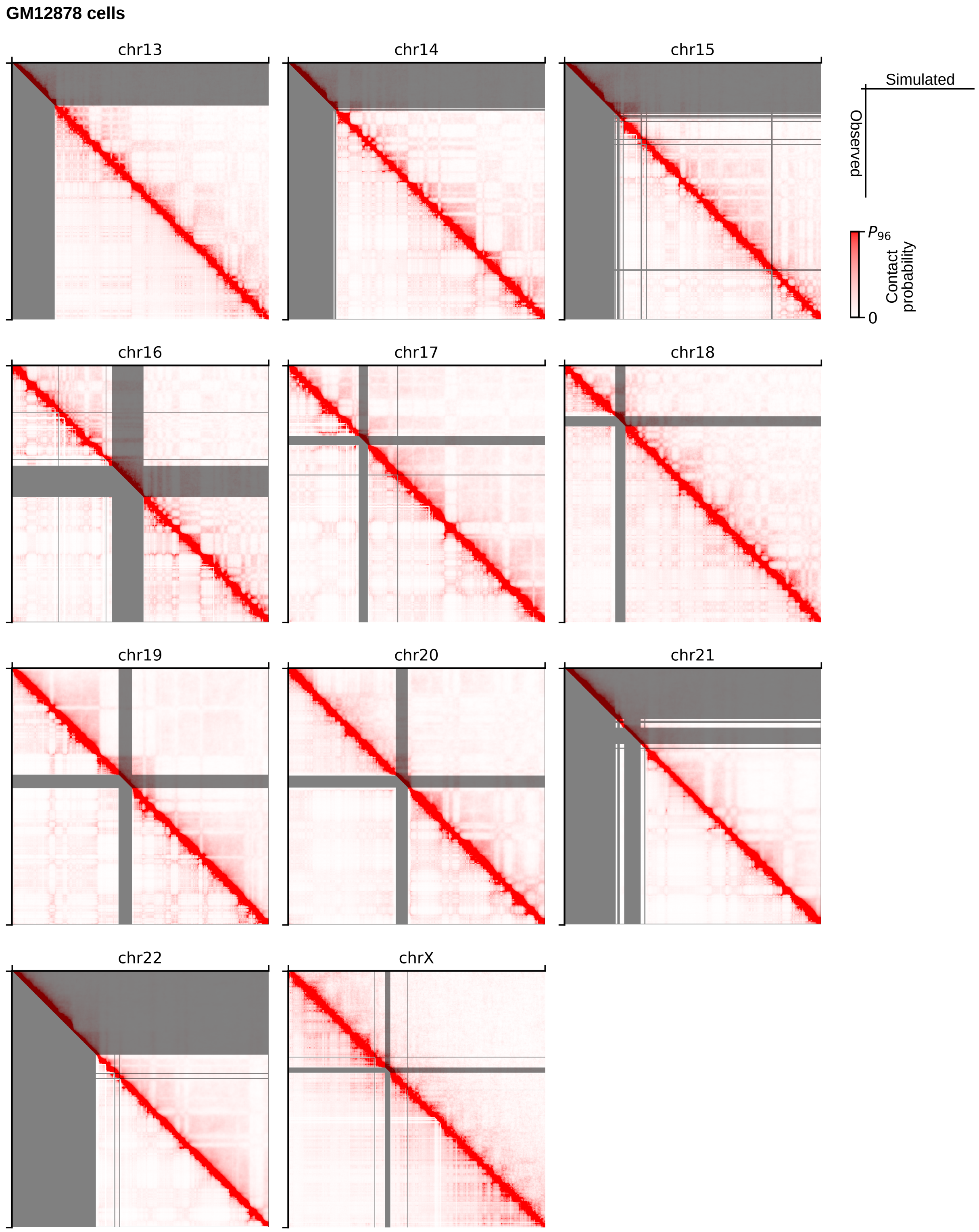

Figure 5: **Comparison of the experimentally observed (Rao et al 2014 *Cell*, lower left in each panel) and simulated (upper right in each panel) contact frequencies of chromosomes 13 to 22 and chromosome X of GM12878.** Data of active and inactive X chromosomes were averaged. Shown with a 100 kb resolution. The observed and the simulated contact frequencies are normalized to the 96-percentile of the respective data. The experimental data are lacking in the gray shaded areas.

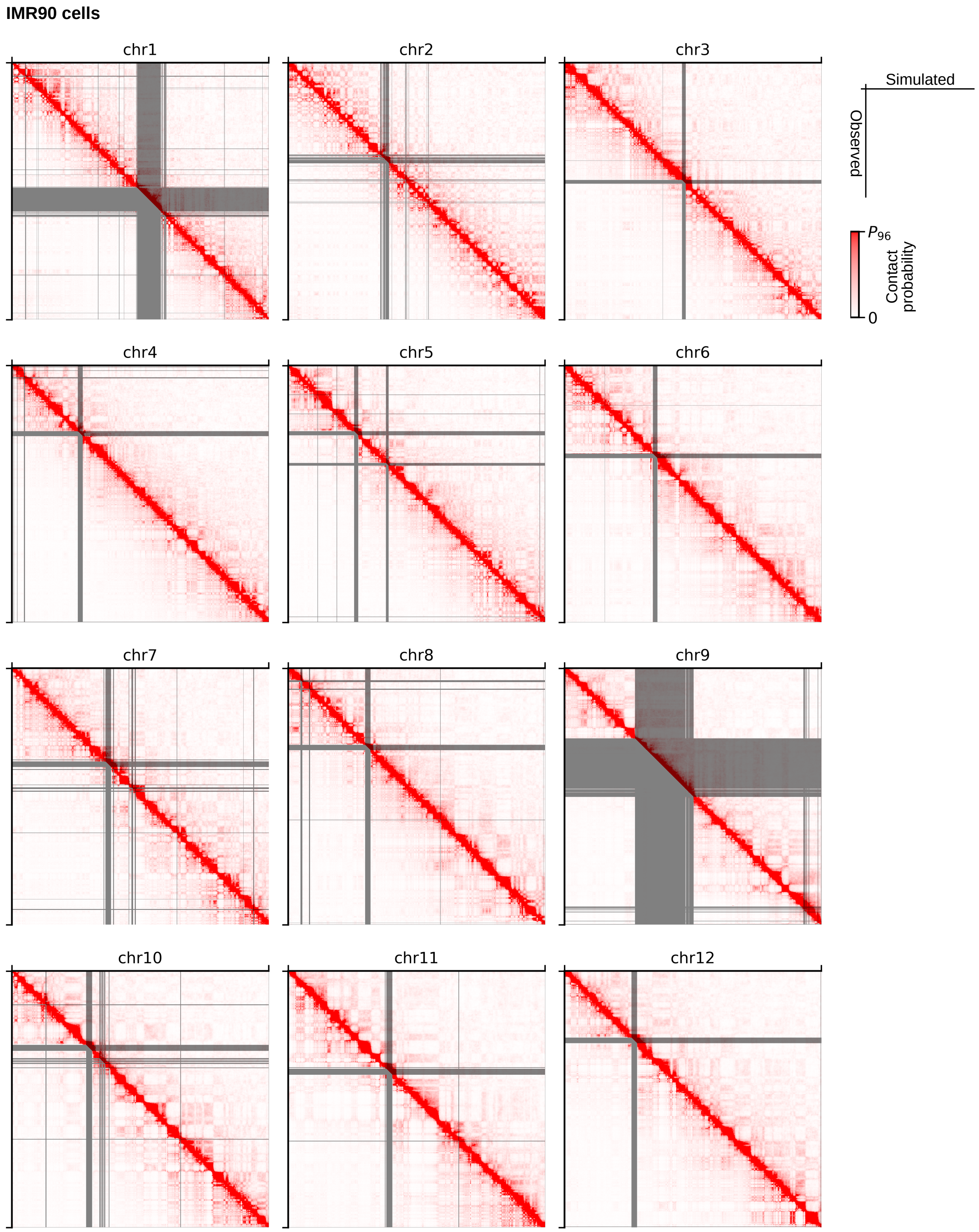

Figure 6: **Comparison of the experimentally observed (Rao et al 2014 *Cell*, lower left in each panel) and simulated (upper right in each panel) contact frequencies of chromosomes 1 to 12 of IMR90.** Shown with a 100 kb resolution. The observed and the simulated contact frequencies are normalized to the 96-percentile of the respective data. The experimental data are lacking in the gray shaded areas.

**IMR90 cells**

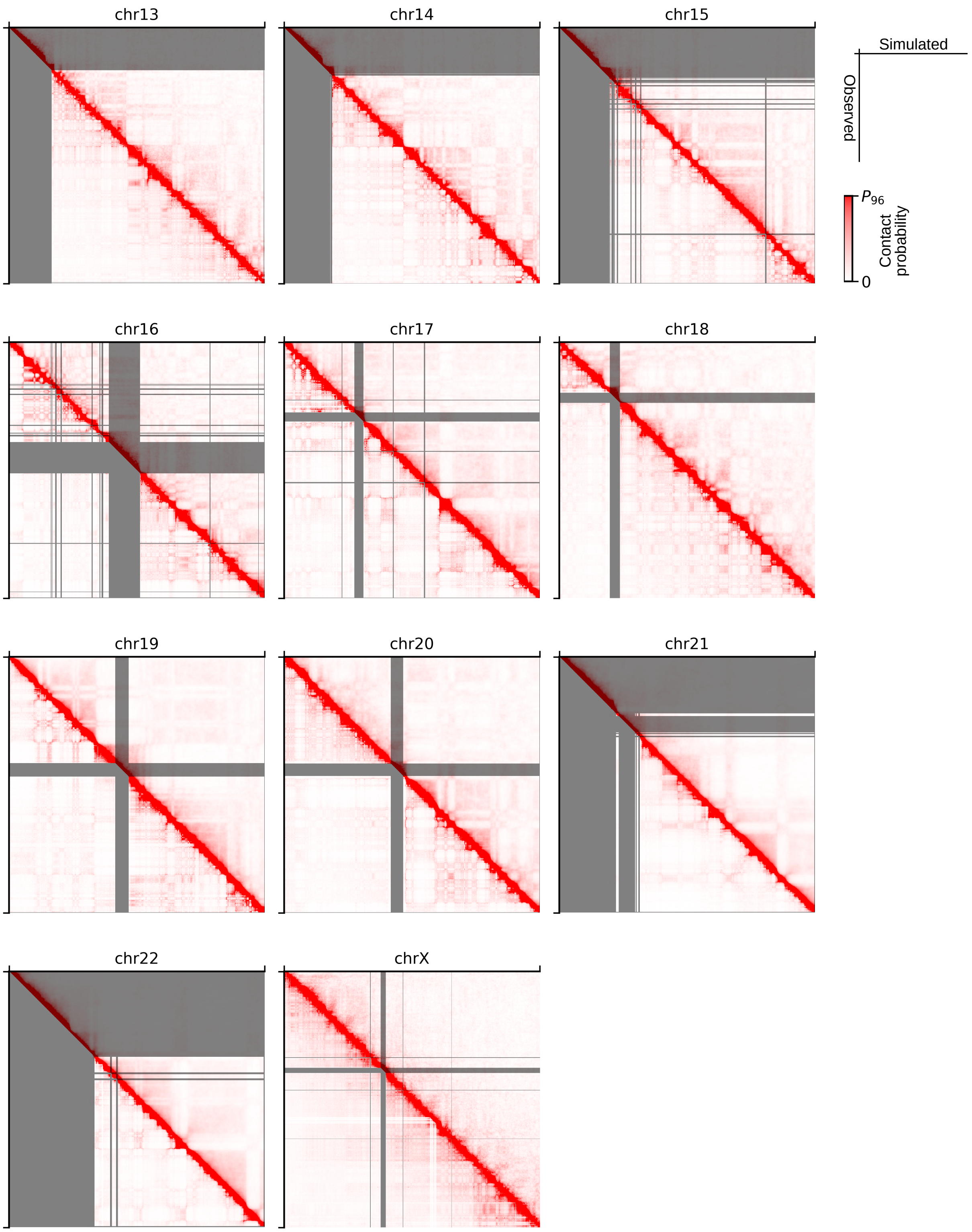

Figure 7: **Comparison of the experimentally observed (Rao et al 2014 *Cell*, lower left in each panel) and simulated (upper right in each panel) contact frequencies of chromosomes 13 to 22 and chromosome X of IMR90.** Data of active and inactive X chromosomes were averaged. Shown with a 100 kb resolution. The observed and the simulated contact frequencies are normalized to the 96-percentile of the respective data. The experimental data are lacking in the gray shaded areas.

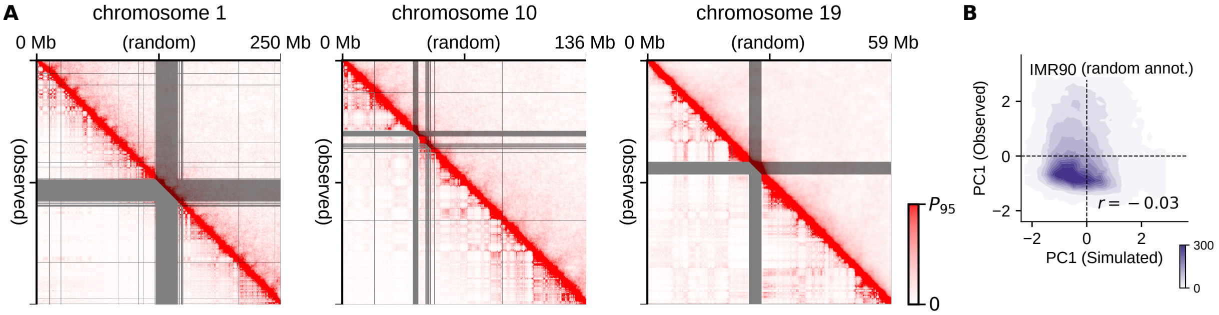

Figure 8: **Comparison of the simulated chromosomes having randomly annotated chromatin types with the experimental data.** (**A**) Comparison of the experimentally observed (Rao et al 2014 *Cell*, lower left in each panel) and simulated (upper right in each panel) contact frequencies of chromosomes 1, 10, and 19 of IMR90, shown with a 100 kb resolution. The observed and the simulated contact frequencies are normalized to the 95-percentile of the respective data. The experimental data are lacking in the gray shaded areas. (**B**) Contour plot comparing the genome-wide distribution of observed compartment signal (PC1) of IMR90 with the one obtained from the simulation. Density is the number of 100-kb chromatin segments in a bin of $0.1 \times 0.1$ square on the plane. Pearson's correlation coefficient is $r = -0.03$. In **A**, a periodic pattern near the diagonal of the contact matrix of simulated chromosome 1 reflects the helical conformation of the chain in telophase. The simulated fine-graded chain of mitotic chromosome takes a helical structure with a $\sim 10$ Mb pitch due to the repulsion among chromatin regions, which disappears from the properly annotated chromosome chains as the chain expands at the entry to interphase, but remains here in the randomly annotated chromosome chain.

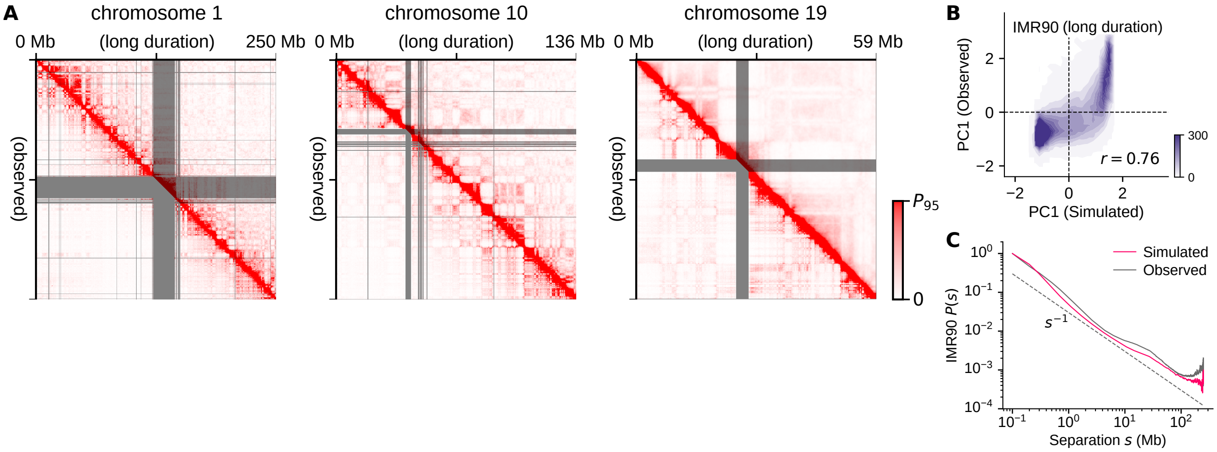

Figure 9: **Comparison of the simulated results obtained by doubling the trajectory length with the experimental data.** Length of the simulation trajectories was doubled from 70,000 steps in the ordinary sampling to 140,000 steps. (**A**) Comparison of the experimentally observed (Rao et al 2014 *Cell*, lower left in each panel) and simulated (upper right in each panel) contact frequencies of chromosomes 1, 10, and 19 of IMR90, shown with a 100 kb resolution. The observed and the simulated contact frequencies are normalized to the 95-percentile of the respective data. The experimental data are lacking in the gray shaded areas. (**B**) Contour plot comparing the genome-wide distribution of observed compartment signal (PC1) of IMR90 with the one obtained from the simulation. Density is the number of 100-kb chromatin segments in a bin of $0.1 \times 0.1$ square on the plane. Pearson's correlation coefficient is $r = 0.76$. (**C**) The simulated (red) and observed (black) contact frequency $P(s)$ averaged over the genome for the sequence separation $s$; this simulated $P(s)$ profile is almost same as the one in Fig. 5D in the main text, showing the robustness of the results against the trajectory extension.

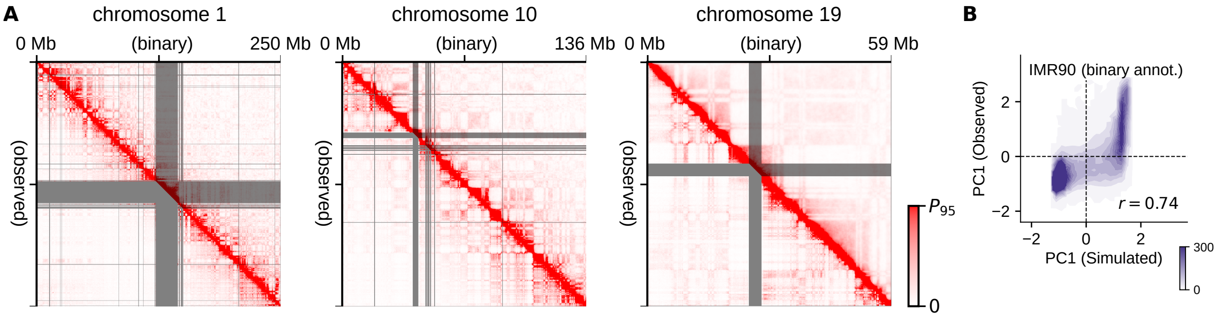

Figure 10: **Comparison of the simulated results obtained with the binary annotation of sequence with the experimental data.** Each 100-kb region in the genome sequence was annotated as type-A ($Z_w > 0$) or type-B ( $Z_w \leq 0$). (**A**) Comparison of the experimentally observed (Rao et al 2014 *Cell*, lower left in each panel) and simulated (upper right in each panel) contact frequencies of chromosomes 1, 10, and 19 of IMR90, shown with a 100 kb resolution. The observed and the simulated contact frequencies are normalized to the 95-percentile of the respective data. The experimental data are lacking in the gray shaded areas. (**B**) Contour plot comparing the genome-wide distribution of observed compartment signal (PC1) of IMR90 with the one obtained from the simulation. Density is the number of 100-kb chromatin segments in a bin of $0.1 \times 0.1$ square on the plane. Pearson's correlation coefficient is $r = 0.74$.

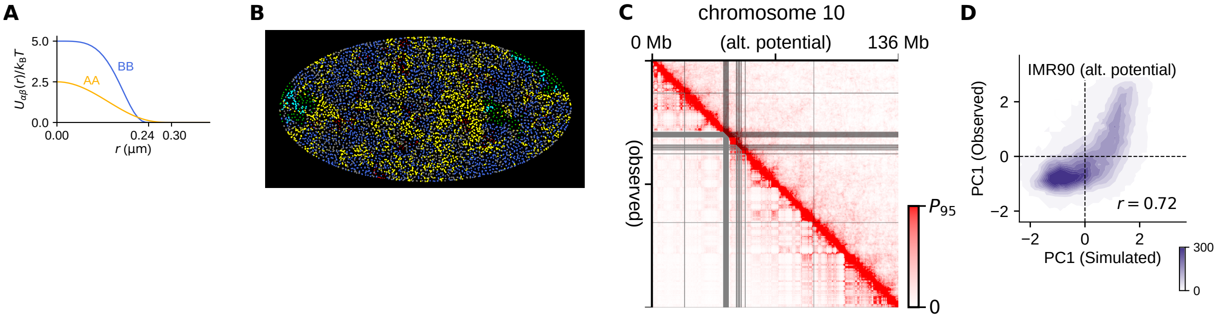

Figure 11: **Simulated results with an alternative type-B potential.** (**A**) The functional shapes of the type-A potential (yellow) and an altenative type-B potential (blue) are supoerposed. The alternative type-B potential is defined as $\tilde{U}_{\rm BB}(r) = \tilde{\varepsilon}(1-(r/\sigma_{\rm B})^6)^3$ for $r \leq \sigma_{\rm B}$ and $\tilde{U}_{\rm BB}(r) = 0$ for $r > \sigma_{\rm B}$, with $\tilde{\varepsilon} = 5k_{\rm B}T$. Compared to $U_{\rm BB}$, $\tilde{U}_{\rm BB}$ is more faithful to the coarse-grained potential derived from the fine-grained simulations of type-B chromatin. (**B**) A cross-sectional view of the IMR90 chromosomes in an interphase nucleus simulated with the 100-kb model using the alternative type-B potential. Other forces, parameters and chromatin annotations are the same as those used in the simulations shown in Fig. 4. Mild phase separation of type-A (yellow), type-u (gray) and type-B (blue) chromatin regions are observed. Nucleoli (green) form around rDNA (cyan). (**C**) Comparison of the experimentally observed (Rao *et al* 2014 Cell, lower left) and simulated (upper right) contact frequencies of chromosome 10 of IMR90, shown with a 100 kb resolution. The observed and the simulated contact frequencies are normalized to the 95-percentile of the respective data. The experimental data is lacking in the gray shaded areas. The plaid pattern in the simulated contact map is less evident than the experimental counterpart because of the mild phase separation. (**D**) Contour plot comparing the genome-wide distribution of observed compartment signal (PC1) of IMR90 with the one obtained from the simulation. Density is the number of 100-kb chromatin segments in a bin of $0.1 \times 0.1$ square on the plane. Pearson's correlation coefficient is $r = 0.72$. The mild phase separation still reproduces the genome-wide compartmentalization of the chromosomes observed in the experiment.

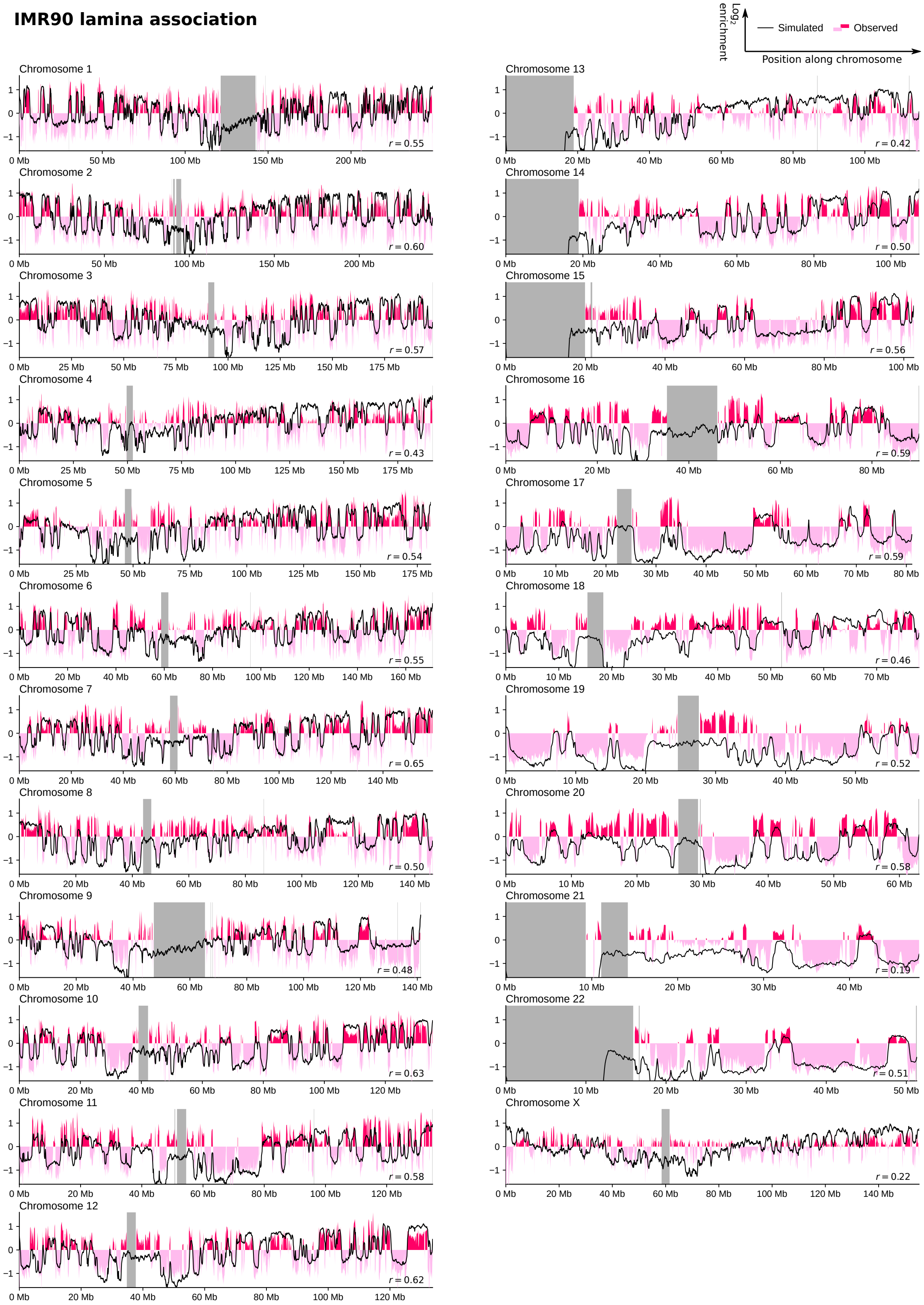

Figure 12: **Simulated LADs of chromosomes of IMR90 are compared with the experimental data.** The simulated (black lines) and experimental (Sadaie et al. 2013 *Genes Dev.*, red and pink areas) data are shown. The experimental data are lacking in the gray shaded regions.

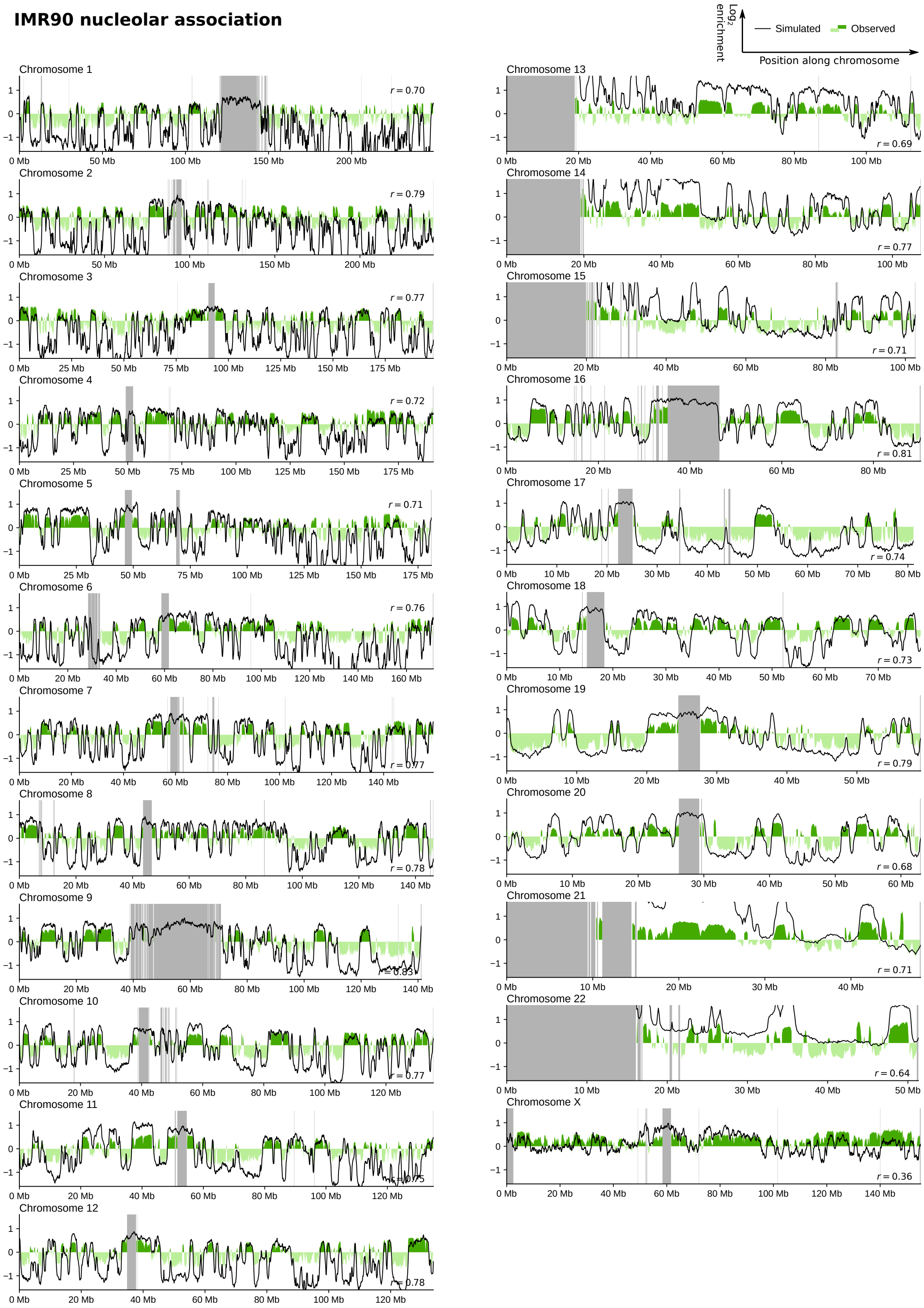

Figure 13: **Simulated NADs of chromosomes of IMR90 are compared with the experimental data.** The simulated (black lines) and experimental (Dillinger et al. 2017 *PLoS One*, green and light green areas) data are shown. The experimental data are lacking in the gray shaded regions.

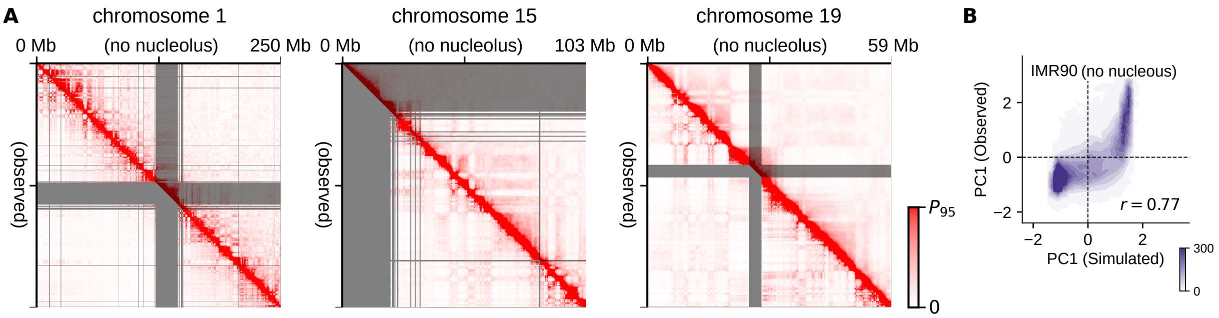

Figure 14: **Comparison of the simulated results in the absence of nucleolus with the experimental data.** (**A**) Comparison of the experimentally observed (Rao et al 2014 *Cell*, lower left in each panel) and simulated (upper right in each panel) contact frequencies of chromosomes 1, 15, and 19 of IMR90, shown with a 100 kb resolution. The observed and the simulated contact frequencies are normalized to the 95-percentile of the respective data. The experimental data are lacking in the gray shaded areas. (**B**) Contour plot comparing the genome-wide distribution of observed compartment signal (PC1) of IMR90 with the one obtained from the simulation. Density is the number of 100-kb chromatin segments in a bin of $0.1 \times 0.1$ square on the plane. Pearson's correlation coefficient is $r = 0.77$. In **A**, the gray shaded area in chromosome 15 contains the rDNA region. In other regions than the rDNA regions, the simulated data are not changed significantly from the one in Figs. S5 and 6.

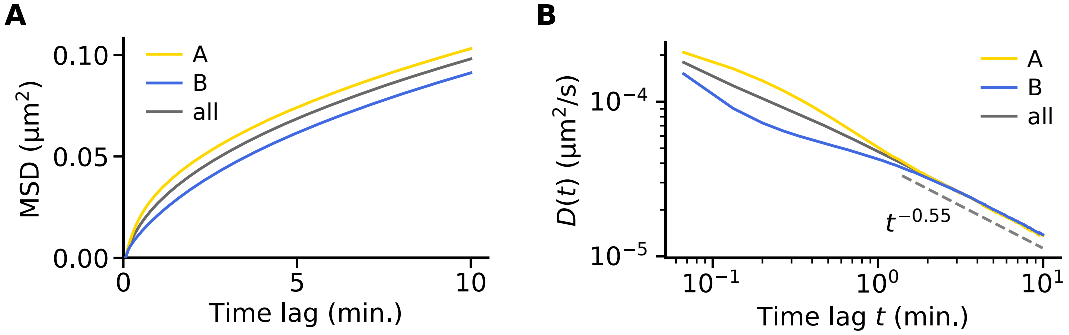

Figure 15: **Simulated chromatin movements in interphase IMR90 nucleous.** (**A**) Mean square displacement (MSD) of 100 kb regions of chromatin. (**B**) Time-dependent diffusion constant, $D(t)$. Averaged over type-A (yellow), type-B (blue), and all (gray) chromatin regions.

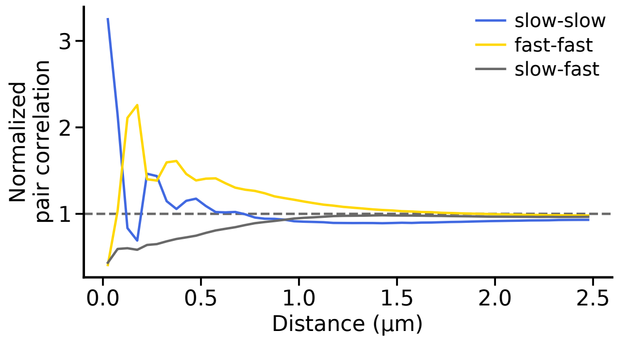

Figure 16: **Normalized pair-correlation functions between slow ($\mathcal{S}$) and fast ($\mathcal{F}$) chro-** [illegible]
